# Response of protein coding genes and microRNAs to temperature changes in four insect species

**DOI:** 10.1101/2024.04.28.591511

**Authors:** Stacey S.K. Tsang, Wenyan Nong, Yichun Xie, Zhe Qu, Ho Yin Yip, Juan Diego Gaitan-Espitia, Amos P. K. Tai, Ying Yeung Yeung, Stephen S. Tobe, William G. Bendena, Jerome H.L. Hui

## Abstract

Insects are the most abundant described living creatures in the world, and they play important roles in our global ecosystem. Climate change affects global biodiversity, and researchers in many fields are striving to better understand the impact of the climate crisis. One such endeavour is the study of temperature-dependent effects on insects. At present, we know little of how climate affects gene expression in insects of different sexes. Here, we took four species of fruit flies of the genus *Drosophila* (*D. melanogaster, D. virilis, D. pseudoobscura,* and *D. erecta*), and subjected the male and female flies of each species to three different temperatures to test their sex-specific gene expression responses. A total of 144 transcriptomic profiles of protein-coding genes and microRNAs were generated. We found that, at the same temperature, there were more male-biased than female-biased protein-coding genes and microRNAs in all four investigated drosophilid species. Interestingly, upon temperature changes, there were more differentially expressed protein-coding genes in females than in males in all four investigated species, while the microRNAs were highly species- and sex-specific. This study provides the first evidence that sex-biased protein-coding gene and microRNA expression responses to temperature change differ between insect species within the same genus, and demonstrates the complexity of sex-specific responses of insects to climate change.

**Highlights:**

- At the same temperature, protein coding gene and microRNA expression showed a greater bias towards males than towards females in all four tested insect species.
- In response to increasing temperature, females of all 4 tested species exhibited more differentially expressed genes than did males, and enrichment analyses showed that they are species-specific.
- Differentially expressed microRNAs did not show a conserved trend between insects upon temperature changes.
- Sex-specific gene and microRNA expression of insects in response to climate change evidently involves a complex adaptation mechanism.

**Graphical Abstract:** 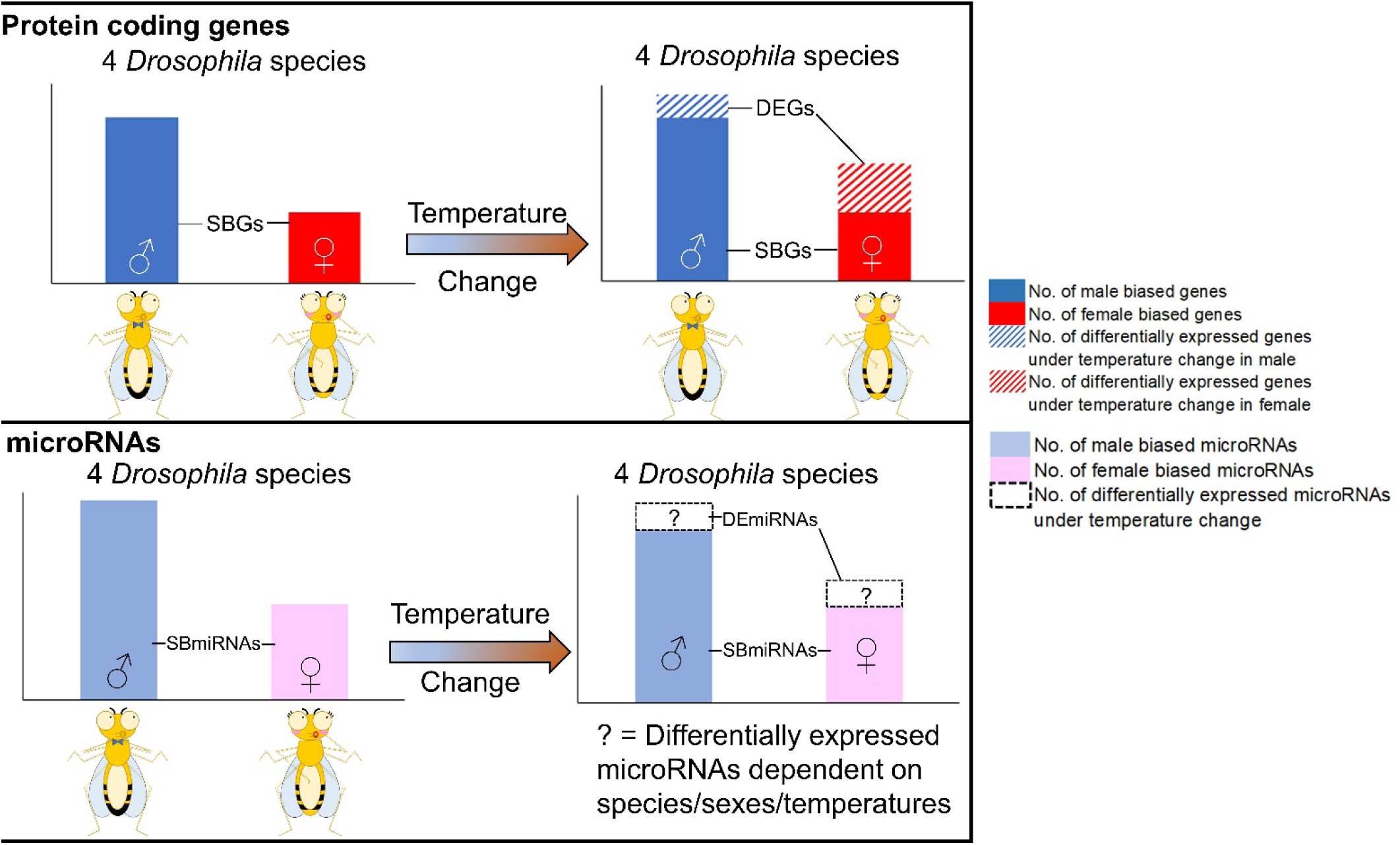

## 1. Introduction

Climate change includes long-term shifts in temperatures and weather patterns, and is one of the biggest threats to the biosphere. Understanding the potential impact on different organisms of phenomena such as heat stress is critical for elucidating the survivorship mechanism and the balance of the ecosystem as a result of climate change [1][2][3][4][5][6].

The arthropods comprise the majority of the described animal species in the world, and insects constitute the largest group of arthropods that dominate the various terrestrial habitats. While insects play important roles in the ecosystem and our society, it has been estimated that more than 60% of the examined insect populations could become extinct during the next century [7]. Temperature directly affects the insect populations. For example, higher temperatures could result in decreases of reproductive success, lethality, and changes in behaviour in certain species [8] [9] [10].

Fruit flies from the various species of the *Drosophila* genus make ideal subjects of study for researchers who wish to understand genetic responses in insects. The genomes of a total number of 32 fruit fly species are available [11]. The genetic responses of different species of drosophilids to environmental changes have been extensively studied in recent decades, and this is an important field of study for improving our understanding of the impact of climate change on insects.

A number of studies have demonstrated that there is a divergence of biological traits, including morphology, physiology, and molecular mechanisms, across and even within species [12] [13]. For instance, it has been shown that populations from different geographical locations can differ, as do sexes of the same species in terms of their impact on protein-coding gene and microRNA expression in different animals e.g. [14][15][16][17][18][19][20]. In terms of response to environmental changes, sexes of the same species also react differently. For example, differences have been reported for male and female marine birds in mercury bioaccumulation and parasitic infections [21], dioecious plants have different levels of herbivore resistance under drought stress between sexes [22], and algae also have sex differences to abiotic stresses [23]. In *Drosophila*, heat tolerance and mitochondrial DNA have been demonstrated to decrease in aging males but not in aging females [24]. A protein-rich diet promoted the expression of transcription factor genes for reproduction in females, but not in males [25]. Furthermore, transcriptional coactivator *Spargel* helps to promote the expression of *stunted* mRNA in female flies and contributes to the male-female difference in body size plasticity.[26]

For noncoding RNAs such as microRNAs, temperature-responsive microRNAs have been proposed to mediate the adaptation in *Drosophila* upon temperature changes. [27][28], even though no evidence has been found for sex-biased microRNAs that preferentially target the sex-biased genes [29]. The relationships between temperature changes, sex-biased protein coding gene expression, and their potential regulation by microRNAs, however, remain largely unexplored. A recent study has suggested that different sexes of insects would respond to temperature changes differently, as documented by the differential protein-coding gene and microRNA expression in butterflies [30].

To further our understanding of the unique adaptations between insect sexes in response to climate change, we utilized *D. melanogaster, D. erecta, D. pseudoobsura* and *D. virilis* and tested the protein coding gene and microRNA repertoires at different temperatures. The four *Drosophila* species were chosen based on their geographical distributions and evolutionary relationships. *D. melanogaster* is a cosmopolitan species, *D. erecta* is native to tropical regions, *D. pseudoobscura* is distributed in both the tropical and temperate zones of western America, while *D. virilis* can be found from tropical to temperate area of eastern Asia [31] and these four species also occupy on key phylogenetic positions in the drosophilid phylogeny [32]. This makes them attractive candidates for exploring the potential regulatory roles played by sex-biased gene expression and for testing whether sex-specific responses to temperature changes are conserved between species.

## 2. Materials and Methods

### 2.1 Temperature treatment

Three of the four *Drosophila* species, viz. *D. erecta* (strain: TSC#14021-0224.01) *D. virilis* (strain: TSC#15010-1051.87) and *D. pseudoobscura* (strain: TSC#14011-0121.94), were purchased from the National Drosophila Species Stock Center. The fourth species, *D. melanogaster* (w1118), was obtained from the Bloomington Drosophila Stock Center. The insects were reared with yeast-cornmeal-agar medium in plastic vials at 25°C and 45-50% relative humidity with light-dark cycle 14hr:10 hr. 6-8 individual female or male virgin flies within 1 day of post-eclosion were placed in a climate box for 24 hours at three different temperatures (18 °C, 25°C, 30 °C), while the relative humidity and light-dark cycle were kept constant. Bearing in mind that different *Drosophila* species have different maximum fatal temperatures, the rationale for choosing these three testing temperatures was: (a) because 25°C is a suitable temperature for maintaining a live culture of most of the *Drosophila* species; (b) because 18°C is known to have a biological impact on *Drosophila* culture such as extended generation time [33], and therefore made an appropriate minimum testing temperature; and (c) because 30 °C, as the highest temperature that all 4 species could still survive after 24 hours, made an appropriate maximum testing temperature.

### 2.2 mRNA and small RNA transcriptomes sequencing

After the treatment, mRNA and small RNA were extracted with mirVana™ miRNA Isolation Kit in accordance with the total RNA extraction protocol. Experiments were repeated on different days to obtain three biological replicates for male and female flies of the four different species at different temperatures. The quantity and quality of the extracted total RNA was checked with Nanodrop spectrophotometer (Thermo Scientific), 1% agarose gel electrophoresis, and Agilent 2100 Bioanalyser (Agilent RNA 6000 Nano Kit). Transcriptome libraries were constructed and sequenced by BGI and Novogene on the Illumina platform. Details of the sequencing data and their accession numbers are provided in Tables S1 and S2.

### 2.3 mRNA and small RNA transcriptomic analysis

Raw reads were trimmed and filtered using Trimmomatic (version 0.39, with parameters ‘ILLUMINACLIP: TruSeq3-PE.fa:2:30:10 SLIDING WINDOW:4:5 LEADING:5 TRAILING:5 MINLEN:25’) [34] and Kraken (version 2.0.8) [35] for contamination removal. The processed reads of each sample were aligned to the reference genome downloaded from Flybase using HISAT2 [36]. Gene read count matrix tables were generated by StringTie [37] and submitted to iDEP.96 [38] for further analysis.

The analyses performed in iDEP.96 were carried out as previously described [38]. The gene ID were first recognized by choosing the target species as respective *Drosophila* species with default parameters (0.5 counts per million (CPM) in 1 library and PCA as EdgeR: log2(CPM+c)). The differentially expressed genes were then further divided into sex-biased genes (i.e. 18°C Male vs 18°C Female, 25°C Male vs 25°C Female and 30°C Male vs 30°C Female) and differentially expressed genes under different temperatures (i.e. 30°C Male vs 18°C Male, 30°C Male vs 25°C Male, 25°C Male vs 18°C Male, 30°C Female vs 18°C Female, 30°C Female vs 25°C Female, 25°C Female vs 18°C Female) were identified with DESeq2 calculation method with default false discovery rate (FDR)<0.1, adjusted p-value < 0.1, |log_2_(fold-change)| > 1. The differentially expressed genes were then further subjected to Gene ontology (GO) biological process terms, cell component terms, molecular functions and KEGG pathway enrichment analysis with adjusted p-value <0.01. The orthology of the differentially expressed genes identified in different species was performed by OrthoFinder (version 2.5.4) [39]. The sex-biased genes identified under 18°C, 25°C and 30°C of each species were compared to reveal those genes displaying biased expression at all 3 tested temperatures, while the differentially expressed genes under different temperature regimes of the same sex were also compared to identify the genes commonly regulated at increased temperatures. Enrichment results were visualized with bubble plots (http://www.bioinformatics.com.cn/srplot) and compared across different species with Venn diagrams (http://bioinformatics.psb.ugent.be/webtools/Venn).

Small RNA sequencing reads were analyzed in accordance with a previously-described method [40]. Adapters were trimmed using cut adapt, and quality control was performed using fastqc. Qualified reads within 18 and 27 bp were used to generate the matrix count table using the ‘quantifier.pl’ module of the mirDeep2 package [41] with ‘-g 0’ to allow 0 mismatches when mapping reads to precursors. The generated count matrix was used for expression analysis in edgeR [42]. Target prediction of microRNAs was carried out with DEGs showing a reverse trend of expression (i.e. targets of up-regulated microRNAs were used to predict for targets within the list of down-regulated genes) with miranda and RNAhybrid. For *D. melanogaster*, prediction results from TargetScanFly were also included for comparison purposes.

### 2.4 Real-time PCR validation

For validating the expression of gene predicted to be targeted by Dme-miR-34, three biological replicates of RNAs from both sexes of *D. melanogaster* after 24 hours treatment under 18°C and 30°C were extracted, using the method mentioned in Section 2.2. These 12 RNA samples were further reverse-transcribed into cDNAs using iScript™ gDNA Clear cDNA Synthesis Kit, in accordance with the manufacturer’s protocol. The Real-time PCR experiment was conducted using the CFX96 Touch Real-Time PCR Detection System (BioRad), with the following conditions: denaturation at 95°C for 3min followed by 40 cycles of (95°C/10 s, 60°C/10 s, and 72°C/15 s). PCRs were run with iTaq Universal SYBR Green Supermix (BioRad). Two technical replicates were performed for each biological replicate. The expression of each gene was normalized to the expression of the housekeeping gene (18S ribosomal RNA gene), and the relative fold changes of each target gene were calculated using the ΔΔCt method. Information about the primers is shown in Table S3.

## 3. Results

### 3.1 Sex-biased protein coding genes in four drosophilid species (same temperature)

Males and females from four species of drosophilid (*Drosophila melanogaster, D. erecta, D. pseudoobscura* and *D. virilis*) were exposed to different temperatures (18°C, 25°C and 30°C), and a total of 144 transcriptomes (72 mRNA and 72 small RNA) were obtained (Fig 1; Tables S1-S2).

**Fig 1.**
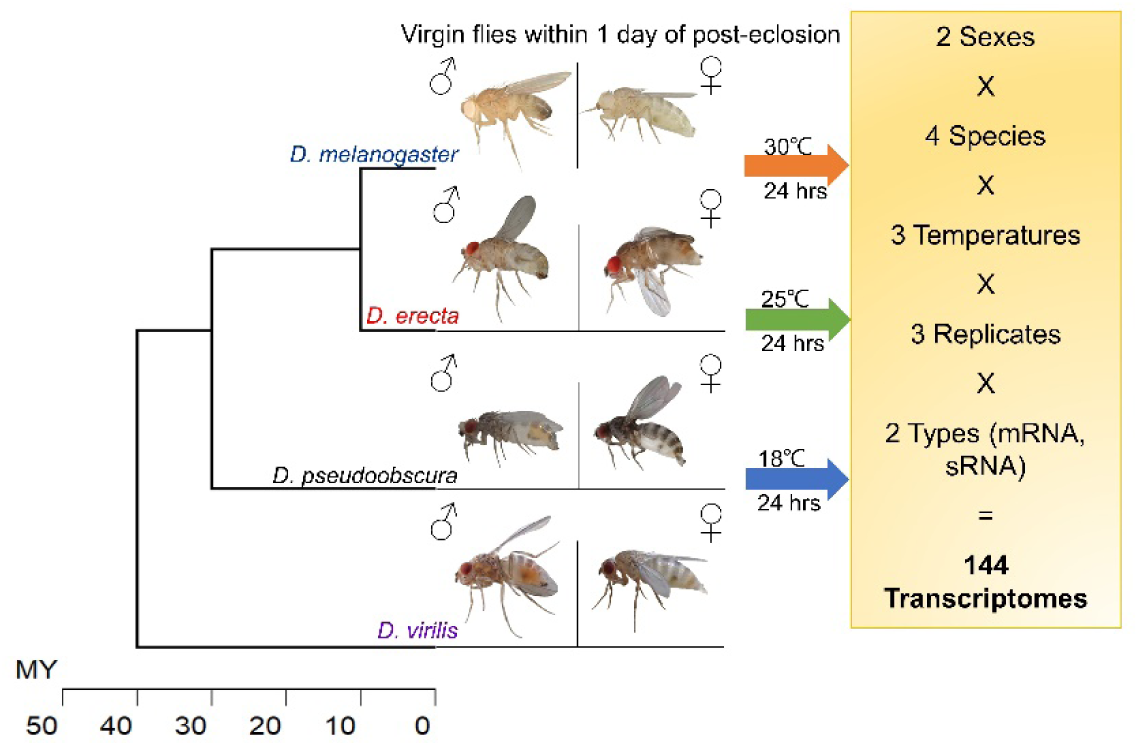
Schematic diagram showing the experimental design of this study

By comparing the male and female flies’ mRNA transcriptomic profiles at the same temperature (e.g. 18°C male vs 18°C female of the same species), we first aimed to reveal the number of sex-biased genes (SBGs) in the four species of drosophilid. Here, the SBGs were defined as genes with significantly different expression levels (|log_2_(fold-change) | > 1) between the two sexes. We found that there were more male-biased genes than female-biased genes in the four investigated species at all four temperatures tested (Fig. 2), which agrees with the findings of previous studies. See [43][44][45] and Tables S4-S5.

**Fig 2.**
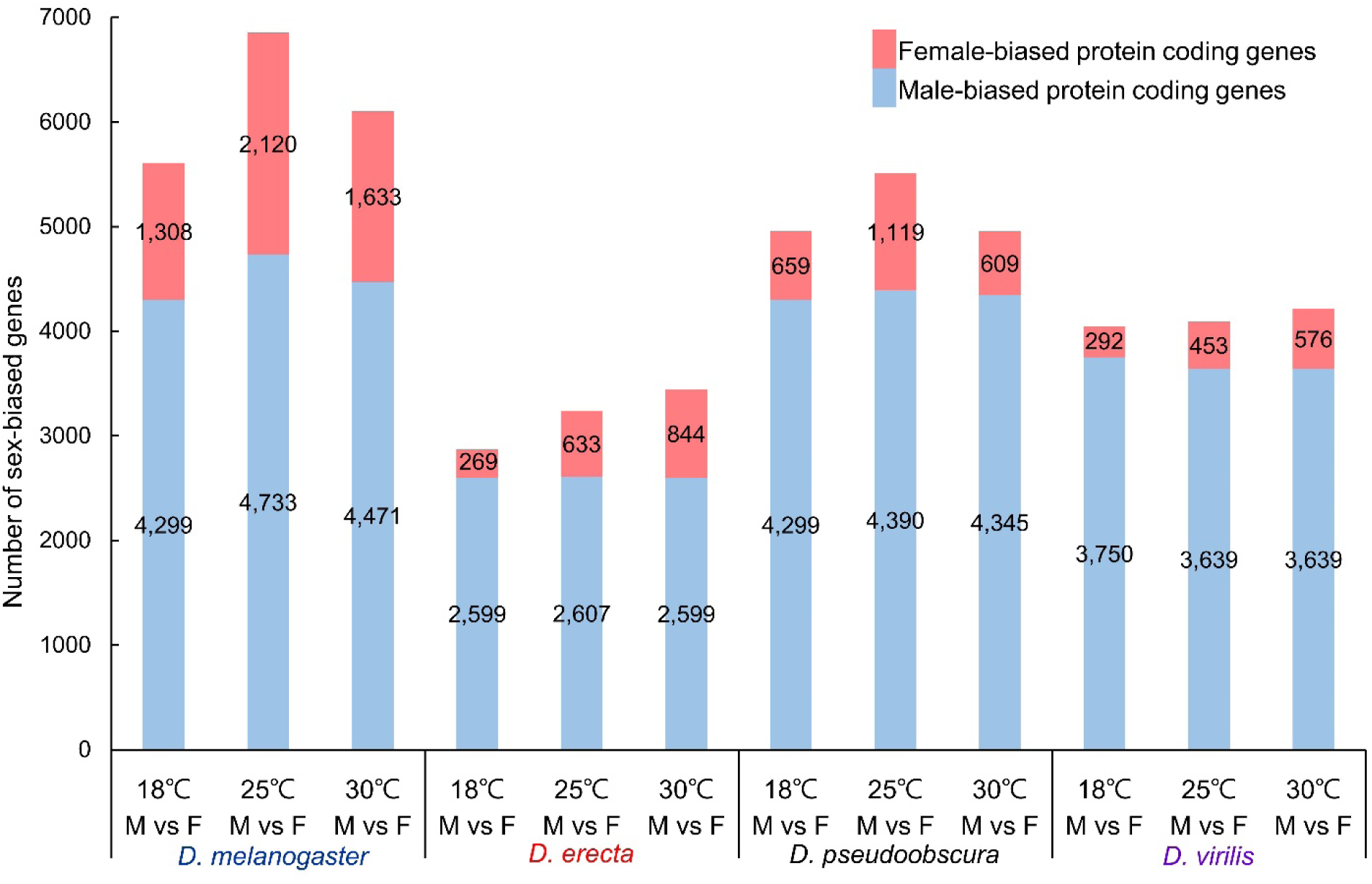
Bar chart showing the number of sex-biased genes at different temperatures. There were significantly more male biased genes than female biased genes of all 4 species at 3 tested temperatures.

To determine whether the above SBGs were conserved between the four drosophilids species, we further carried out a comparison between the ortholog groups of the SBGs, gene ontology (GO) (Figs. S2-4) and KEGG pathway (Fig. S1) enrichment analyses. We established that there were conserved gene pathways (such as the ribosome biogenesis gene pathway) in all four drosophilid species (Fig. 3; Fig. S5). In the case of the SBGs found to be conserved in all 4 species at all three tested temperatures, these genes were mainly related to cilium movement. In terms of molecular function, “nucleic acid binding”, “helicase activity” and “structural constituent of cuticle” were commonly enriched in 4 species but at different temperatures (Table S6A). For instance, the structural constituent of cuticle is male-biased under 25°C in *D. melanogaster*, but the same molecular function was female-biased under 18 °C and 25°C in *D. erecta*, at 30 °C in *D. pseudoobscura*, and at either 18 °C or 30 °C in *D. virilis* (Table S6C, Fig. S4).

**Fig 3.**
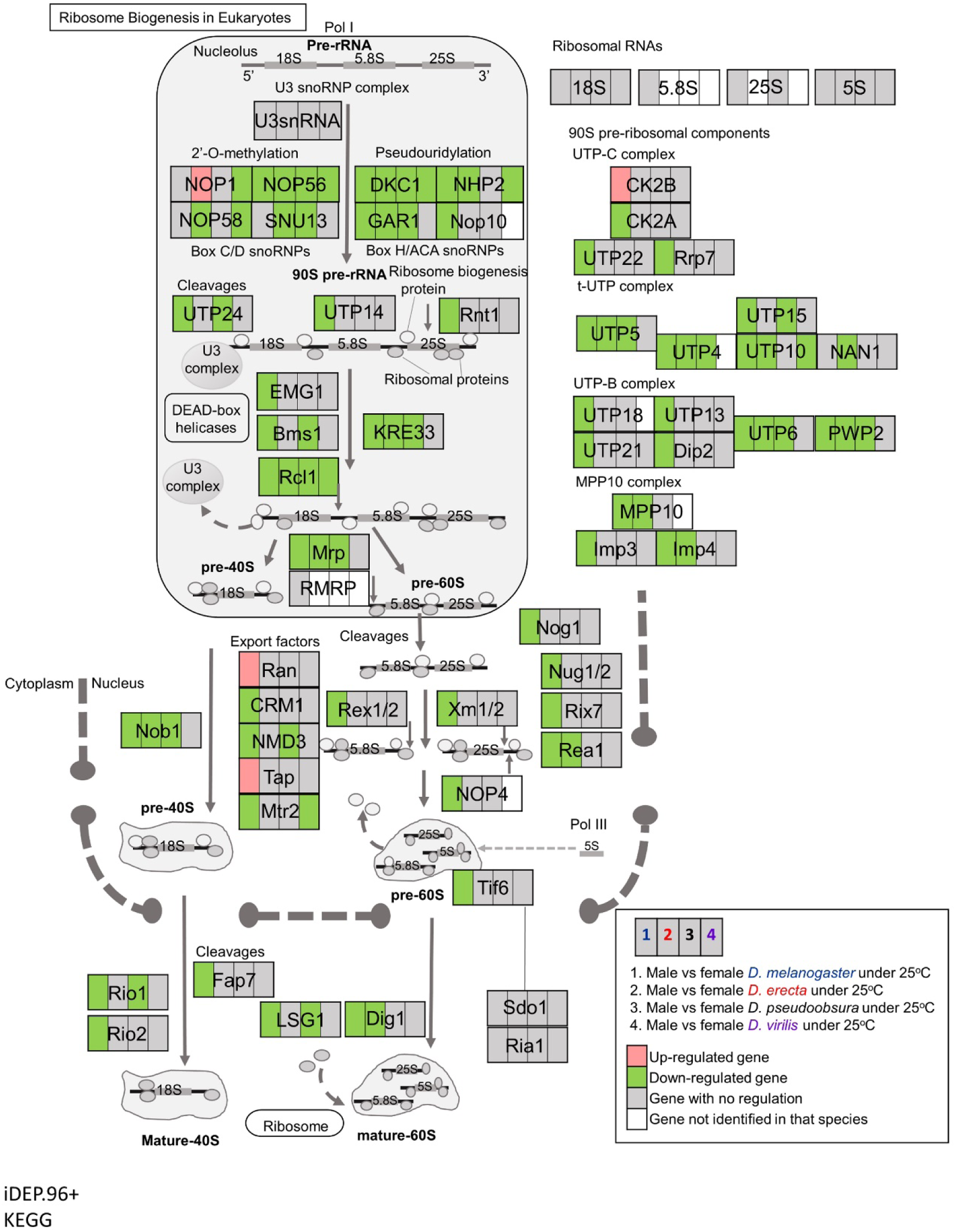
Conserved pathway (Ribosome biogenesis in eukaryotes) enriched by sex-biased genes of 4 species at 25°C. Please refer to Fig. S5 for further details of the regulation of each gene under different conditions.

Several well-known genes related to sex differentiation, including doublesex (dsx), transformer (tra), and Sex-lethal (Sxl), were also investigated. Our findings are summarized in Table S7. We found that their regulation in response to temperature changes were species-specific and the expression were not constantly sex-biased in all tested temperatures. For example, transformer (tra) was sex-biased in *D. pseudoobscura* and *D. virilis* at 18 °C.

### 3.2 Sex-biased microRNAs in four drosophilid species (same temperature)

To reveal the sex-biased microRNAs in the different species under the various test conditions, we also compared the small RNA transcriptomic profiles of male and female flies at the same temperature (e.g. 18°C male vs 18°C female of the same species). The sex-biased microRNAs here were defined as microRNAs, with an expression level significantly different (|log_2_(fold-change) | > 1) between the two sexes. We found that there were more male-biased microRNAs than female-biased microRNAs in the investigated drosophilid species, similar to our findings in the protein coding genes (Fig 4; Tables S8-S10A)

**Fig 4.**
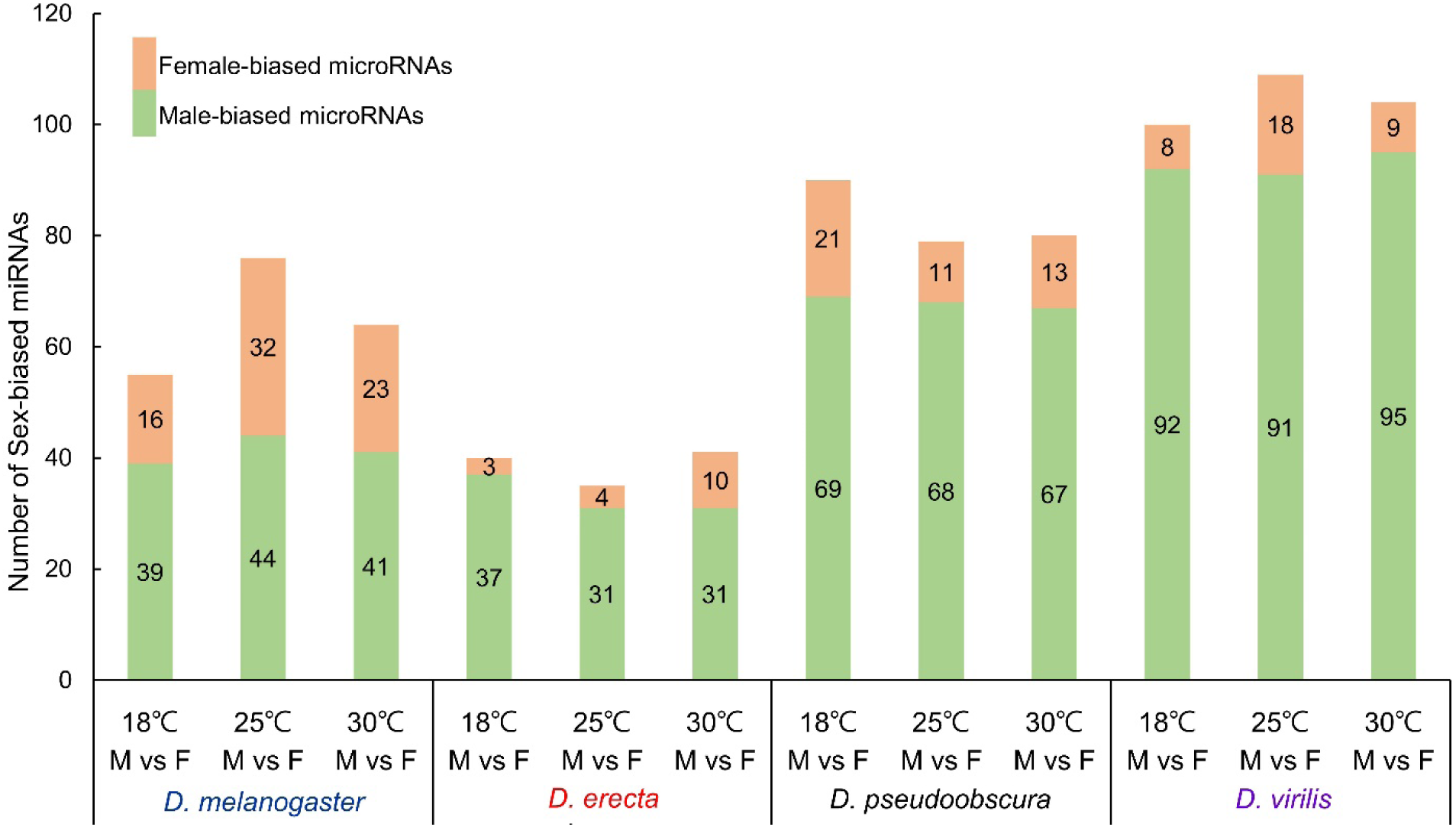
Bar chart showing the number of sex-biased microRNAs at different temperatures. There were significantly more male biased microRNAs than female biased microRNAs of all 4 species at 3 tested temperatures.

Further comparison of microRNAs with SBGs revealed that *Dyskerin Pseudouridine Synthase 1* (*DKC1*) was commonly targeted by sex-biased microRNAs in the four drosophilid species, including miR-975 targeted *DKC1* in both *D. melanogaster* and *D. erecta* (Table 1; Table S11). In addition, there were 6 sex-biased microRNAs that were predicted to target different genes in the ribosome biogenesis pathway, including miR-964, miR-973, miR-975, miR-983, miR-992, and miR-2498 (Table 2). Cumulatively, these analyses suggested a conserved sex-biased microRNA gene regulatory network established in the ancestor of these drosophilids ∼40 million years ago.

### 3.3 Sex-biased protein coding genes in four drosophilid species (different temperatures)

To explore how different drosophilid species and their sexes respond to temperature changes, we then compared the mRNA transcriptomes of insects of the same sex and species across different temperature treatments (e.g. 18°C vs 30°C in males). Unexpectedly, we discovered that there were more differentially expressed genes (DEGs) in female flies than in male flies during temperature changes (Fig. 5, Figs. S6-7 and Table S12). GO and KEGG pathway enrichment analyses further supported that different responsive genes were utilized during temperature changes between sexes (Table S13 and Figs. S8-11). For instance, the neuroactive ligand receptor interaction pathway was found to be enriched in all four species, but in different degrees between sexes (Fig. 6; Fig. S12). In sum, this trend of gene response reacting to the environment, with more DEGs in female flies than in male flies, remained true for all four investigated species, implying a potential adaptation mechanism established in the *Drosophila* ancestor.

**Fig 5.**
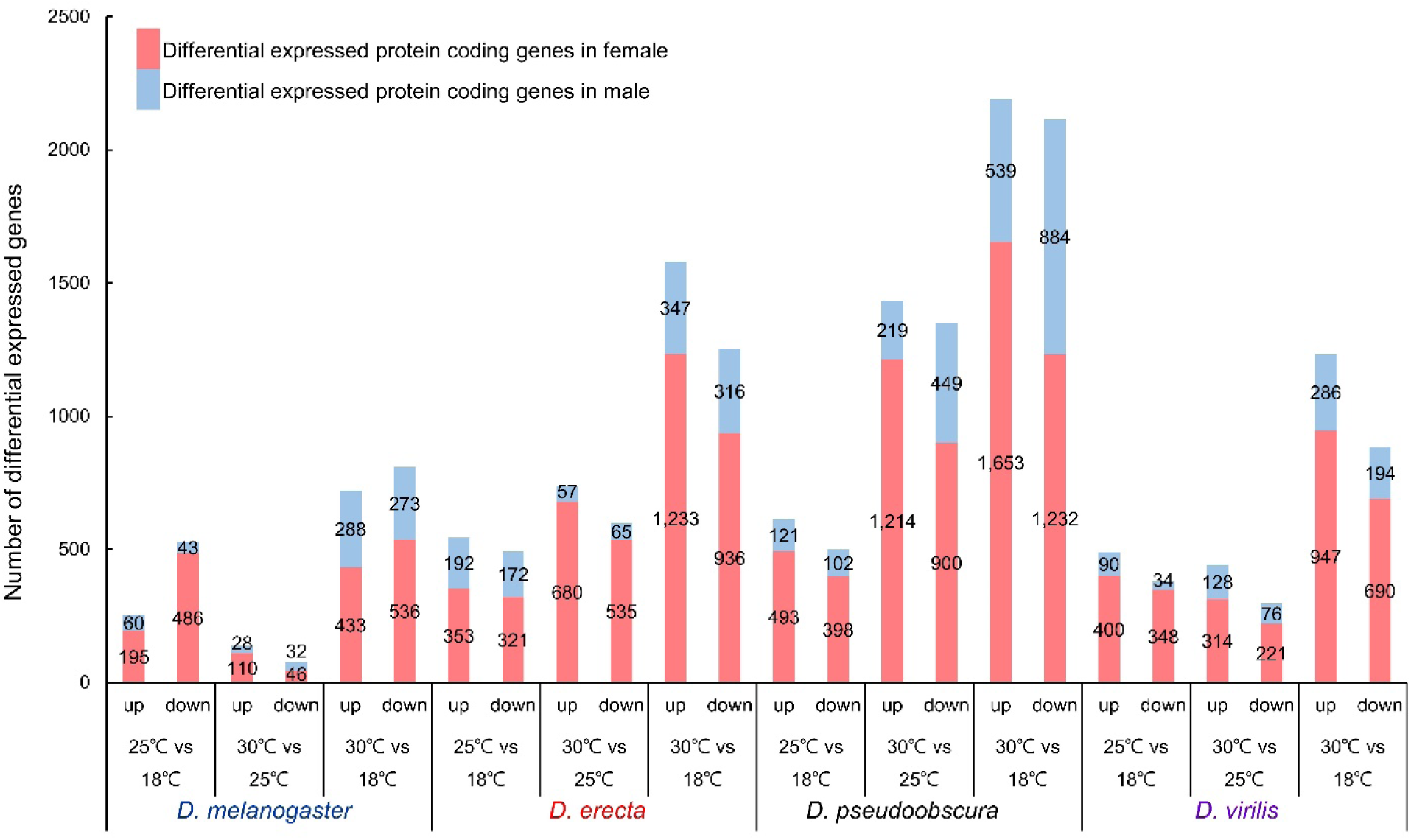
Bar chart showing a comparison of the number of differentially expressed genes at different temperatures within the same sex. The number of differentially expressed genes in females were significantly more than those in males in all 4 species at all temperatures tested.

**Fig 6.**
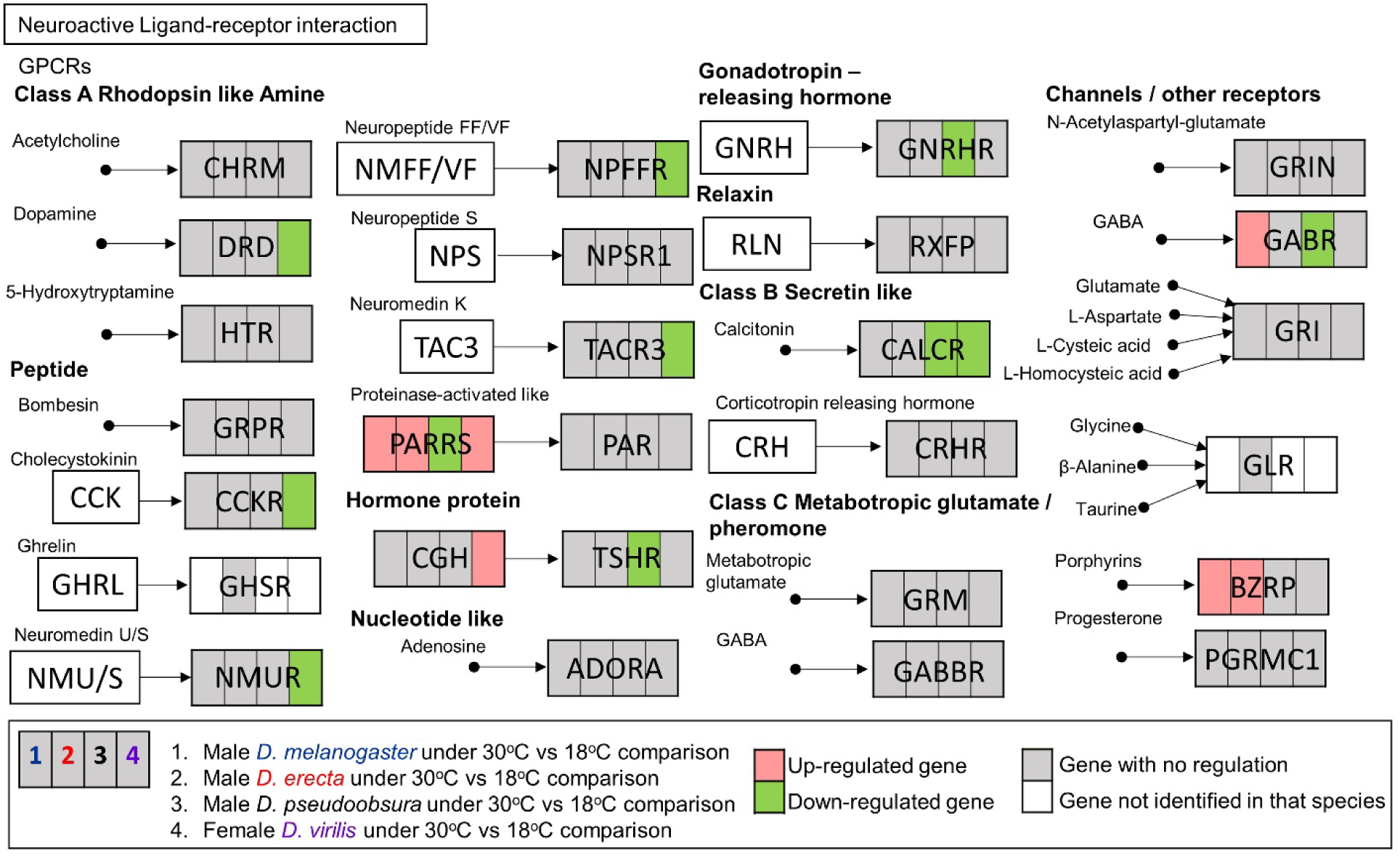
A comparison of the conserved pathway (Neuroactive Ligand-receptor interaction) enriched by differentially expressed genes of the 4 species at 30 °C vs 18 °C. Please refer to Fig. S12 for further details of the regulation of each gene under different conditions.

### 3.4 Sex-biased microRNAs in four drosophilid species (different temperatures)

The comparison of differentially expressed microRNAs across temperatures nevertheless did not show a common trend, such as was shown in the case of protein-coding genes in response to temperature changes (i.e. more DEGs in female flies than in male flies); and the number of differentially expressed microRNAs varied between different scenarios and across species (Fig. 7; Figs. S13-14).

**Fig 7.**
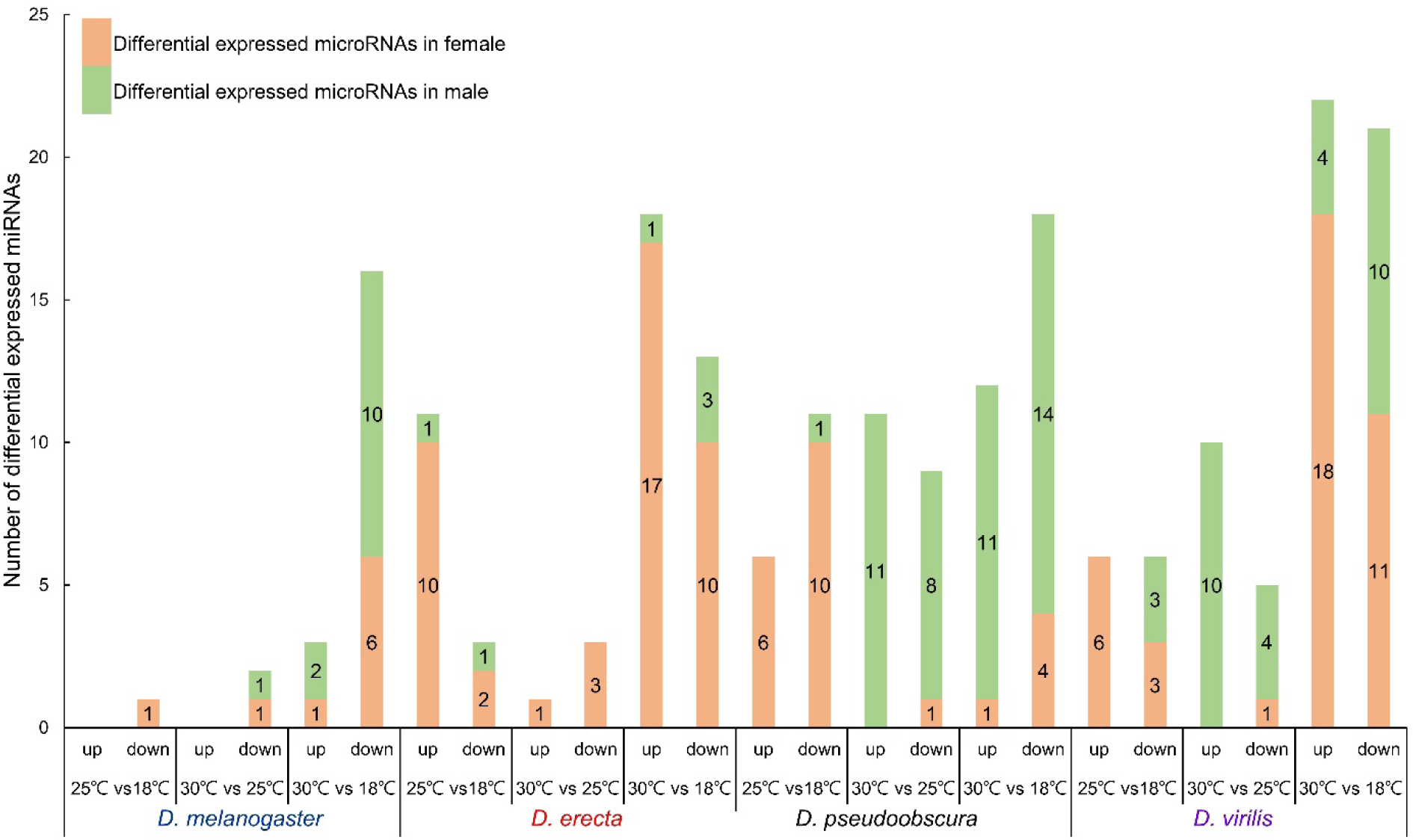
Bar chart showing a comparison of the number of differentially expressed microRNAs at different temperatures within the same sex.

We found that most of the microRNAs responding to temperature changes were sex-specific (Figs. S15-18 and Table S14-S17). For instance, 24 and 4 microRNAs were differentially regulated during temperature changes in males and females of *D. pseudoobscura* respectively, and only 1 of these microRNAs was shared between sexes.

The comparison of the differentially expressed microRNAs upon temperature changes across species found that 7 common microRNAs (miR-278, miR-282, miR-124, miR-308, miR-311, and both members in the miR-277/34 cluster) were identified in 3 species, but their trends of expression were different (Tables S10B and S18). For instance, at 30°C, miR-277-5p was downregulated in both D. melanogaster and D. virilis, while miR-277-3p was upregulated in D. erecta (Fig 8). These data suggested that microRNAs respond to temperature changes differently to protein coding genes during insect evolution and adaptation to the environment. Furthermore, the gene CG10462 that was being targeted by miR-34-3p, showing both sex-biased and differential expression upon temperature treatment, was selected for RT-PCR validation. As shown in Fig. 9a, CG10462 has a female biased expression at 30°C but not at 18°C, and is also down-regulated in females between 18°C and 30°C. These findings all agreed with the expression trends as determined in the transcriptomic analyses (Fig. 9b).

**Fig 8.**
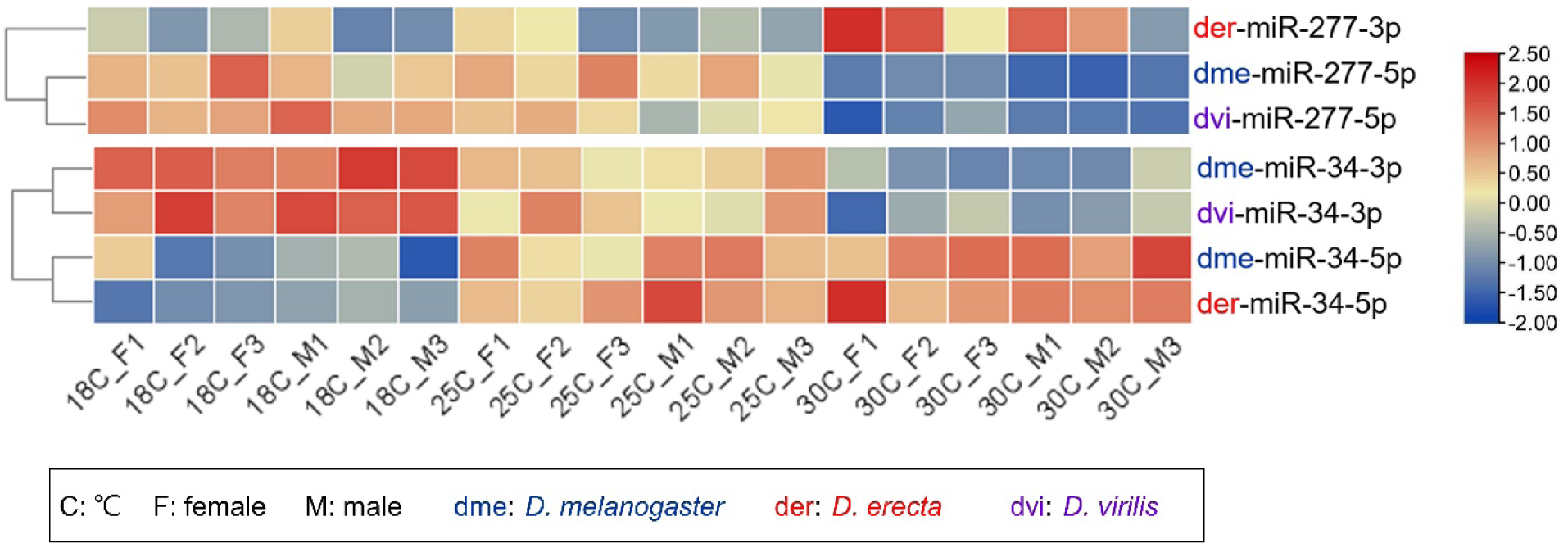
Heatmap showing the differential expression of the miR-277 and miR-34 clusters, which show the same trend of differential expression in both males and females among the 3 species tested. The arms of microRNAs showing differential expression varies between species. Please refer to Fig. S19 for the heatmaps showing the expression of miR-277 and miR-34 clusters of all 4 drosophila species.

**Fig 9.**
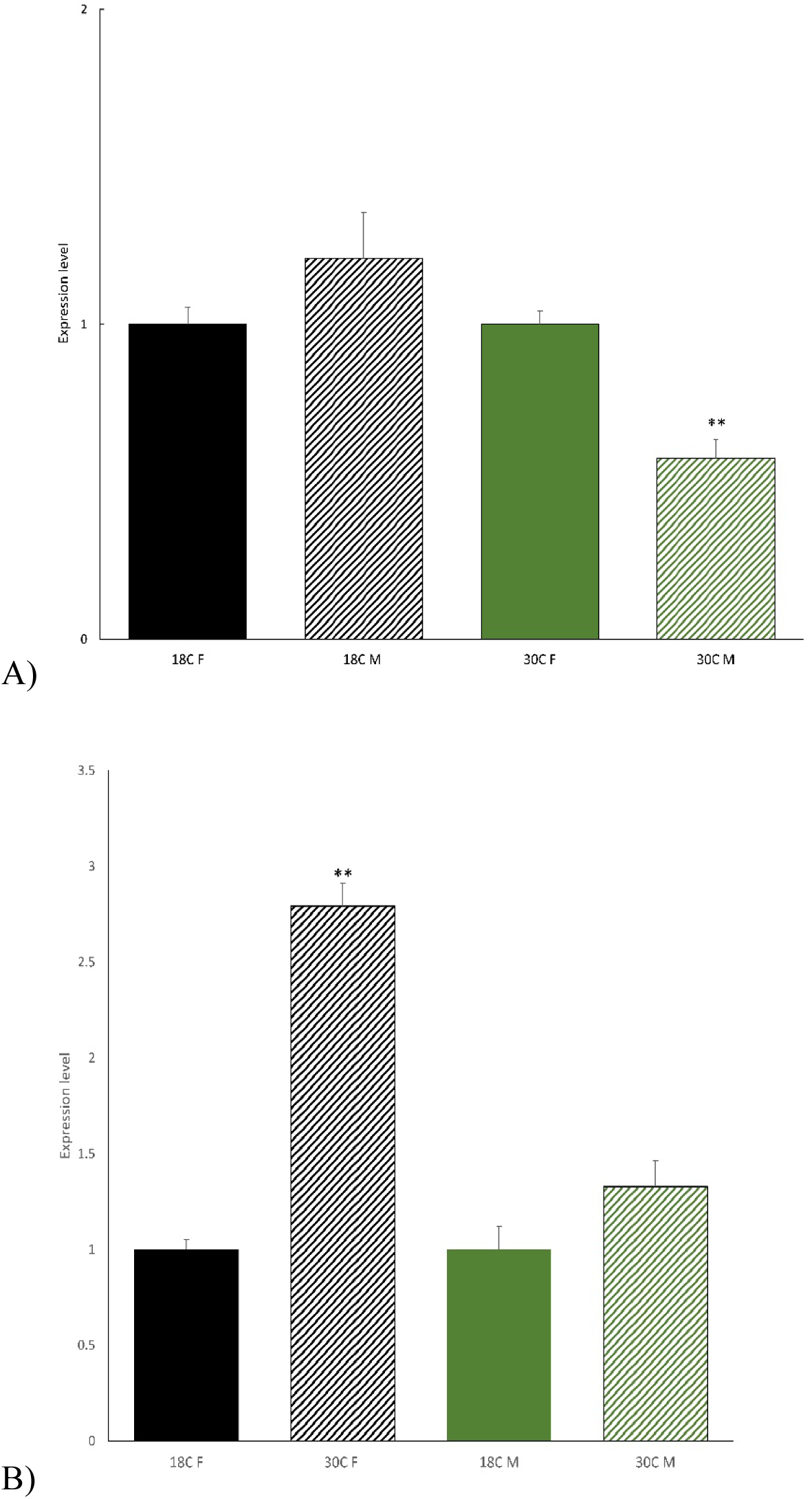
Expression level of predicted target gene CG10462 of miR-34-3p in males and females of *D. melanogaster* under 18 °C and 30 °C treatments. Error bars indicate the standard error of three independent biological replicates. **P ≤ 0.01 based on Student’s t-test. Solid bars are the controls of the striped bars in corresponding colours. a) Asterisks indicate significant differences in gene expression between 30 °C and 18 °C among individuals of the same sex; b) Asterisks indicate a significant gene expression difference between males and females at the same temperature.

## 4. Discussion

Male sex-biased protein coding gene and microRNA expression has long been known to occur in *Drosophila* (e.g.[43][44][46][47][48][49][50][51][52][53][54][55][56][57]), but to date only a relatively small number of studies have explored how diet, drought and chemicals exerted different effects between sexes of *D. melanogaster* (e.g.[12][58][59][60][25][61]). Whether different sexes of insects, in this case drosophilids, would respond to temperature changes in a conserved or lineage-specific manner in terms of their protein-coding gene and microRNA expression, remains unexplored. Here, we have advanced our understanding of this area in several respects.

On the one hand, we discovered that more female-biased protein-coding genes were differentially expressed during temperature changes, and this trend was observed in all four species of drosophilids we investigated. This does not only highlight the differences between sexes of drosophilids in response to temperature changes, but also suggests that this was a system established in the last common ancestor of drosophilids, which responded to temperature changes for their adaptation ∼40 million years ago. Few studies to date have investigated the response between sexes to environmental cues in insects, or have made comparisons across species. In a recent study, it was found that there were conserved sex-biased genes across different species of butterflies (at the same temperature), and more female-biased protein-coding genes were differentially expressed when temperature increased in the common yellow butterfly *Eurema hecabe* [30]. Our current understanding of this topic remains in its infancy.

On the other hand, for post-transcriptional regulator microRNAs, conflicting views on whether there are similar or different selection forces on them in comparison to protein coding genes have been proposed during the evolution of humans and in other animals (e.g.[62][63][64] [65]). As we did not observe a conserved trend for sex-biased microRNAs during temperature changes having more differential expression in our investigated species and conditions, in contrast to the situations of protein coding genes identified above, we have now added that microRNAs evolved along a different route to protein coding genes in species and sexes from the perspective of environmental response during drosophilid evolution. Similarly, as our current understanding of this topic is also in its infancy, the findings of this study open a new avenue to further testing the protein coding genes and microRNAs in insects in response to climate change, such as whether the situations revealed in the drosophilids here are specifically confined to these insects, evolutionary divergence time, geographical distributions, environmental conditions, and/or sex system.

## 5. Conclusion

Our study has demonstrated that sex is a factor which must be included in investigating environmental stress and adaptions. The data sets provided macroscopic and fundamental information regarding thermal stimulated sex-biased expression for further studies. Previous reports on sex-specific microRNAs expression at different temperatures have indicated that microRNA regulation is not the sole mechanism to be taken into account in considering the sex-specific differential expression of protein coding genes. However, this study provided hints for reaching a better understanding of the sex-biased regulation mechanism and discovering the evolutionary trajectory of this sex-specific control.

## Acknowledgements

This study was supported by grants from the Hong Kong Research Grant Council General Research Fund (No. 14100420), the Collaborative Research Fund (No. C4015-20EF), the CUHK Group Research Scheme (No. 3110154), the CUHK Strategic Seed funding for Collaborative Research Scheme (No. 3133356), and a CUHK Direct Grant (No. 4053547). S.S.K.T. was supported by postgraduate studentships provided by The Chinese University of Hong Kong.

## Data availability

The raw reads generated in this study have been deposited to the NCBI database under the BioProject accession number PRJNA974874.

https://www.ncbi.nlm.nih.gov/bioproject/?term=PRJNA974874

## Supplementary Figure

**Fig. S1.**
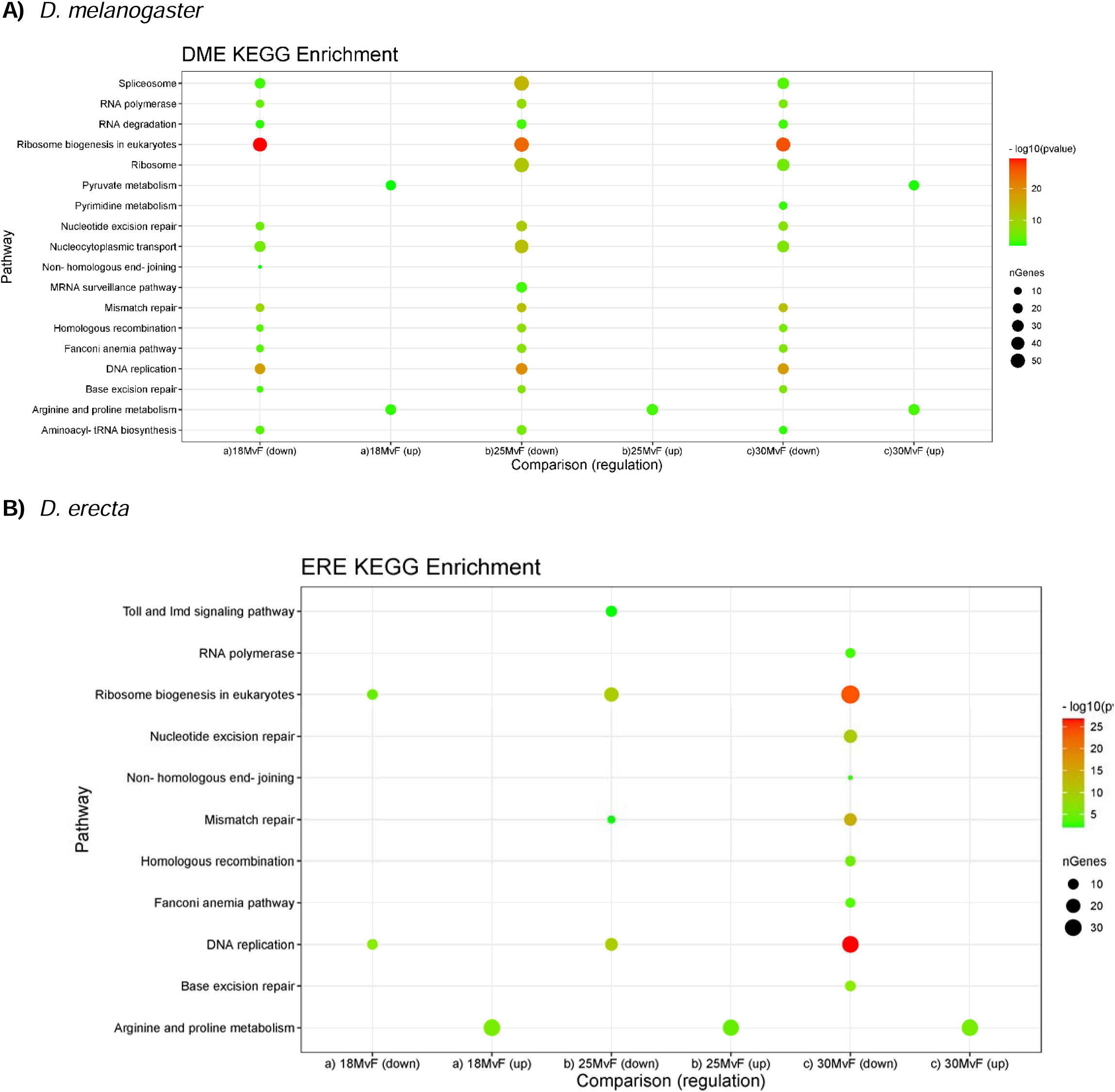
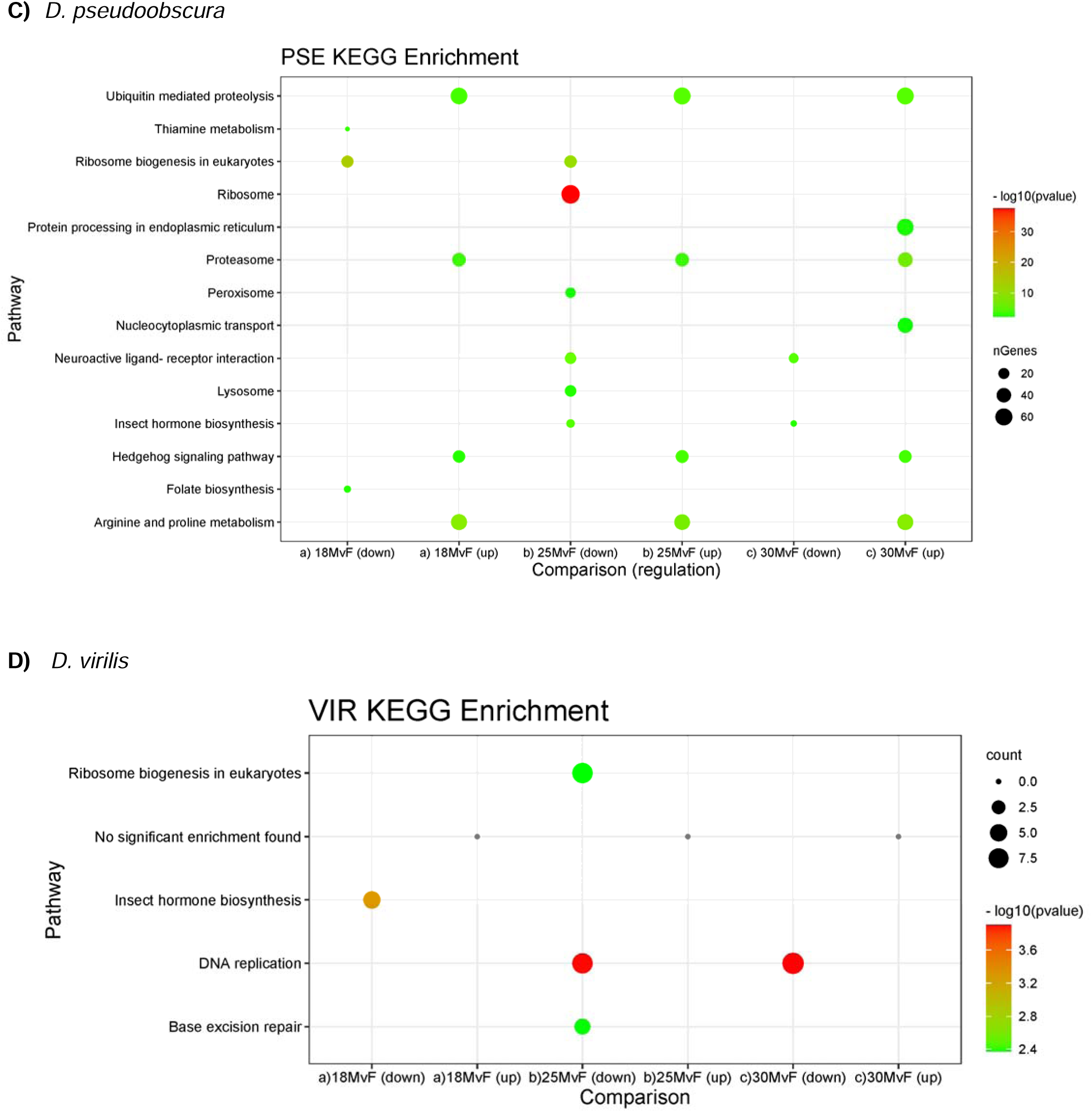
KEGG Pathway enrichment for Sex-biased Genes of 4 *Drosophila* species

**Fig. S2.**
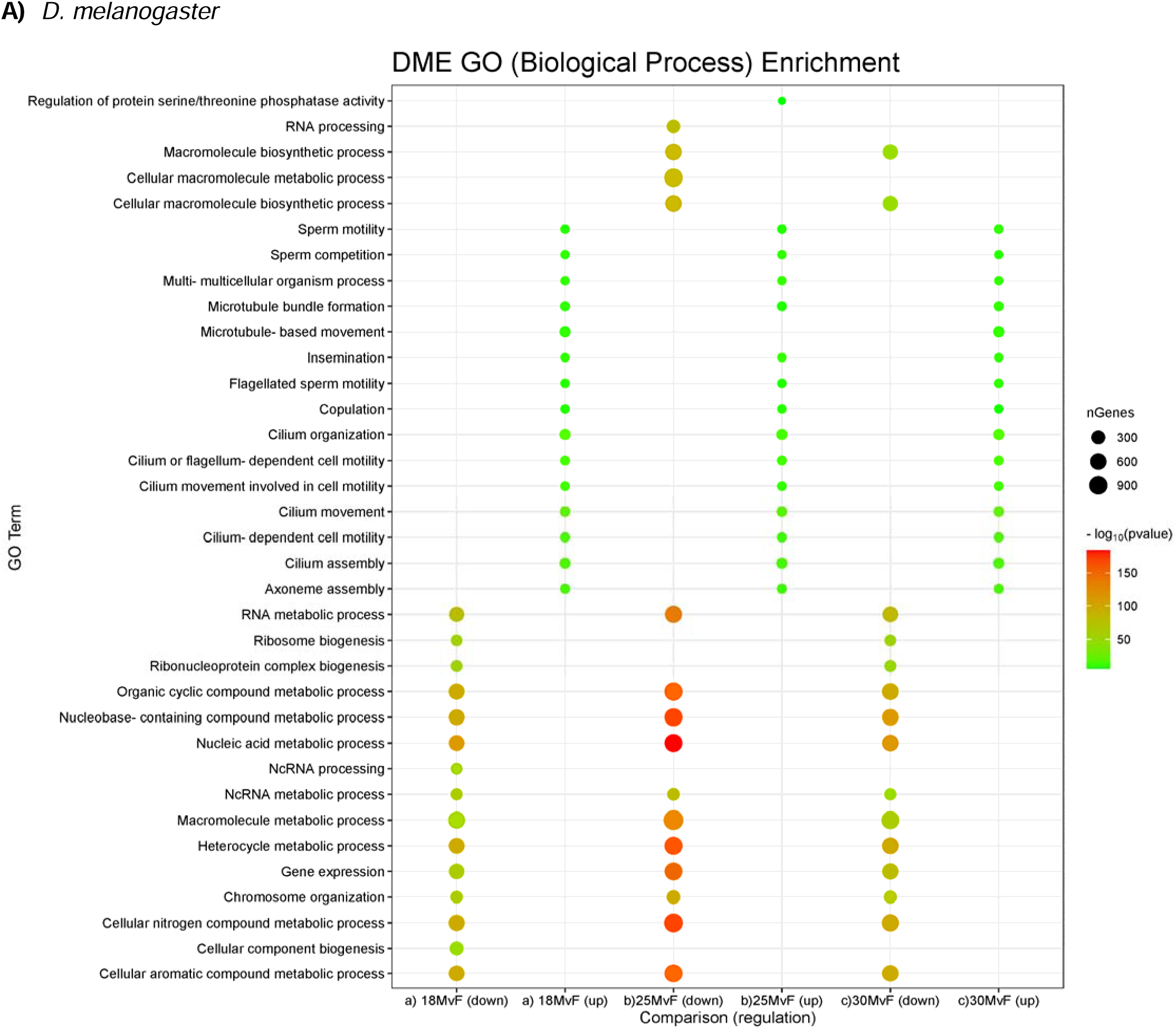
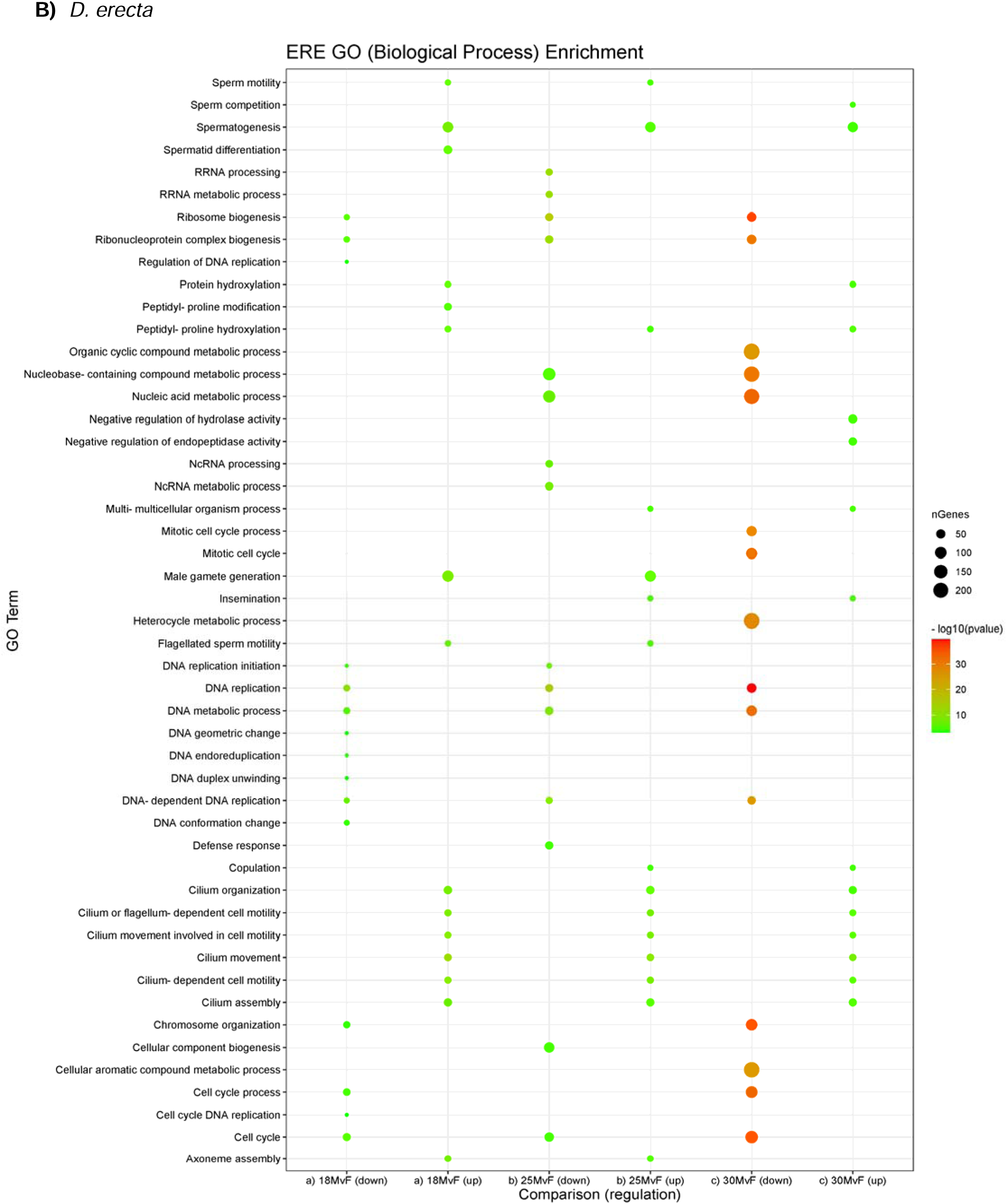
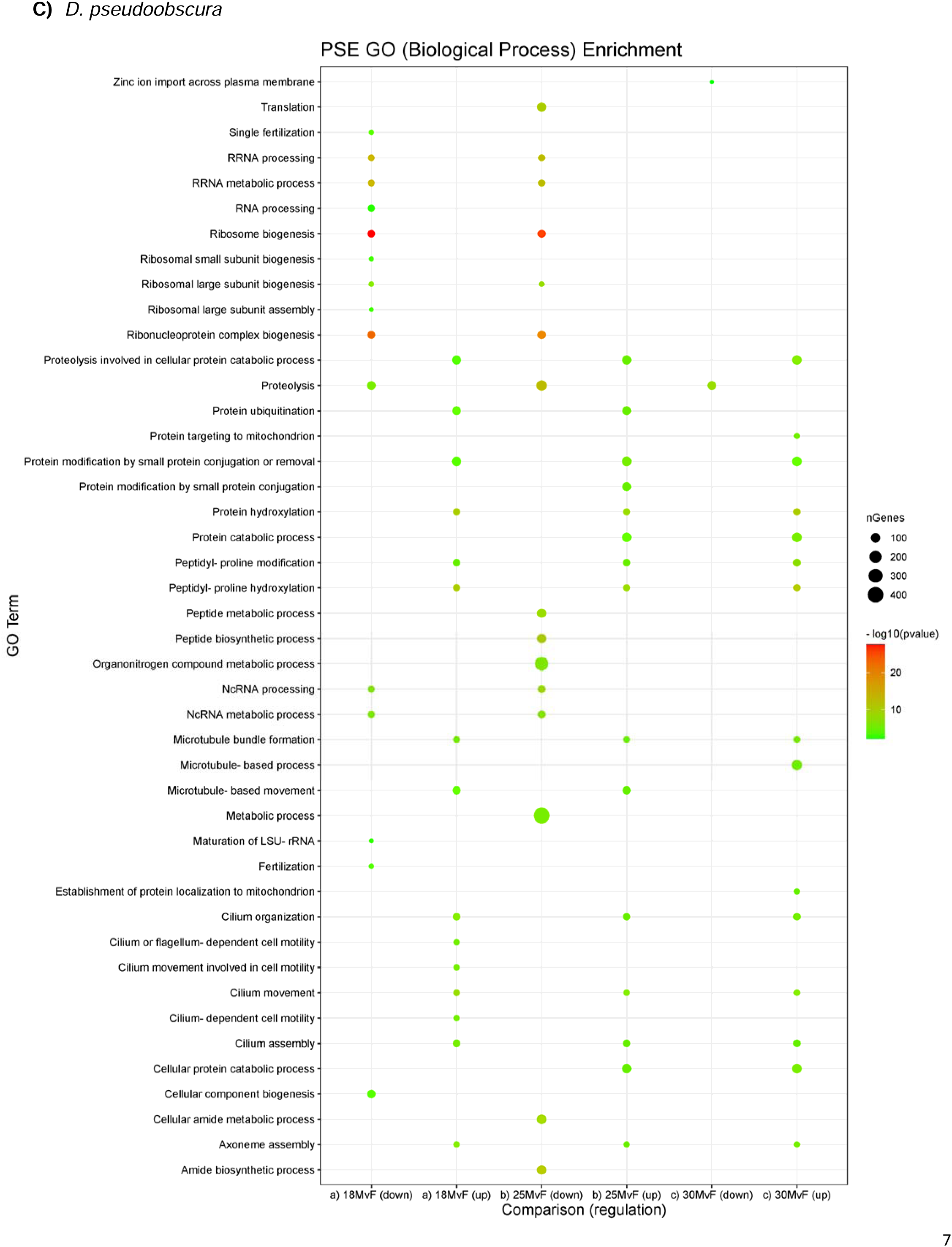
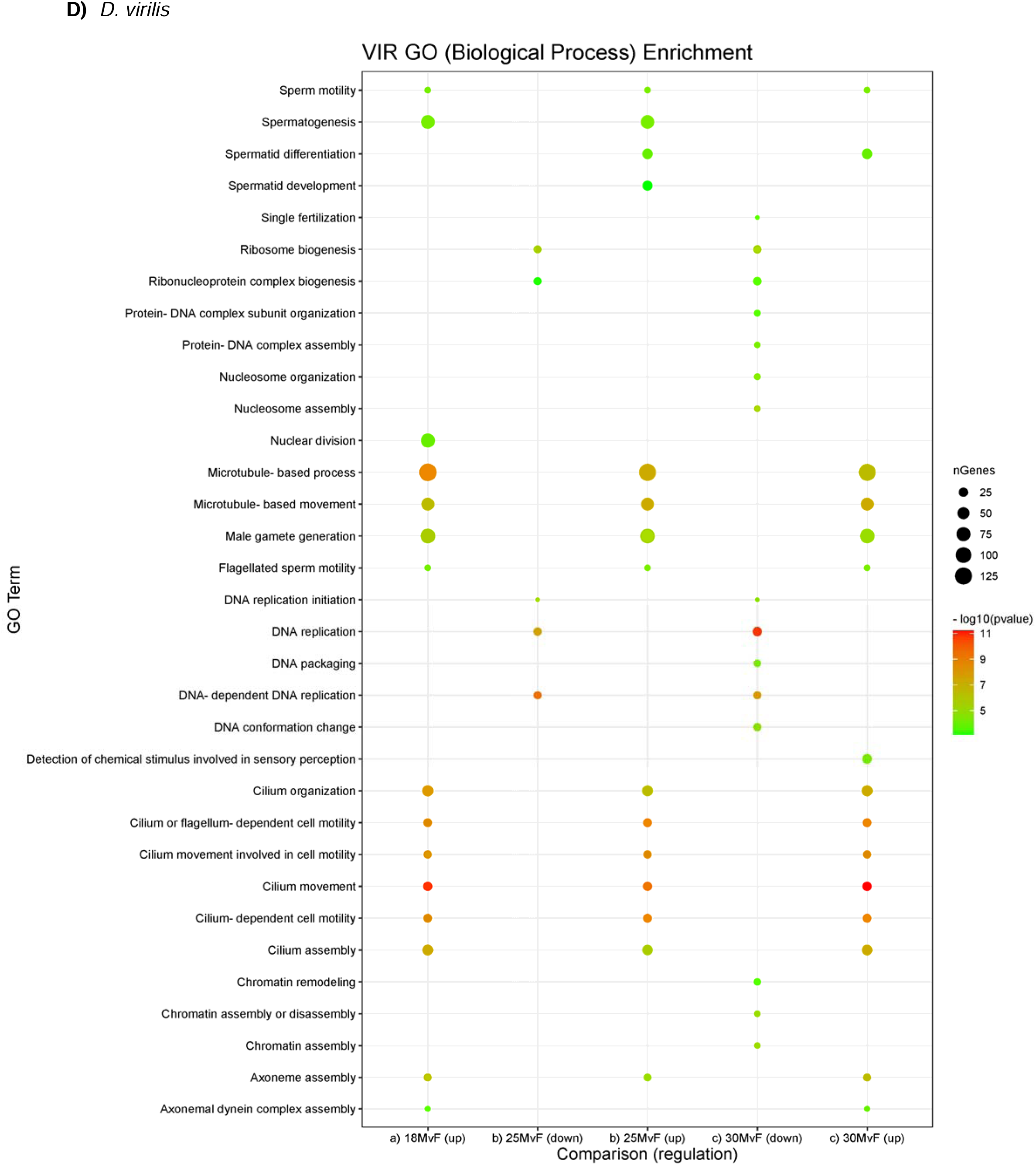
GO (Biological Process) Enrichment for Sex-biased Genes

**Fig. S3.**
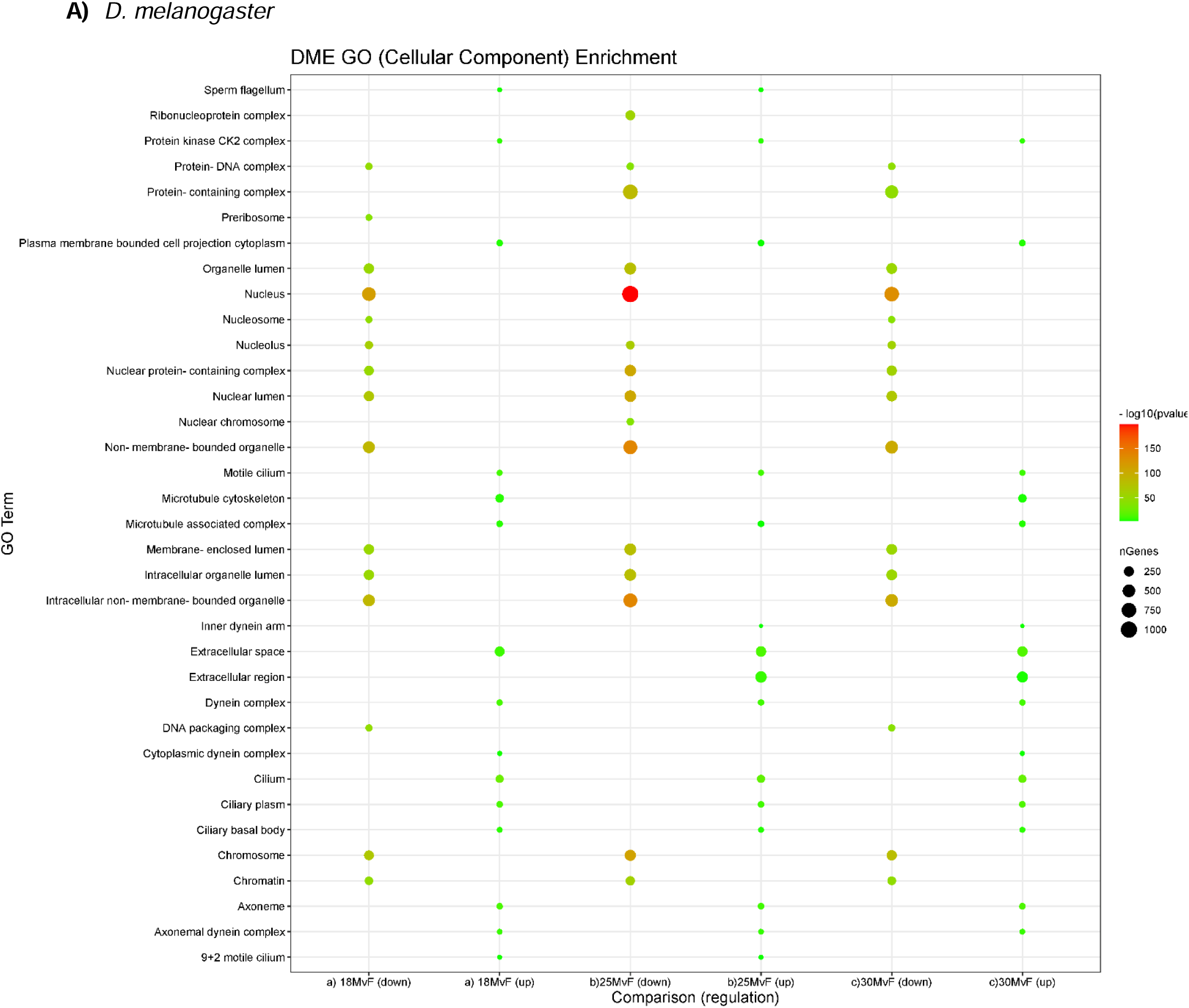
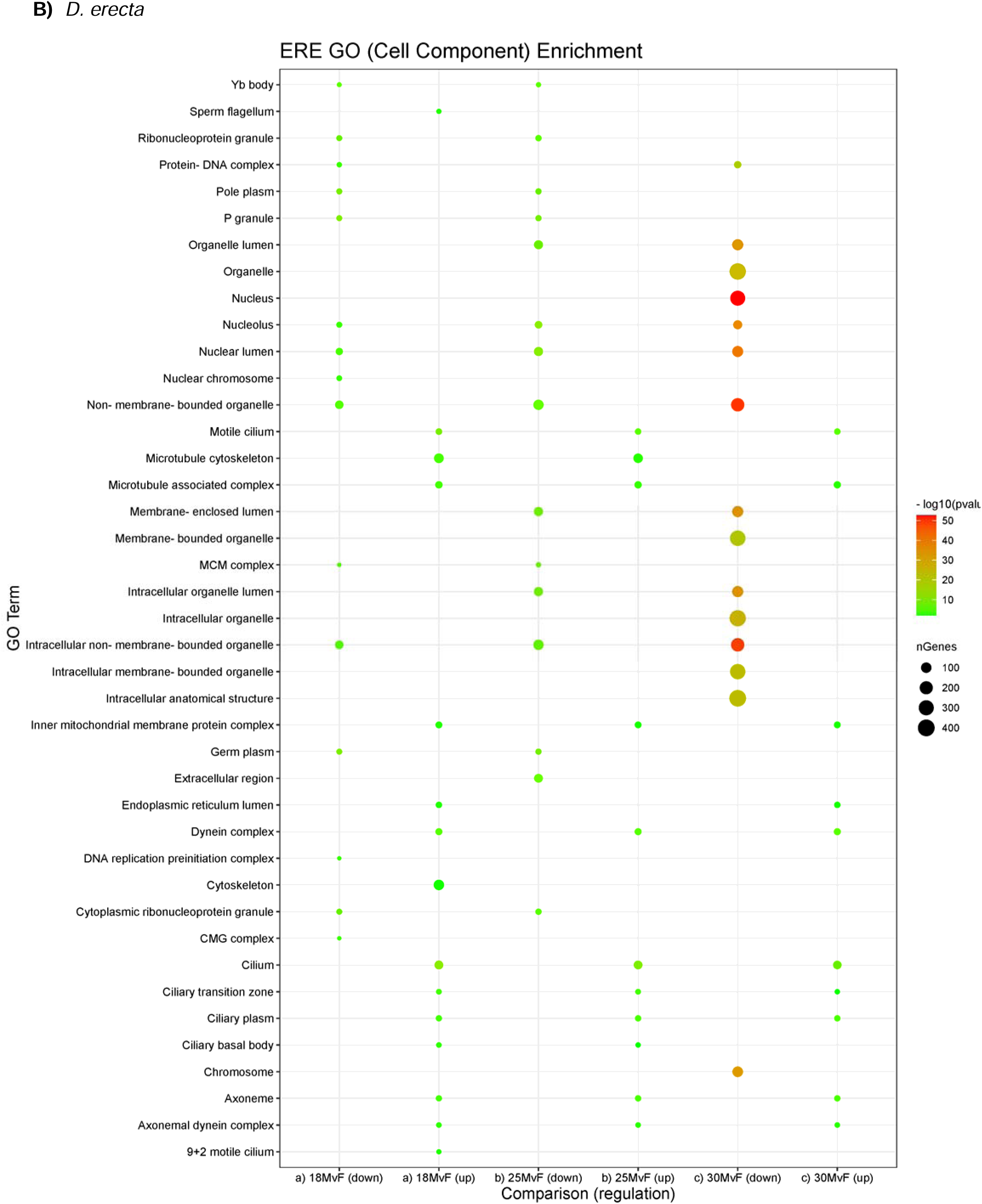
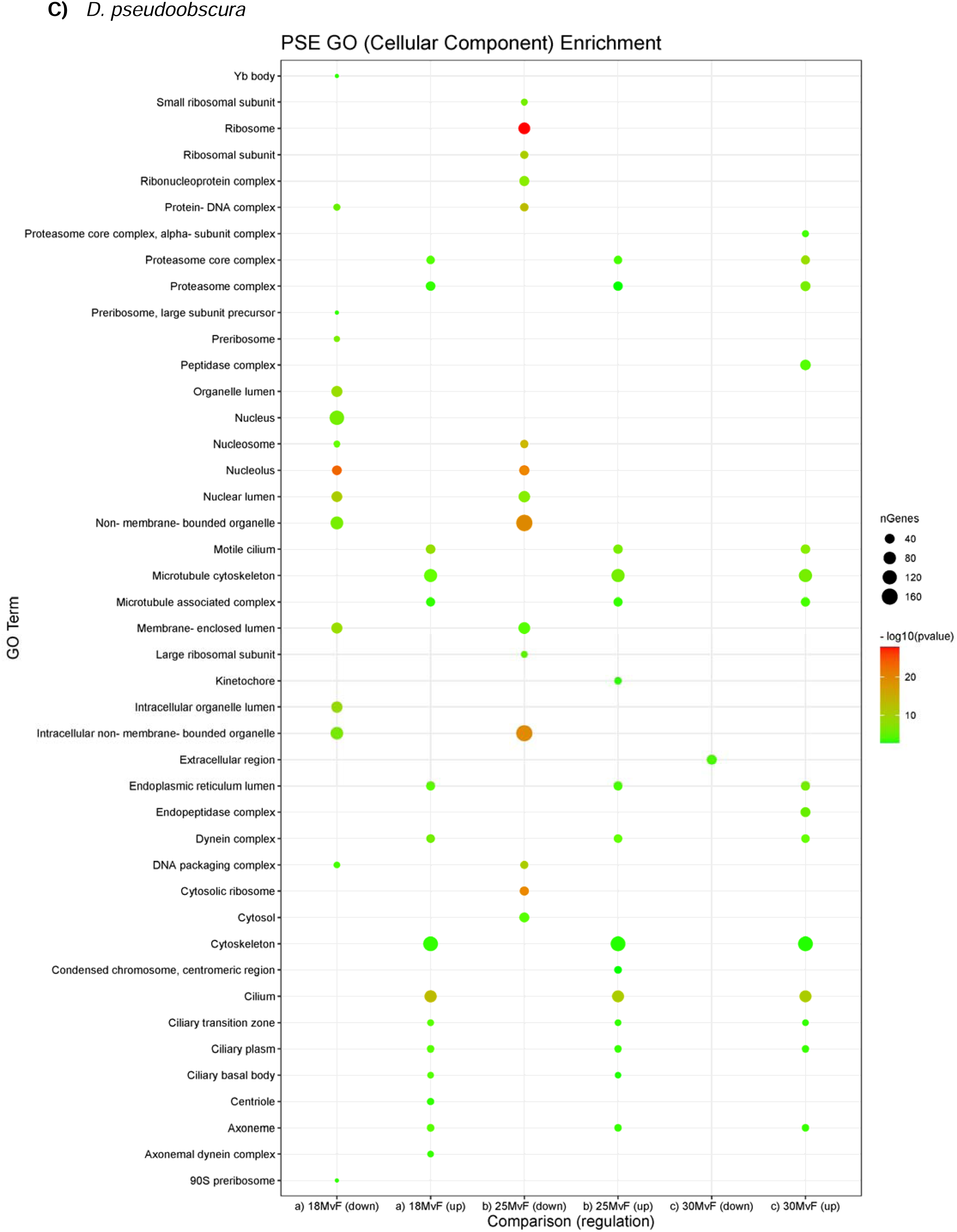
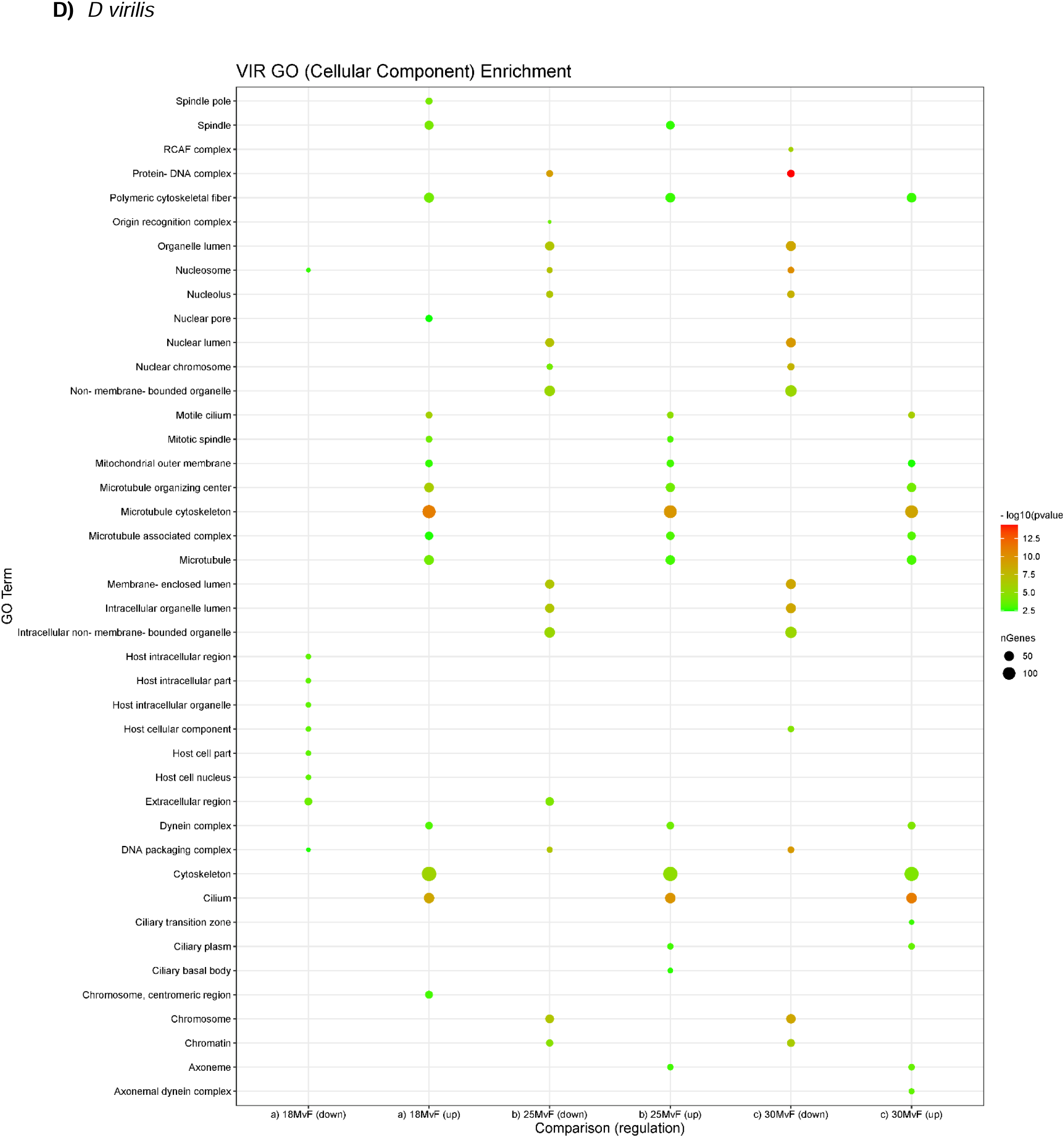
GO (Cellular Component) Enrichment for Sex-biased Genes

**Fig. S4.**
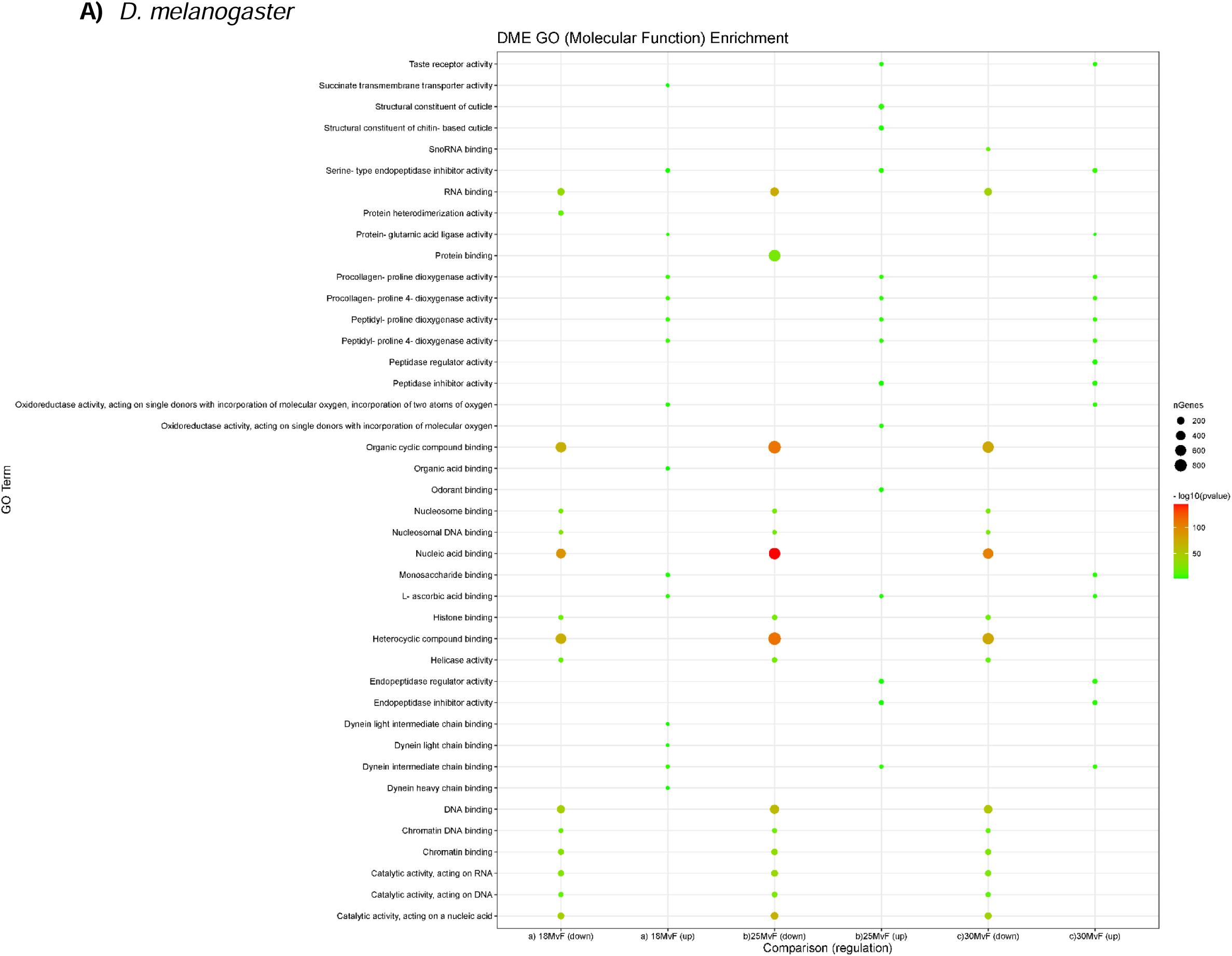
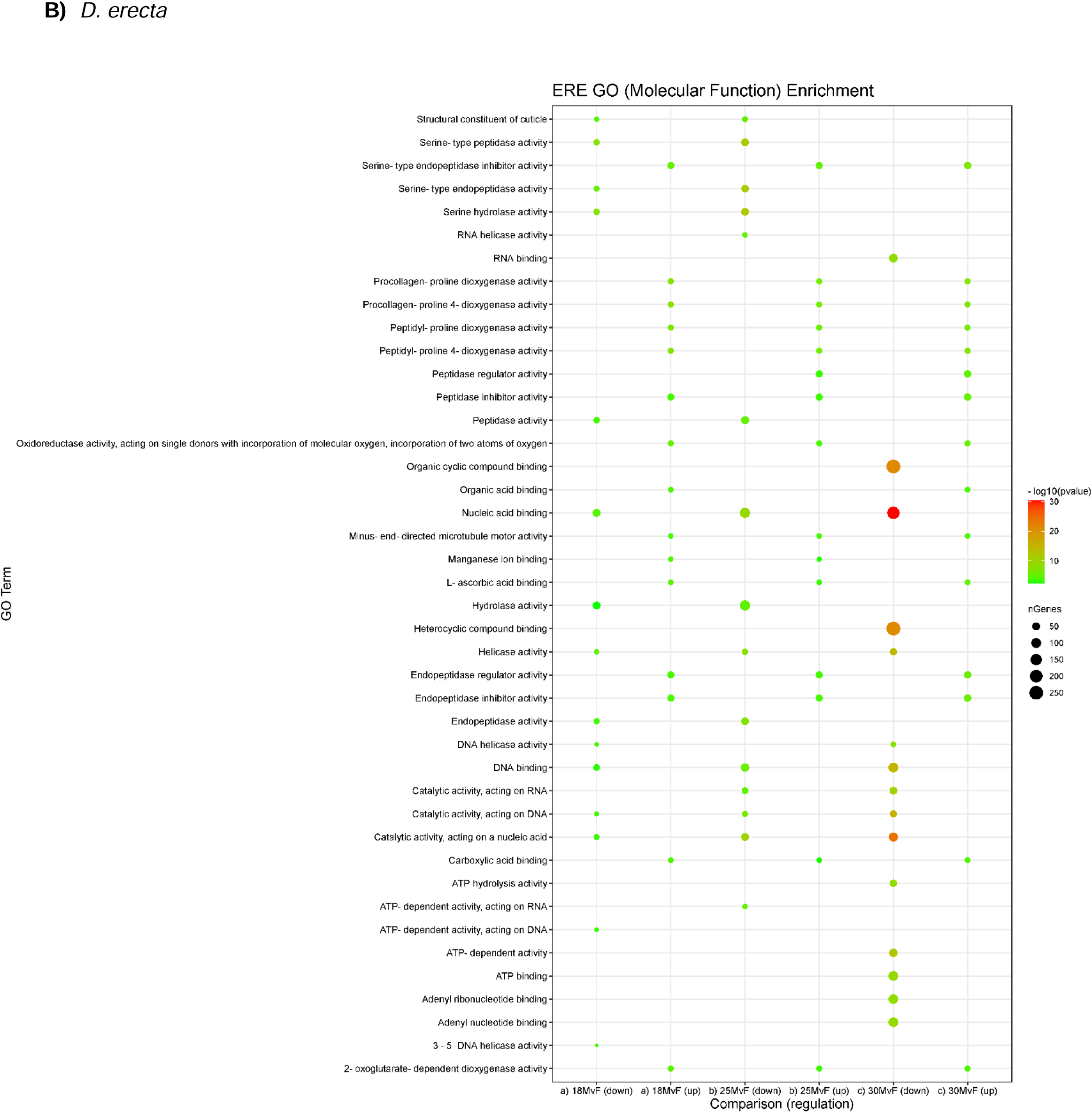
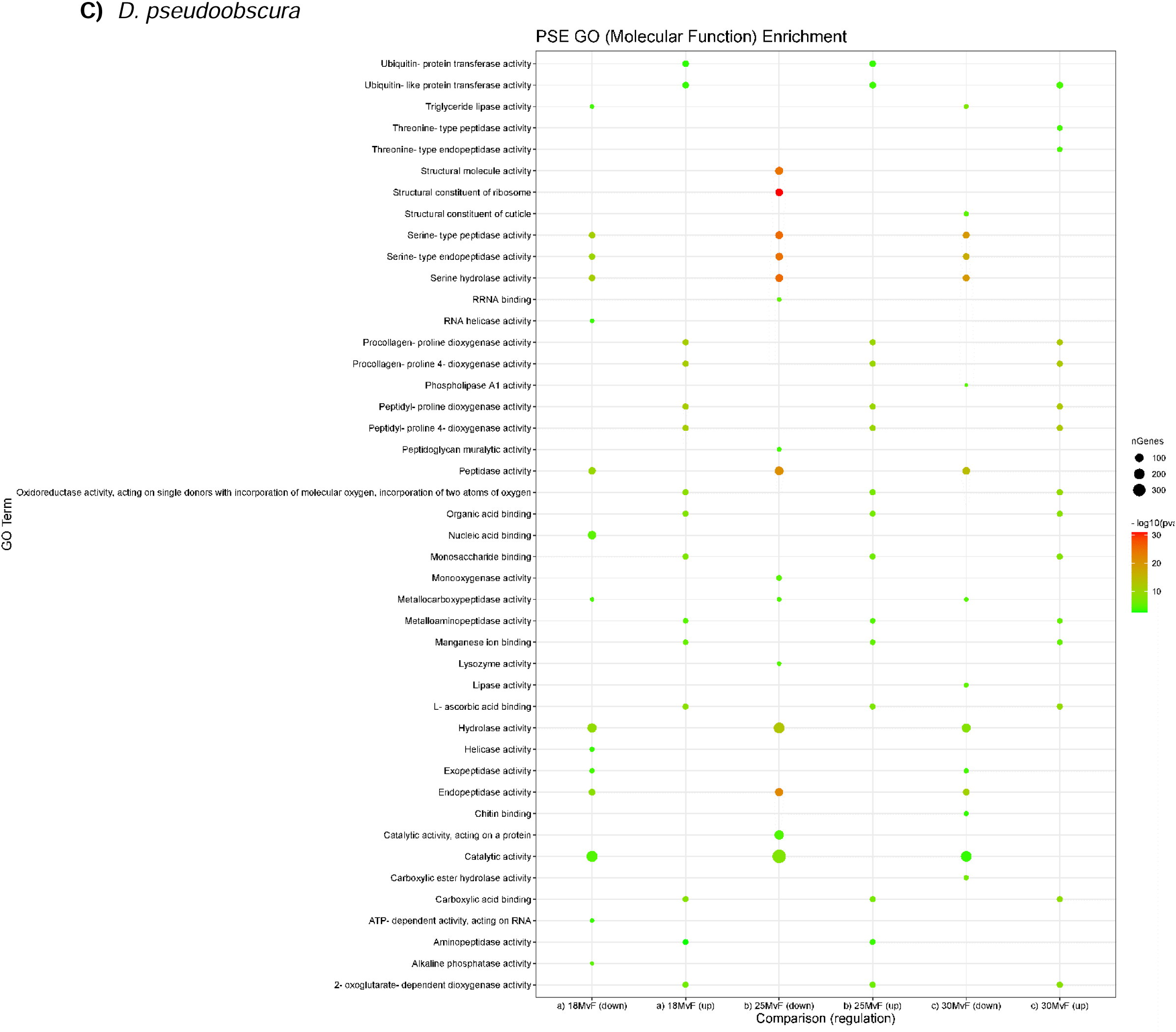
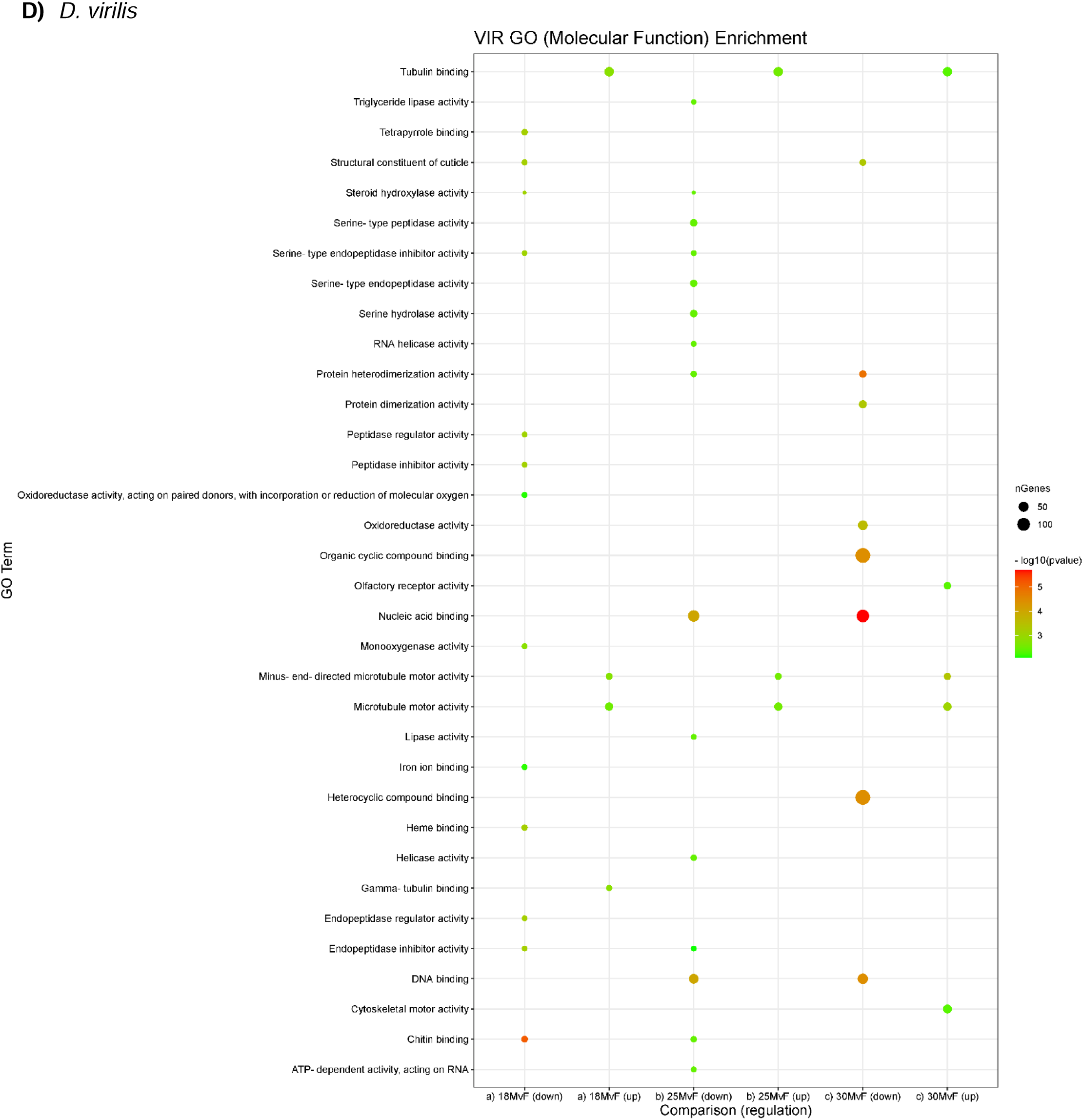
GO (Molecular Functions) Enrichment for Sex-biased Genes

**Fig. S5.**
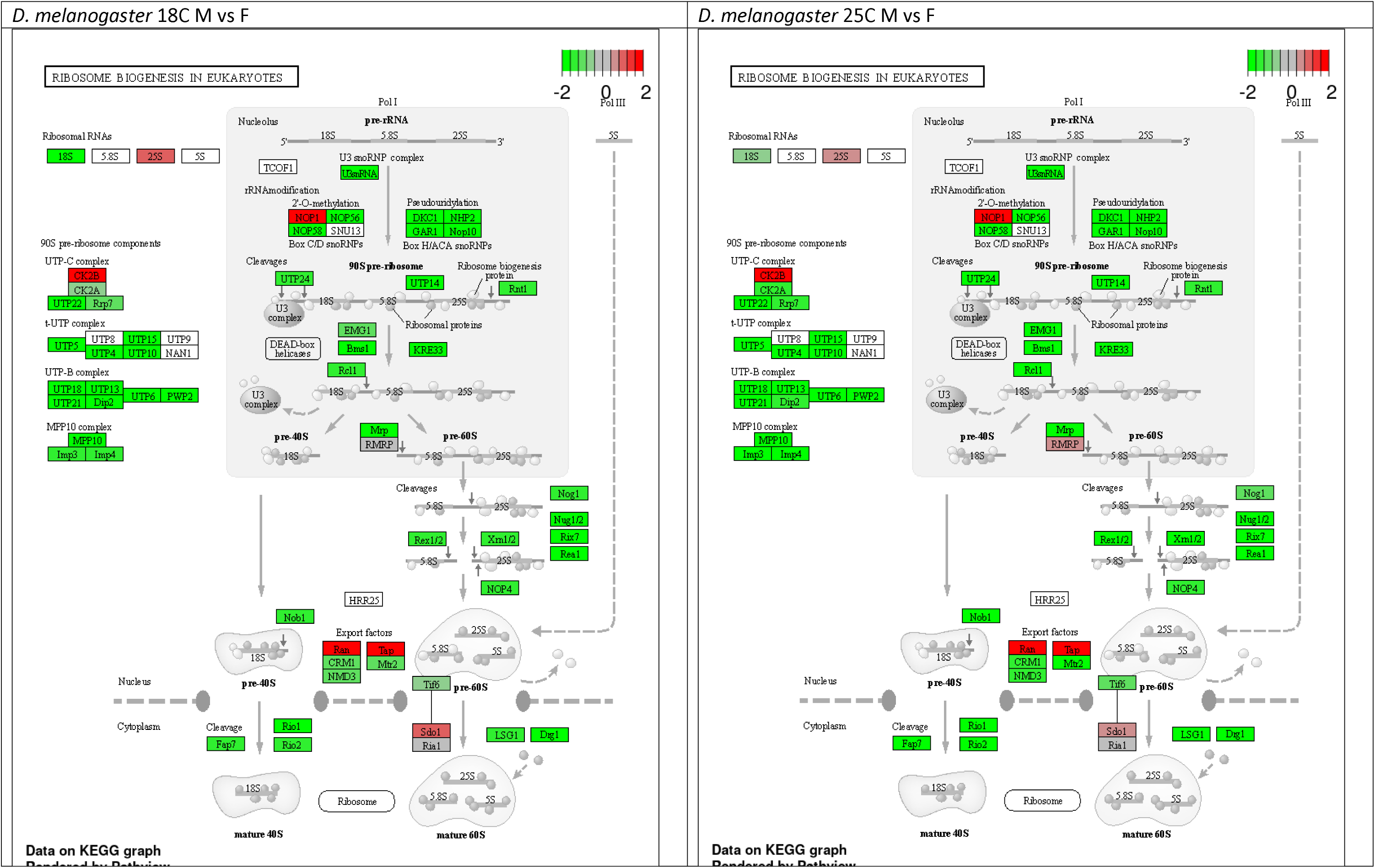
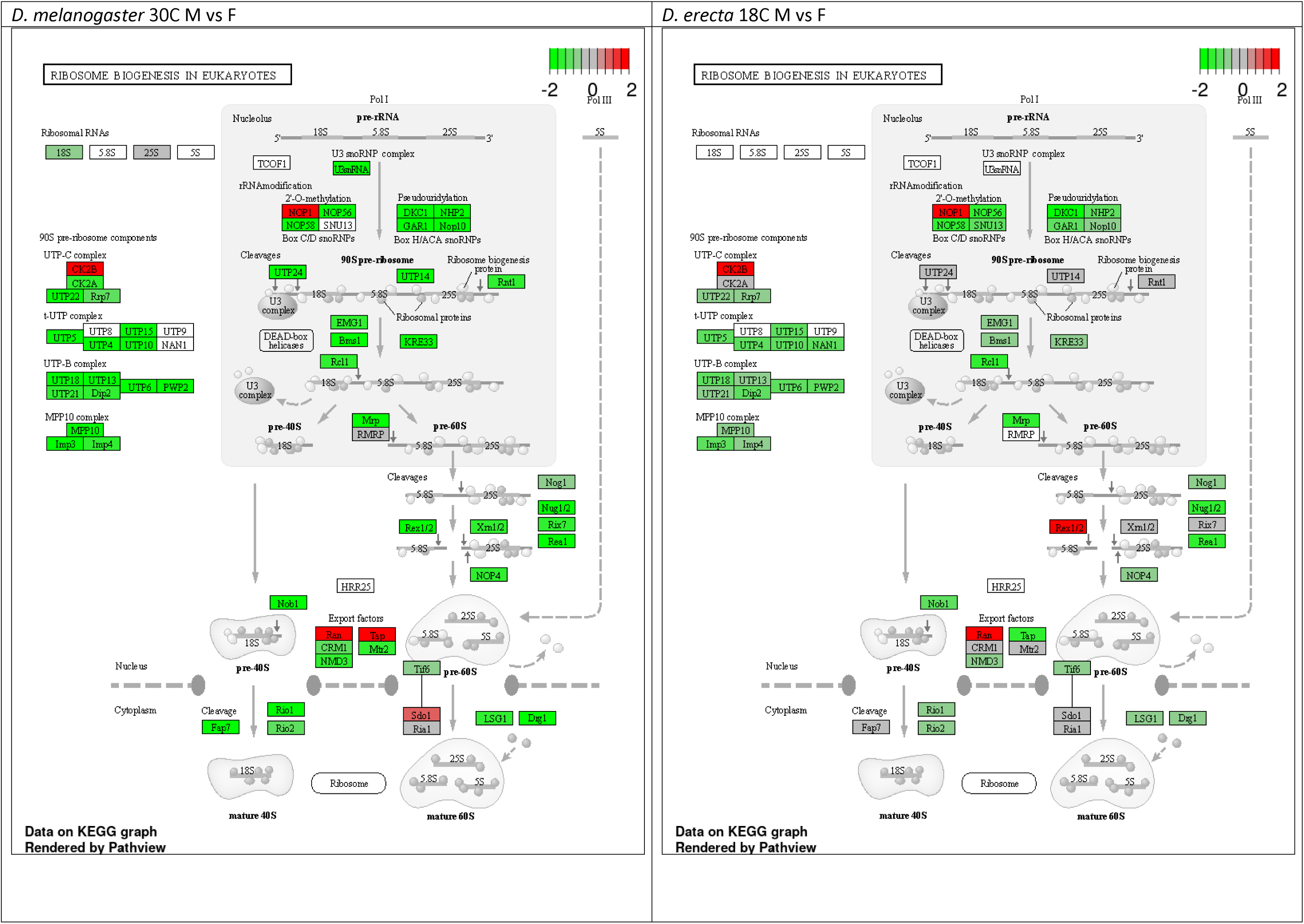
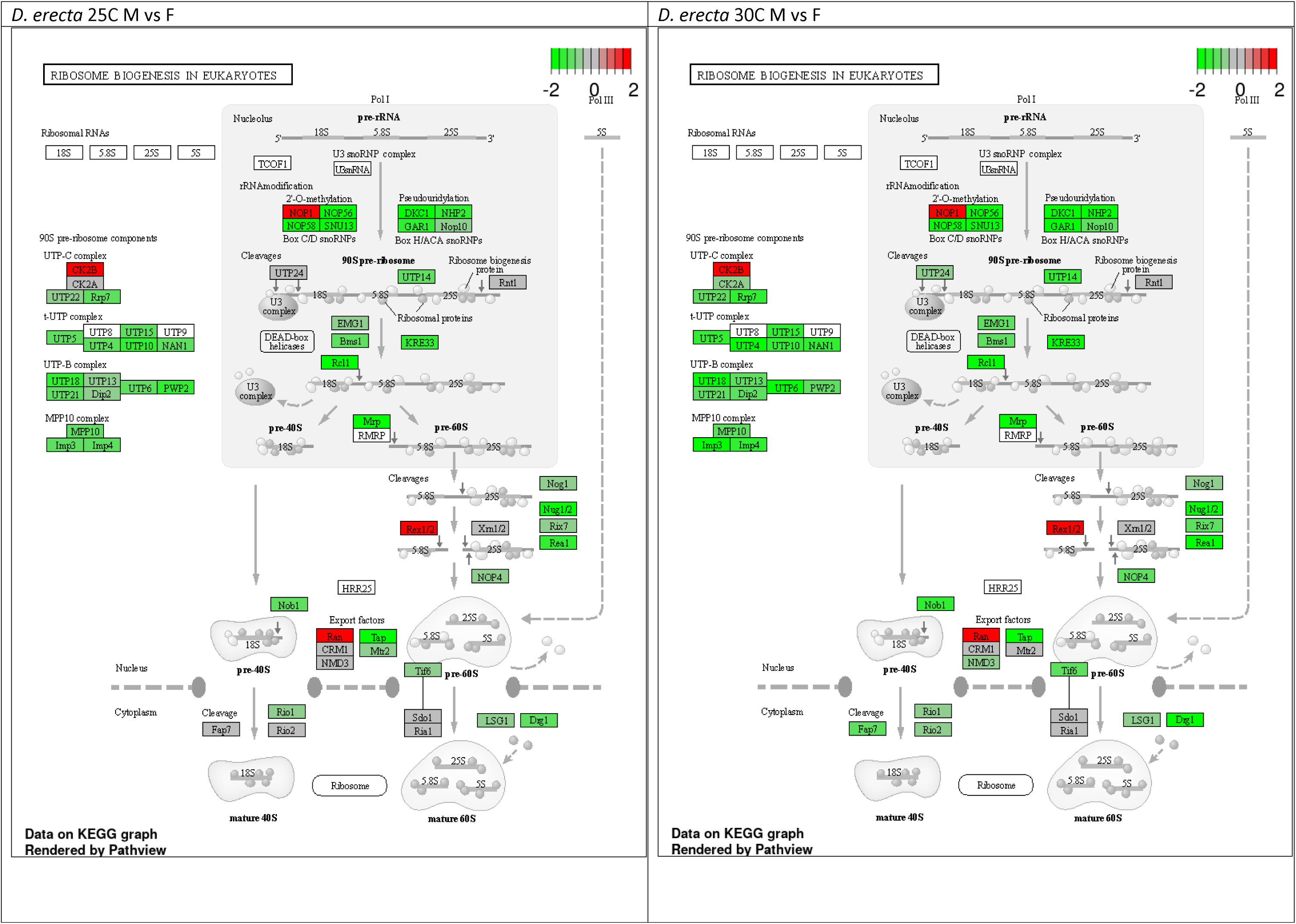
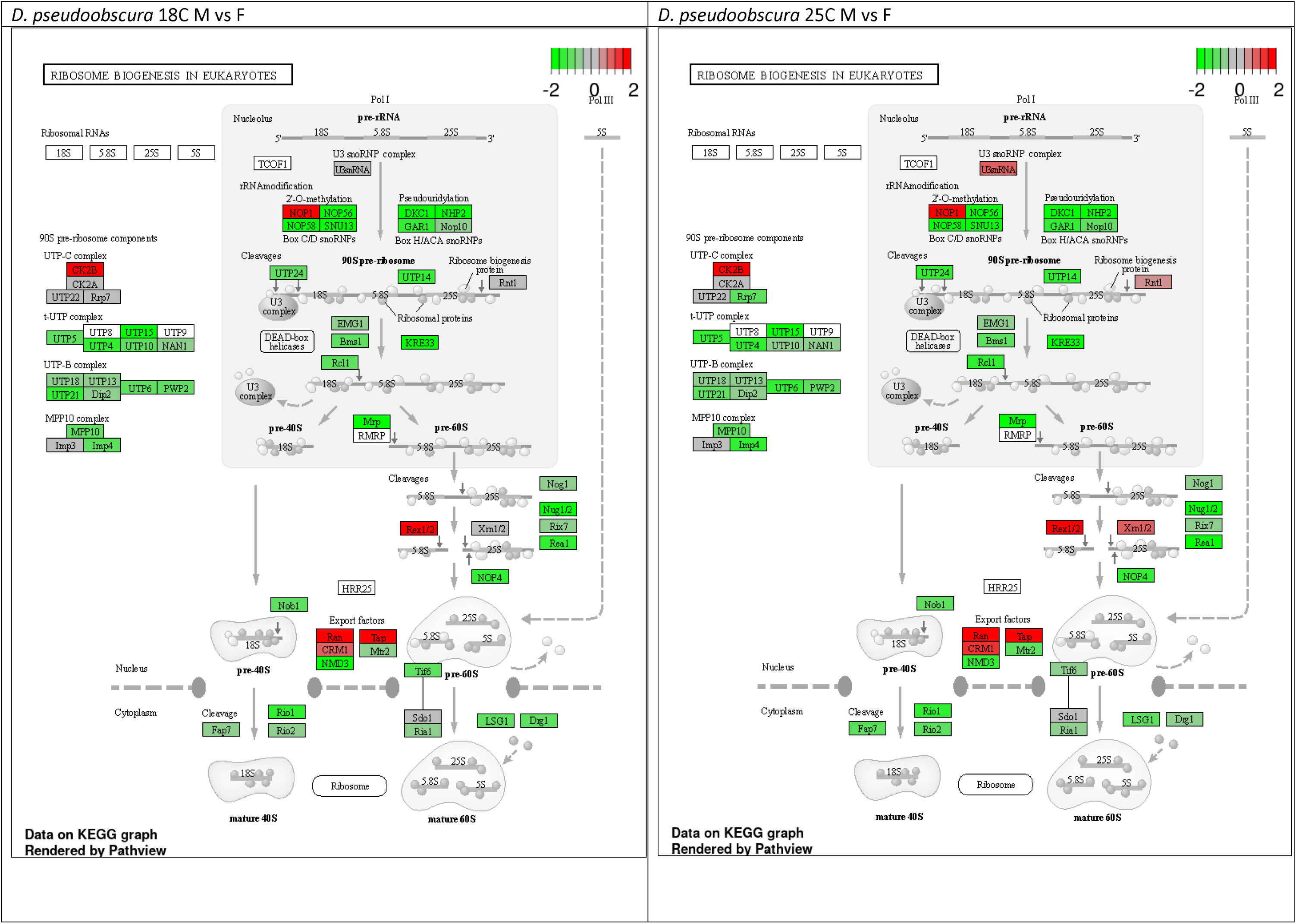
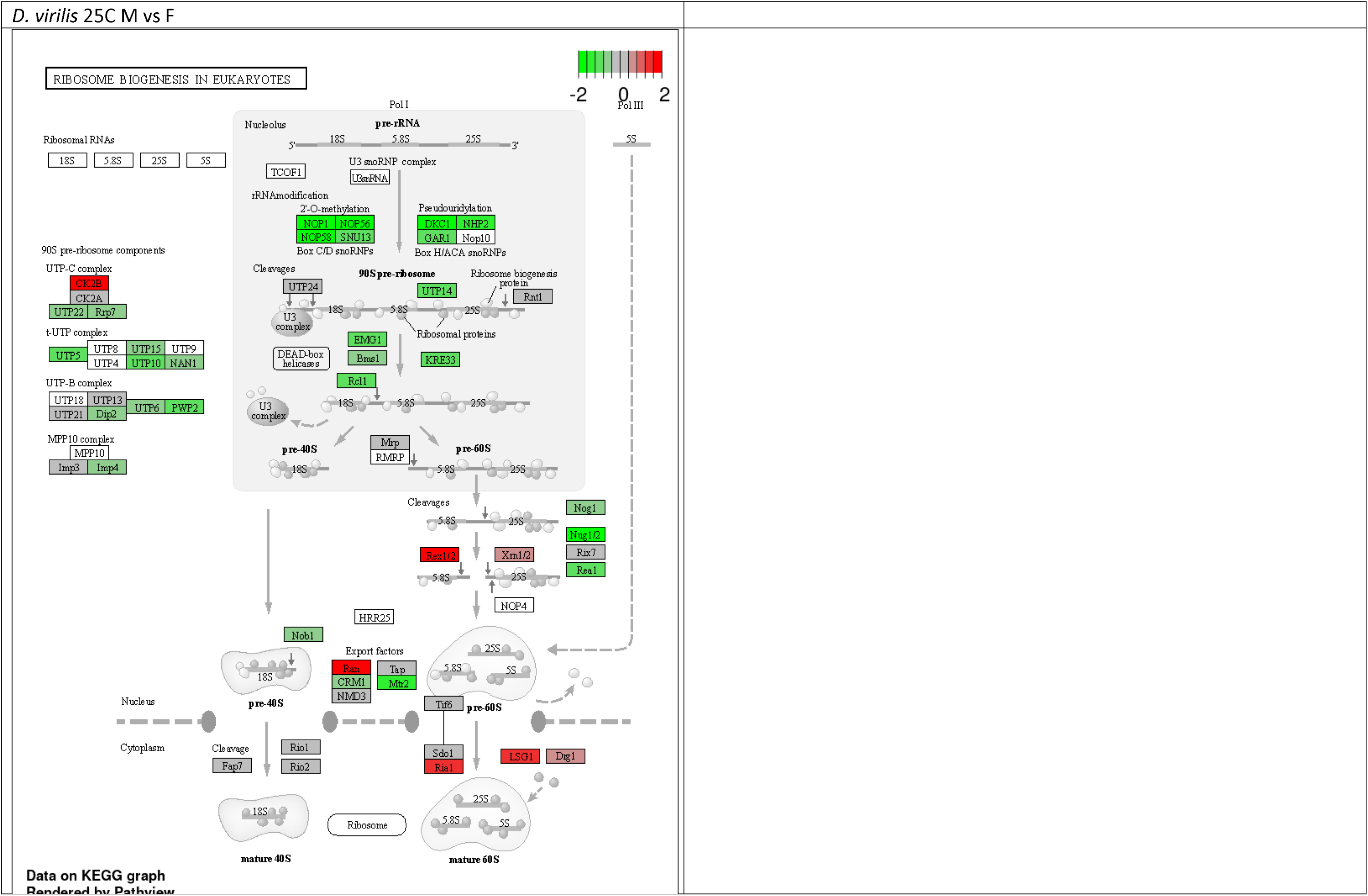
Regulation of genes in Ribosome biogenesis in eukaryotes pathway across different species under sexual comparison of different temperature.

**Fig. S6.**
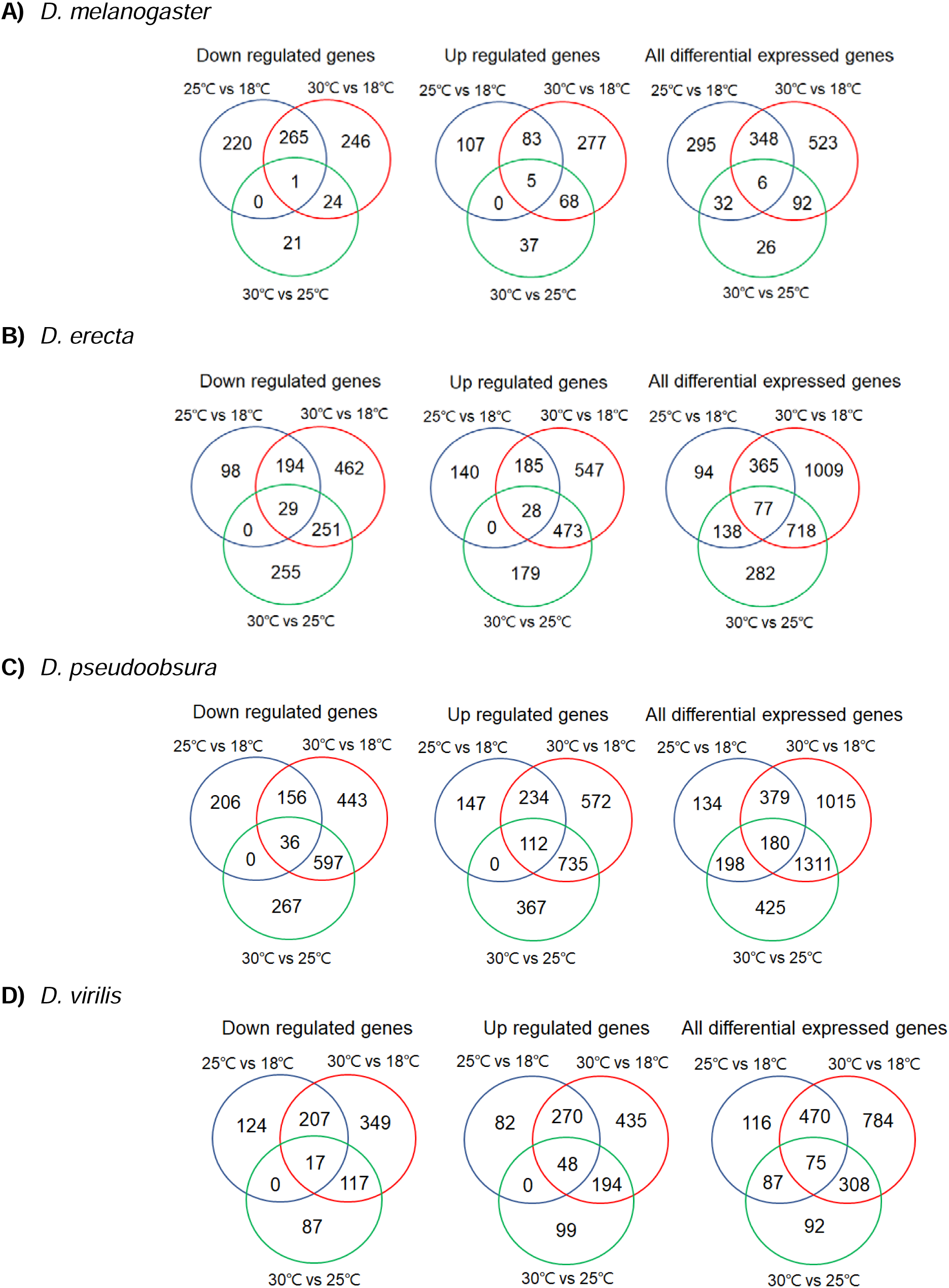
Venn diagrams showing differential expressed genes in female under different temperature regimes

**Fig. S7.**
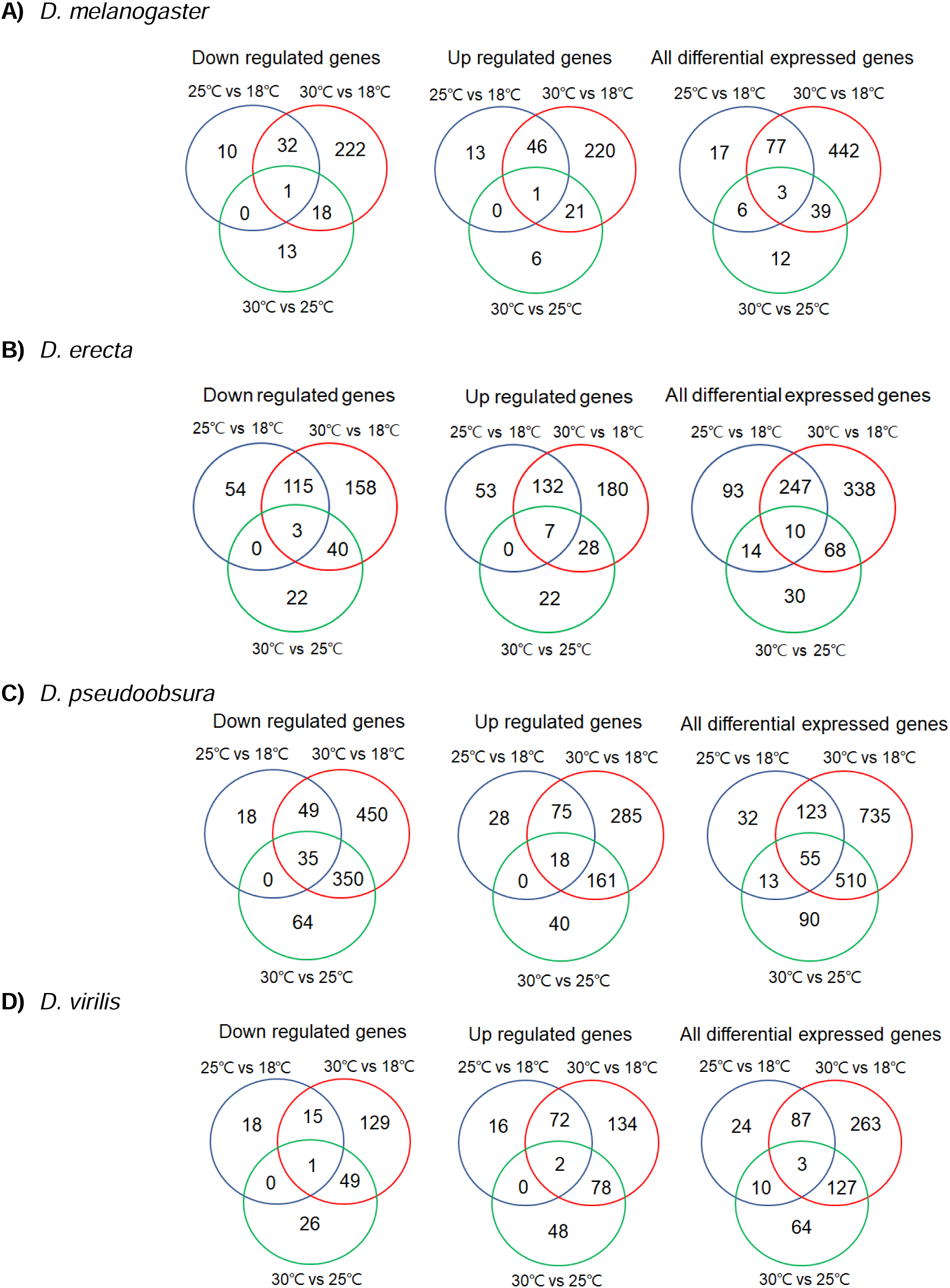
Venn diagrams showing differential expressed genes in male under different temperature regimes

**Fig. S8.**
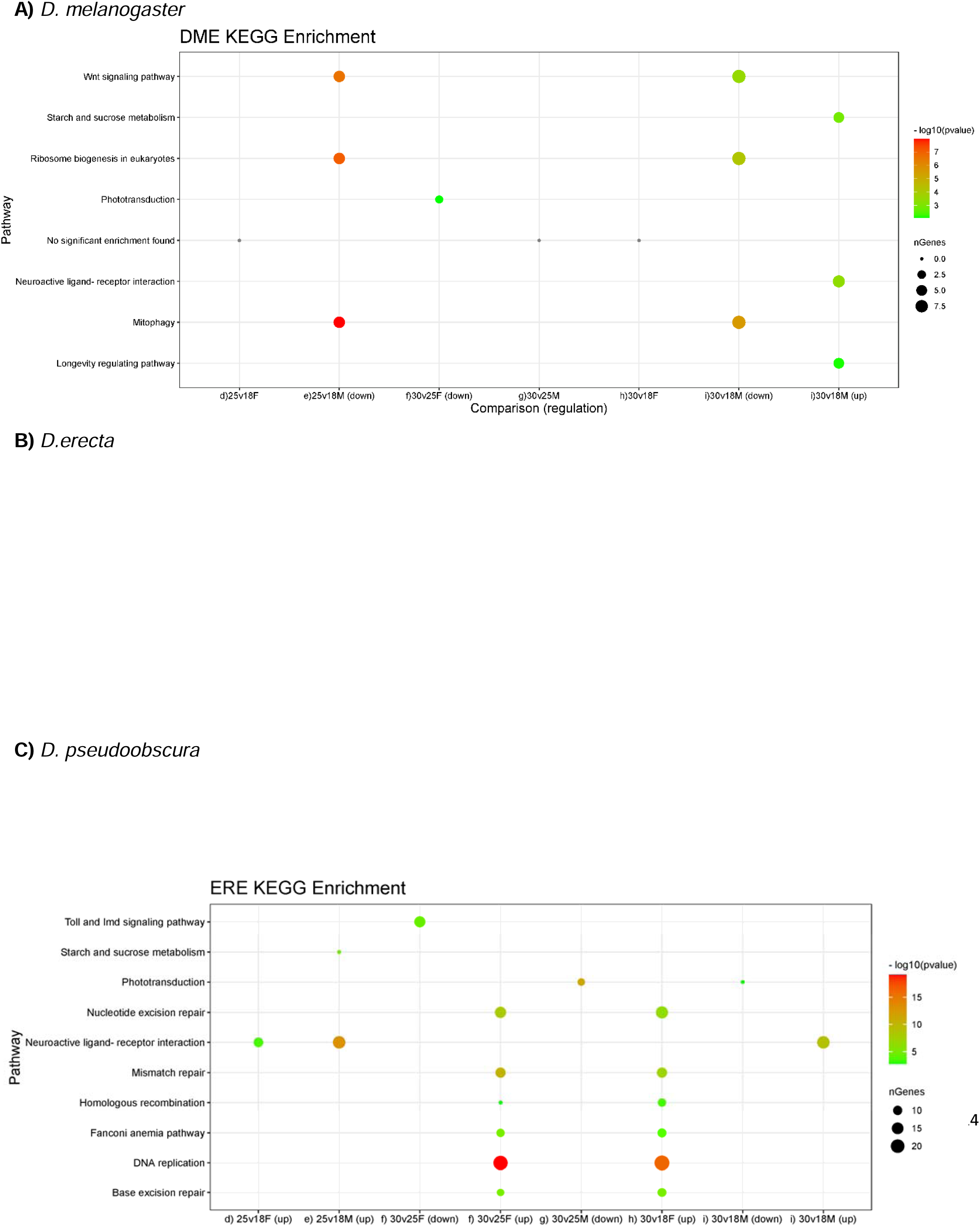
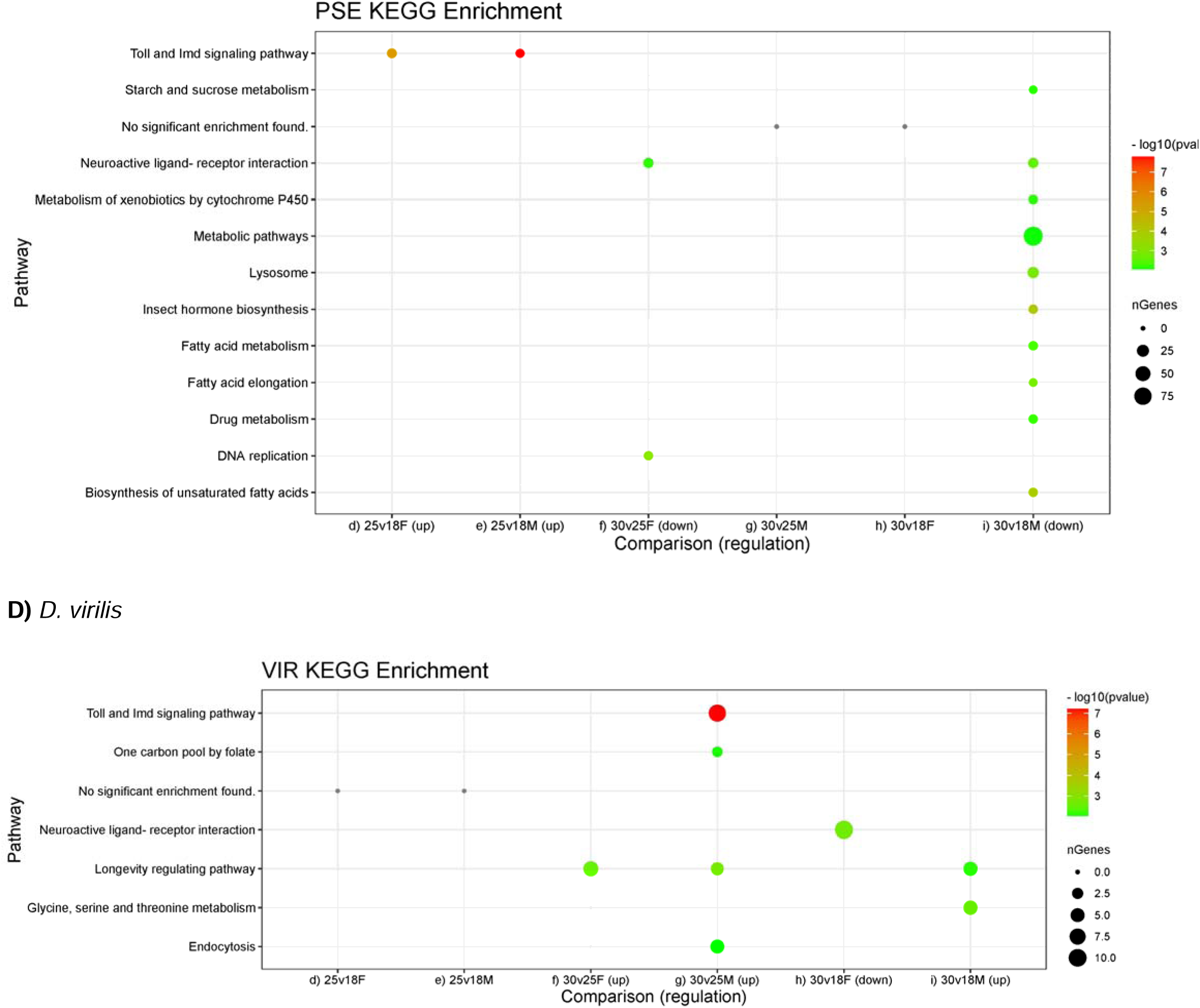
KEGG Pathway enrichment for differential expressed genes under different temperatures of 4 Drosophila species

**Fig. S9.**
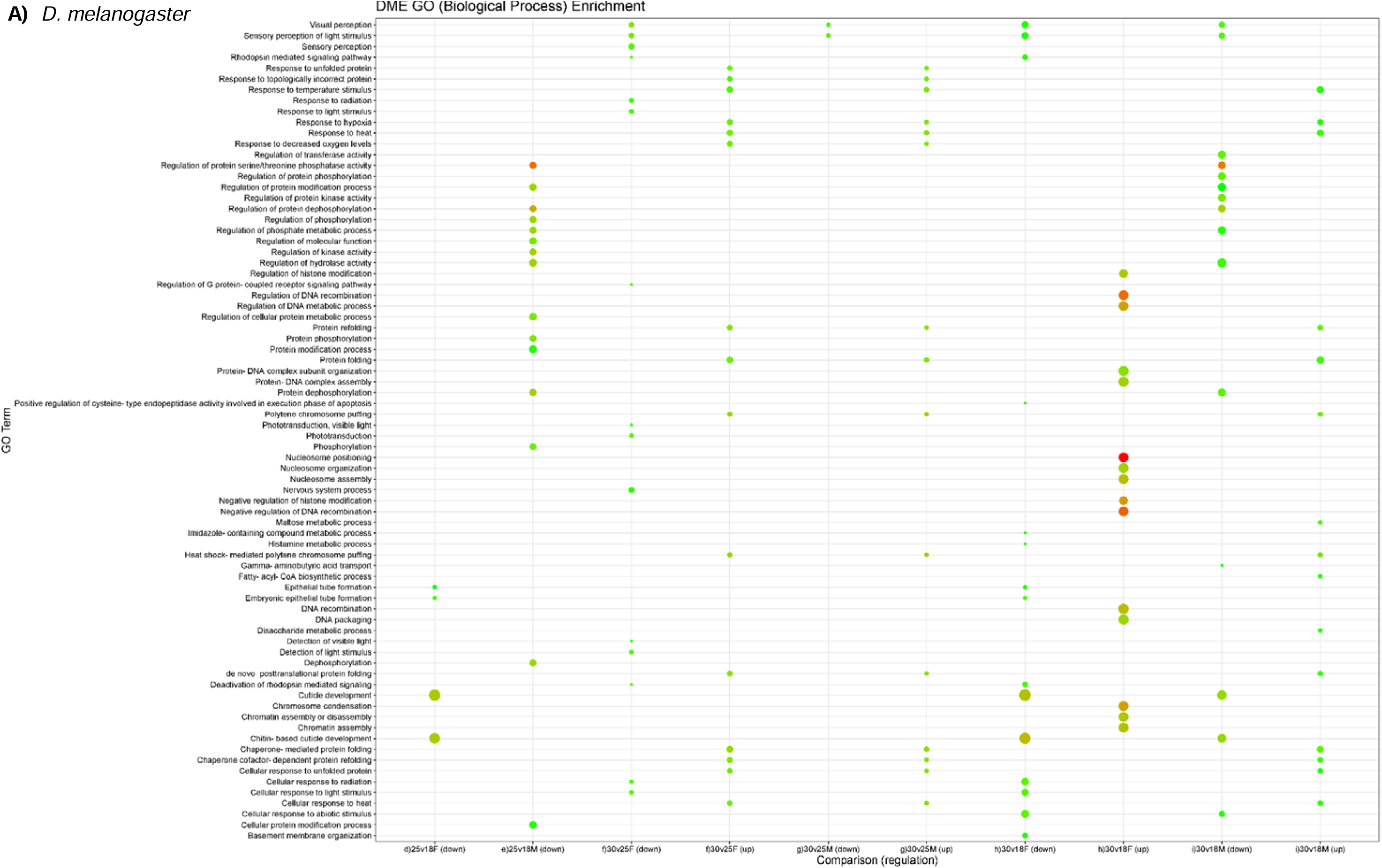
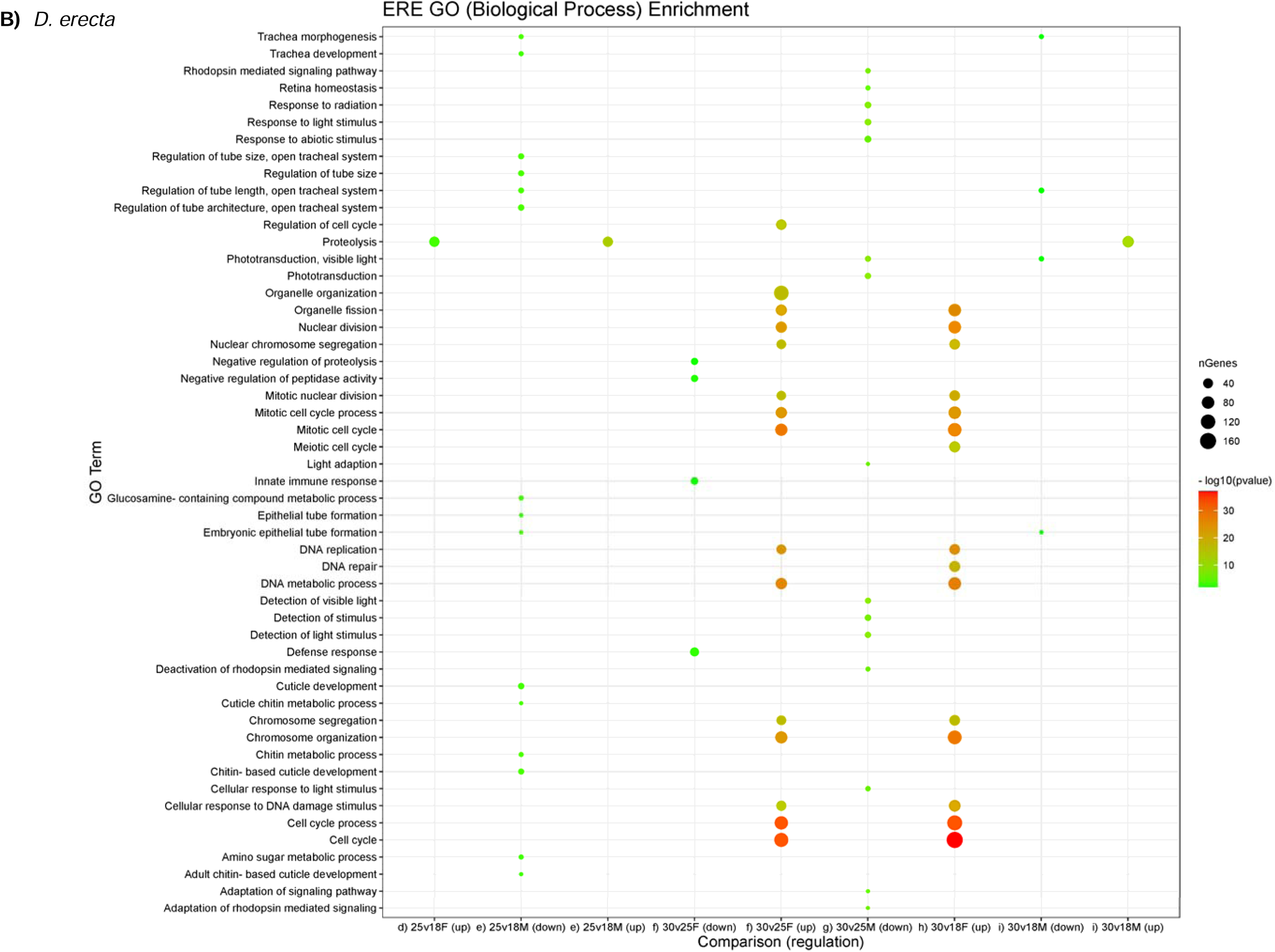
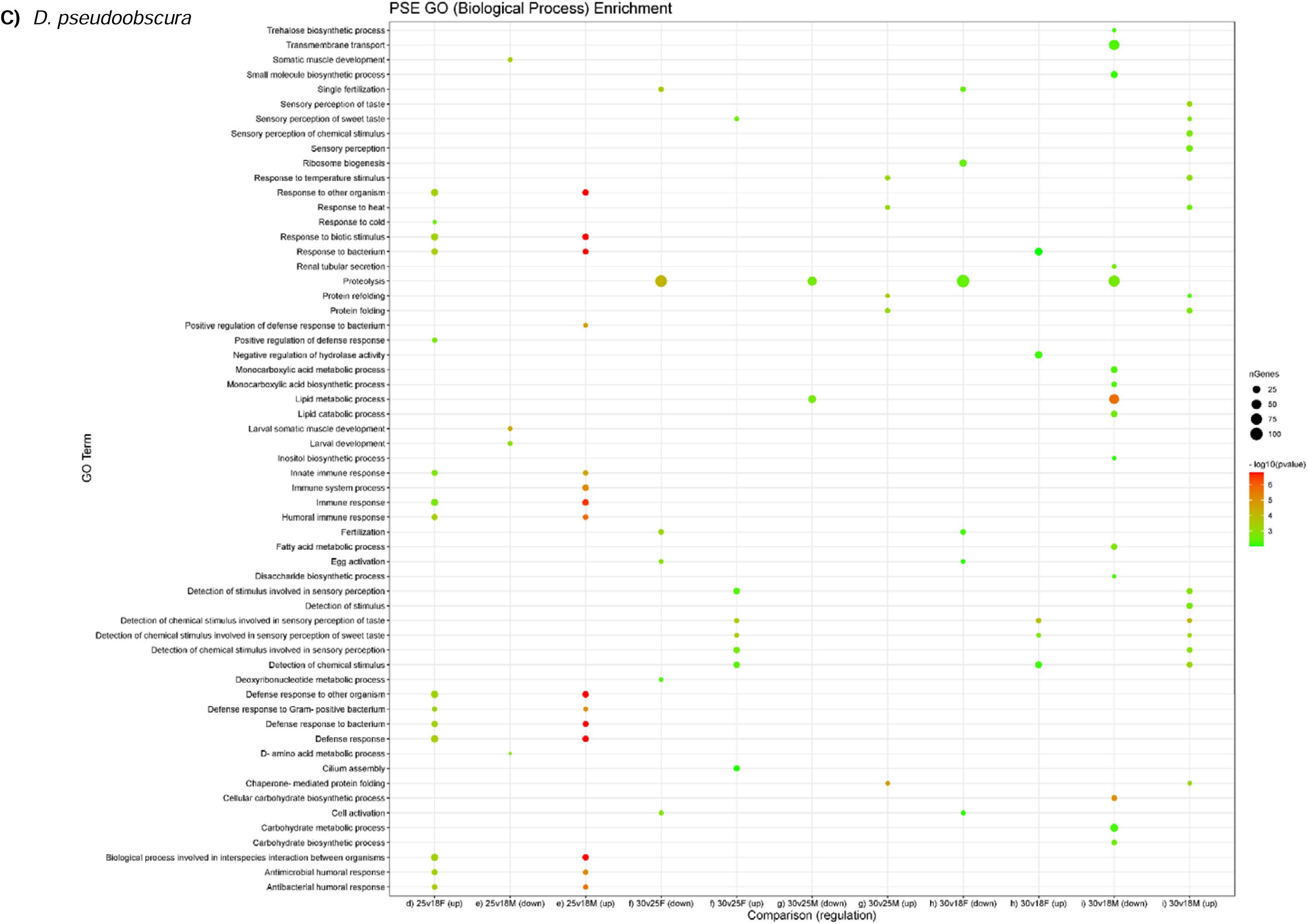
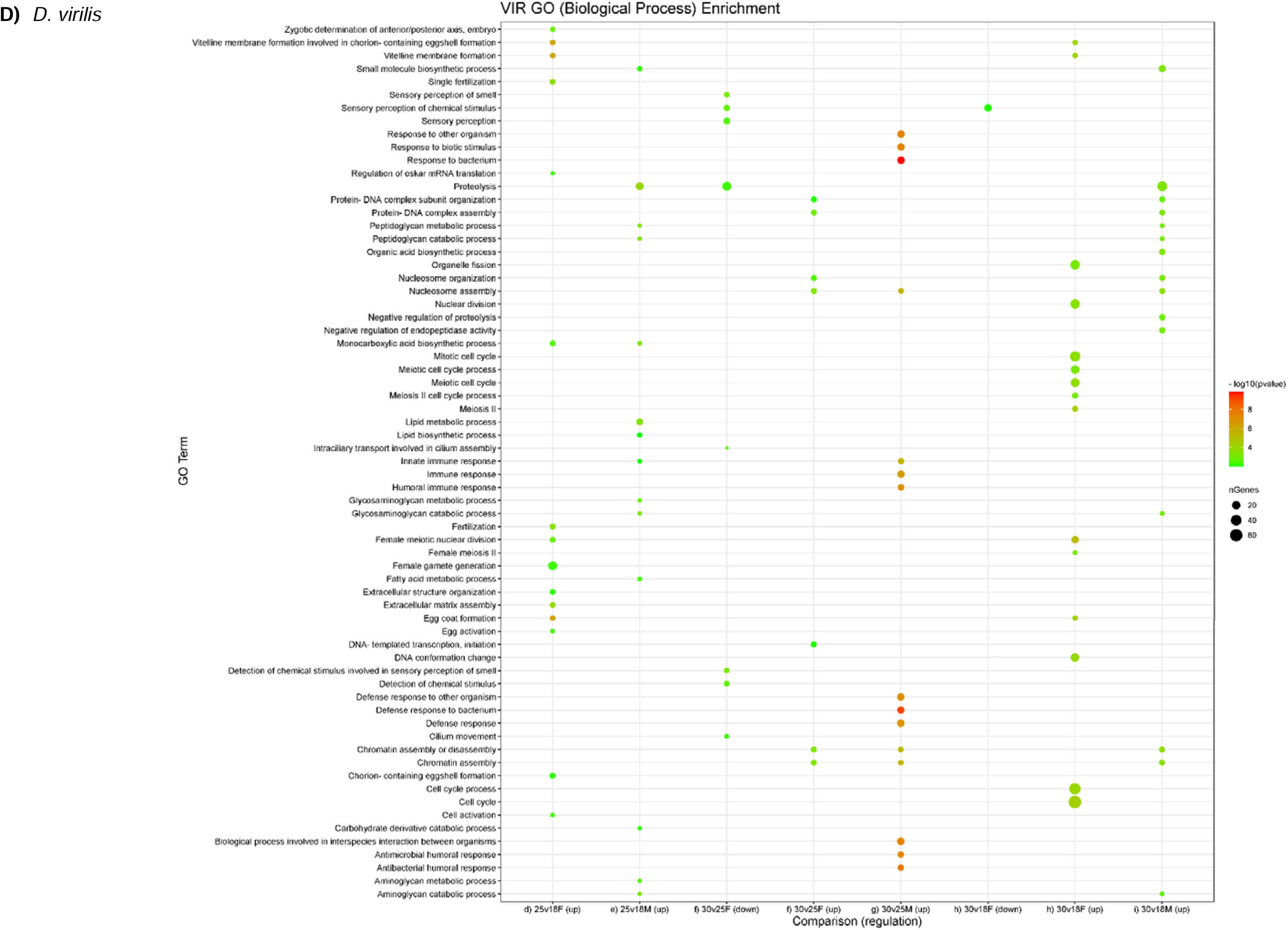
GO (Biological Process) Enrichment for differential expressed genes under different temperature

**Fig. S10.**
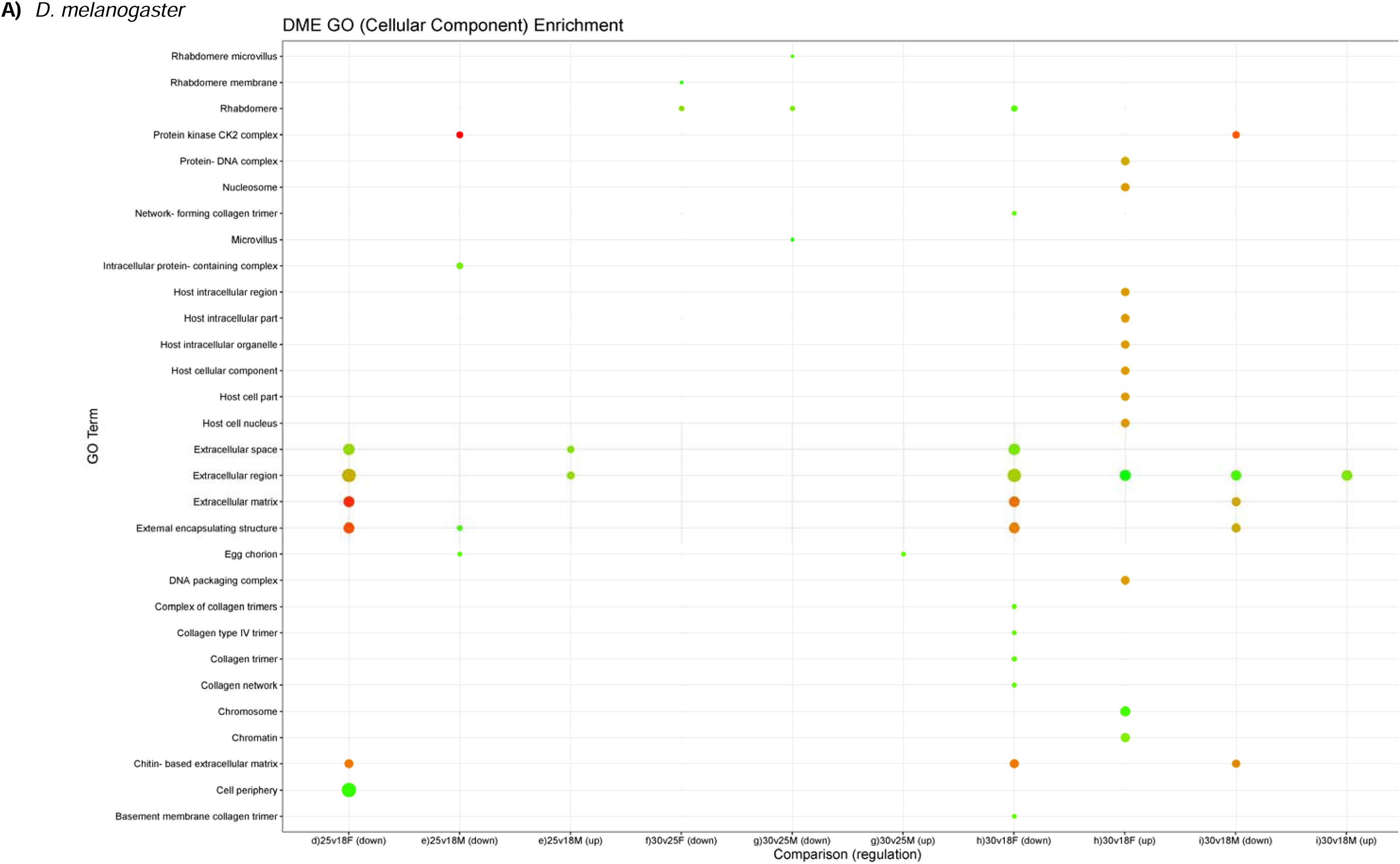
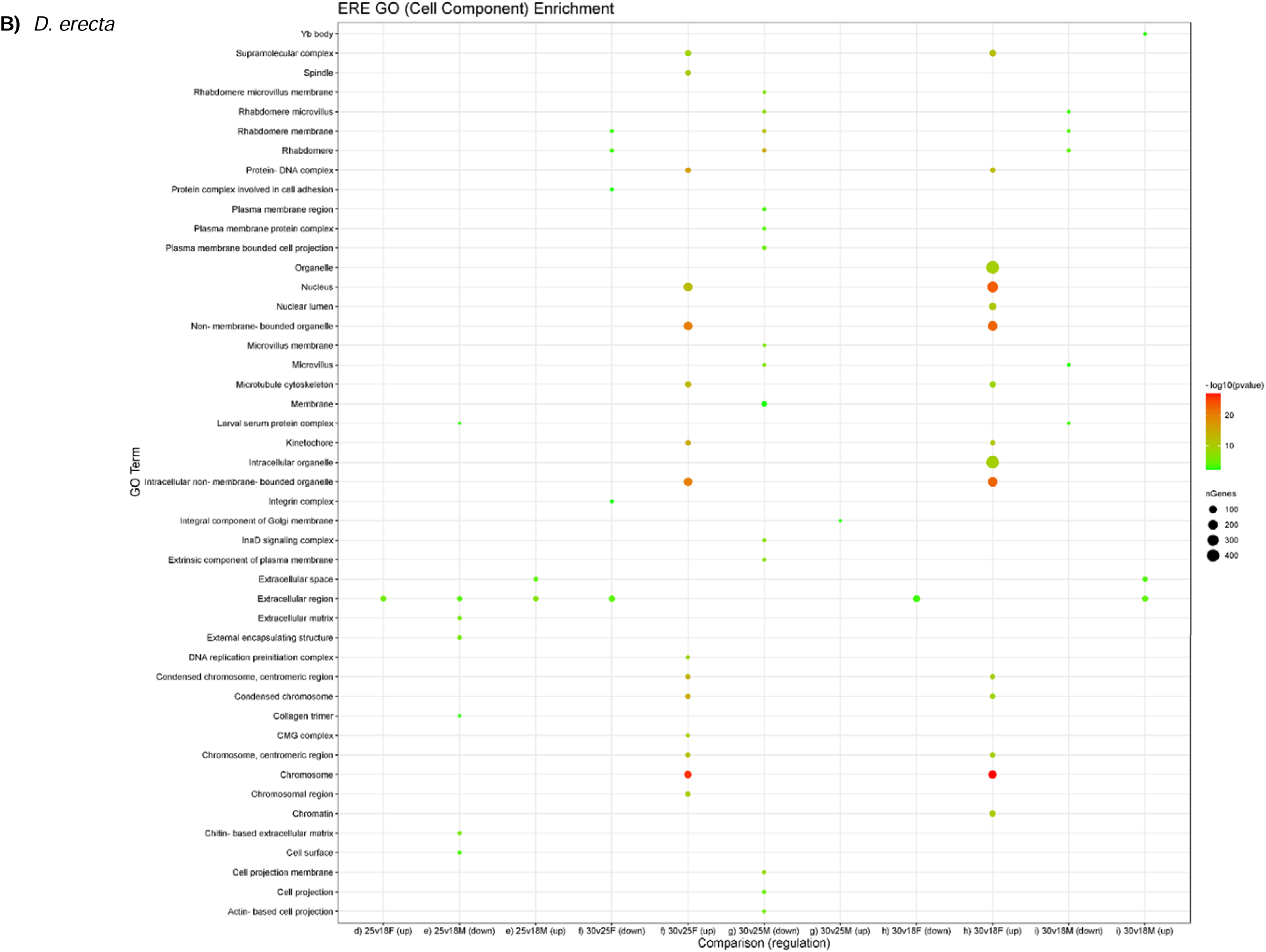
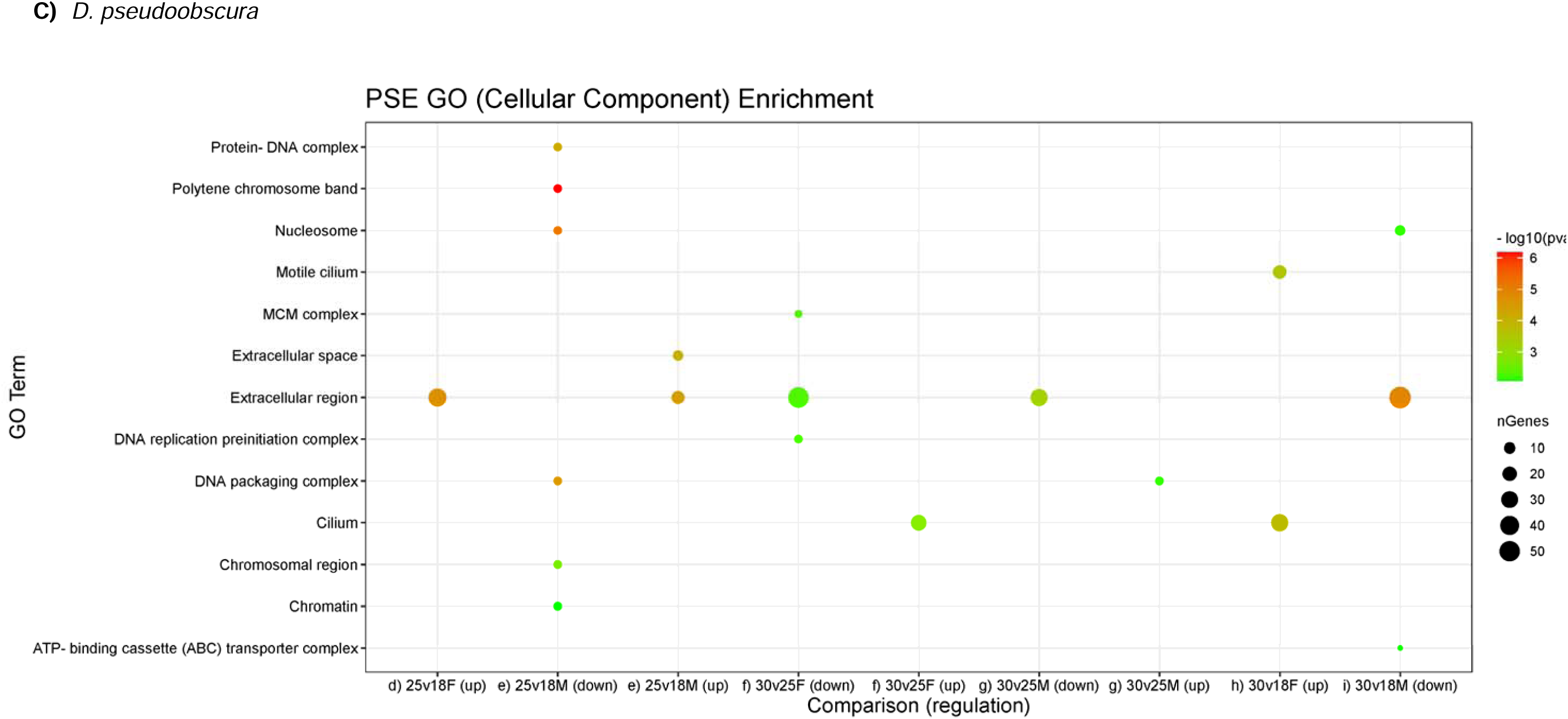
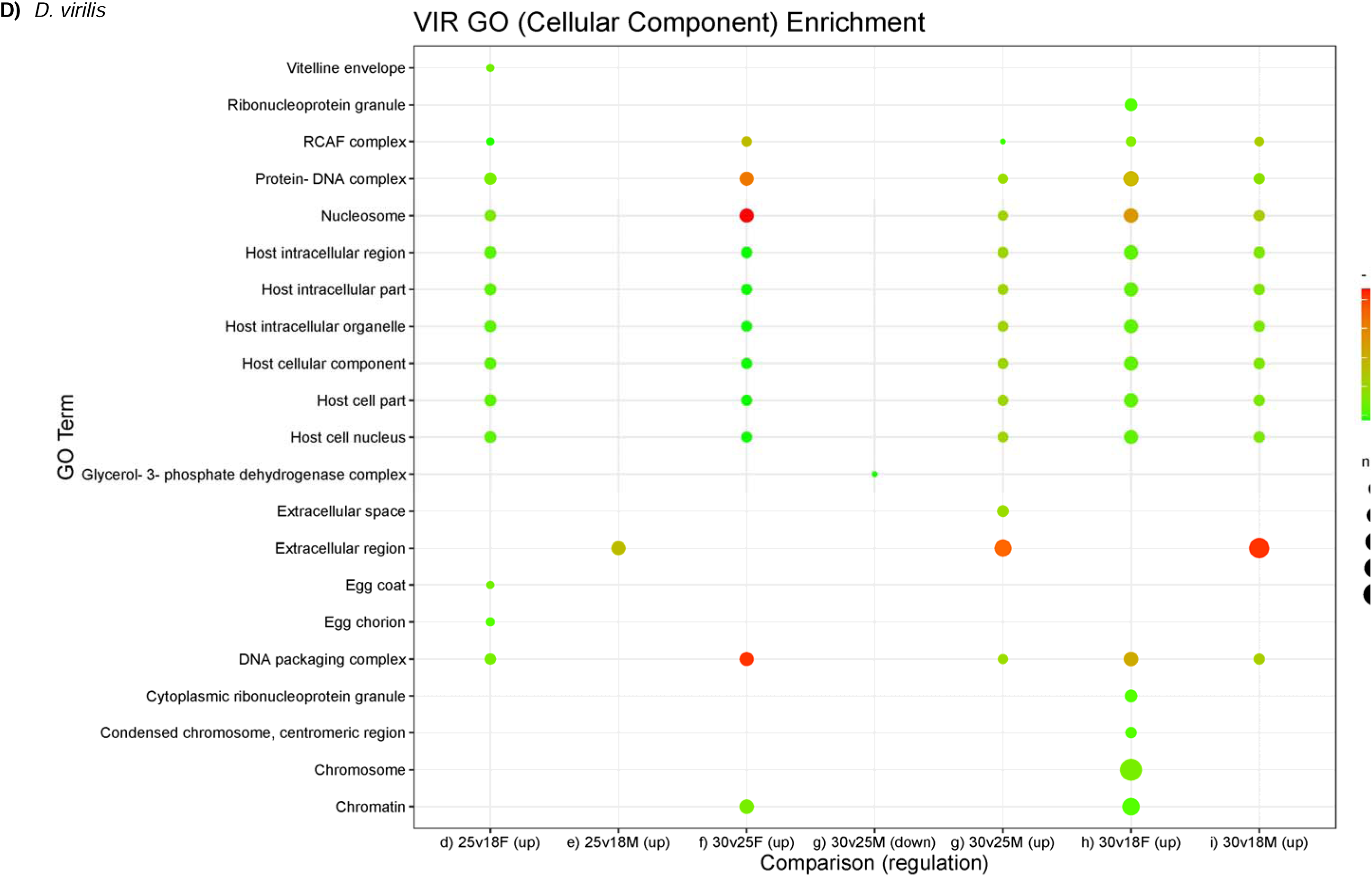
GO (Cellular Component) Enrichment for differential expressed genes under different temperatures

**Fig. S11.**
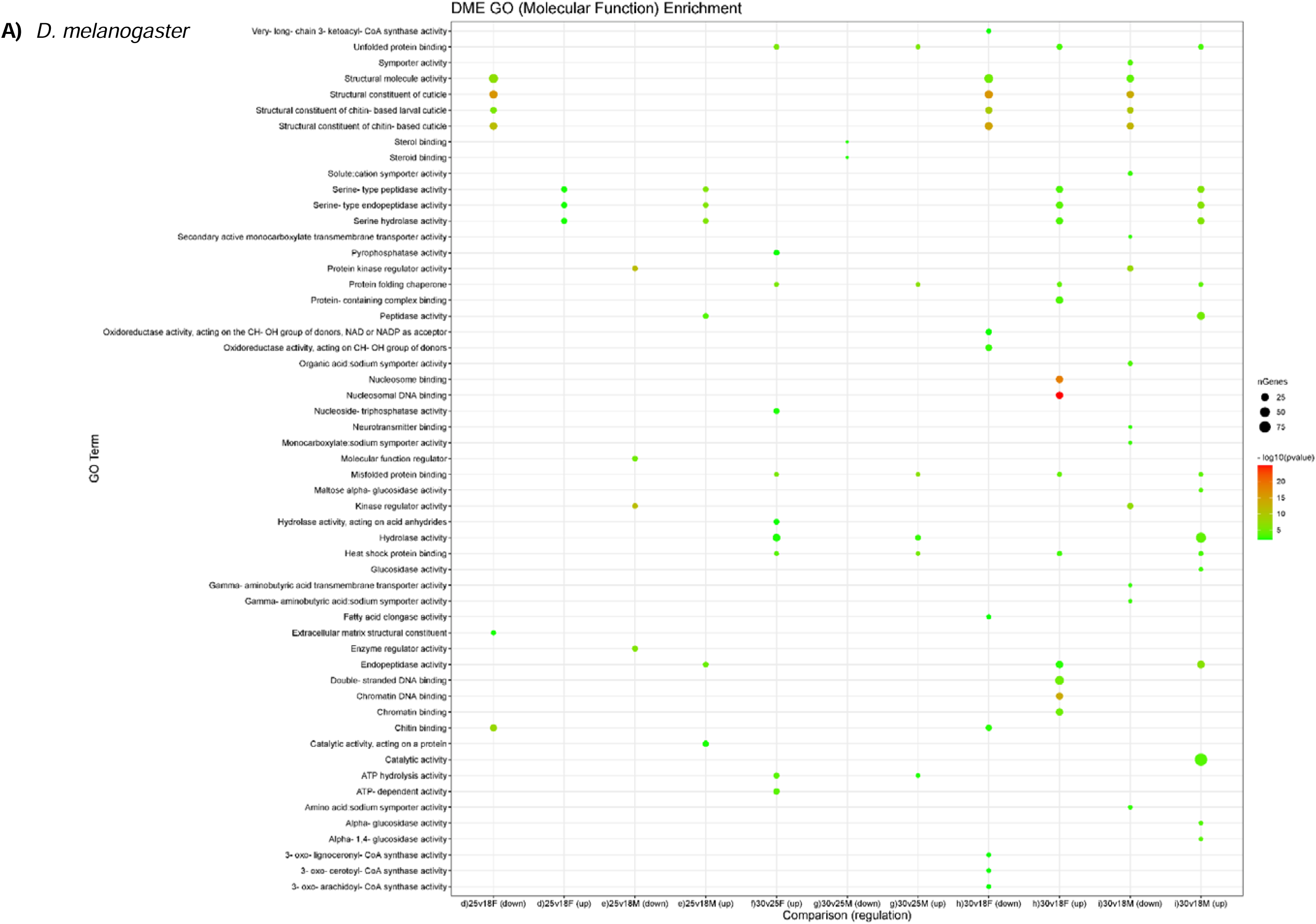
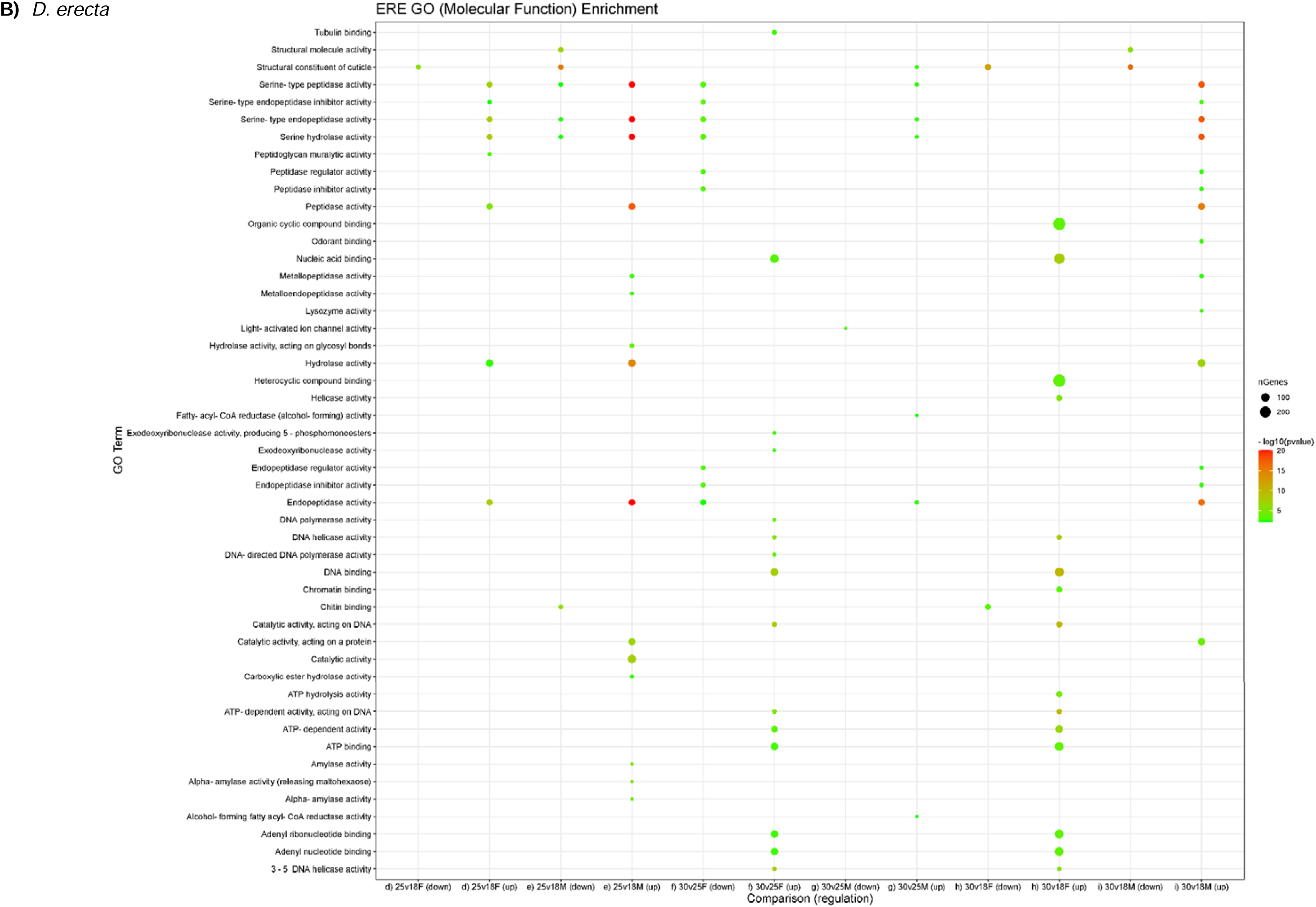
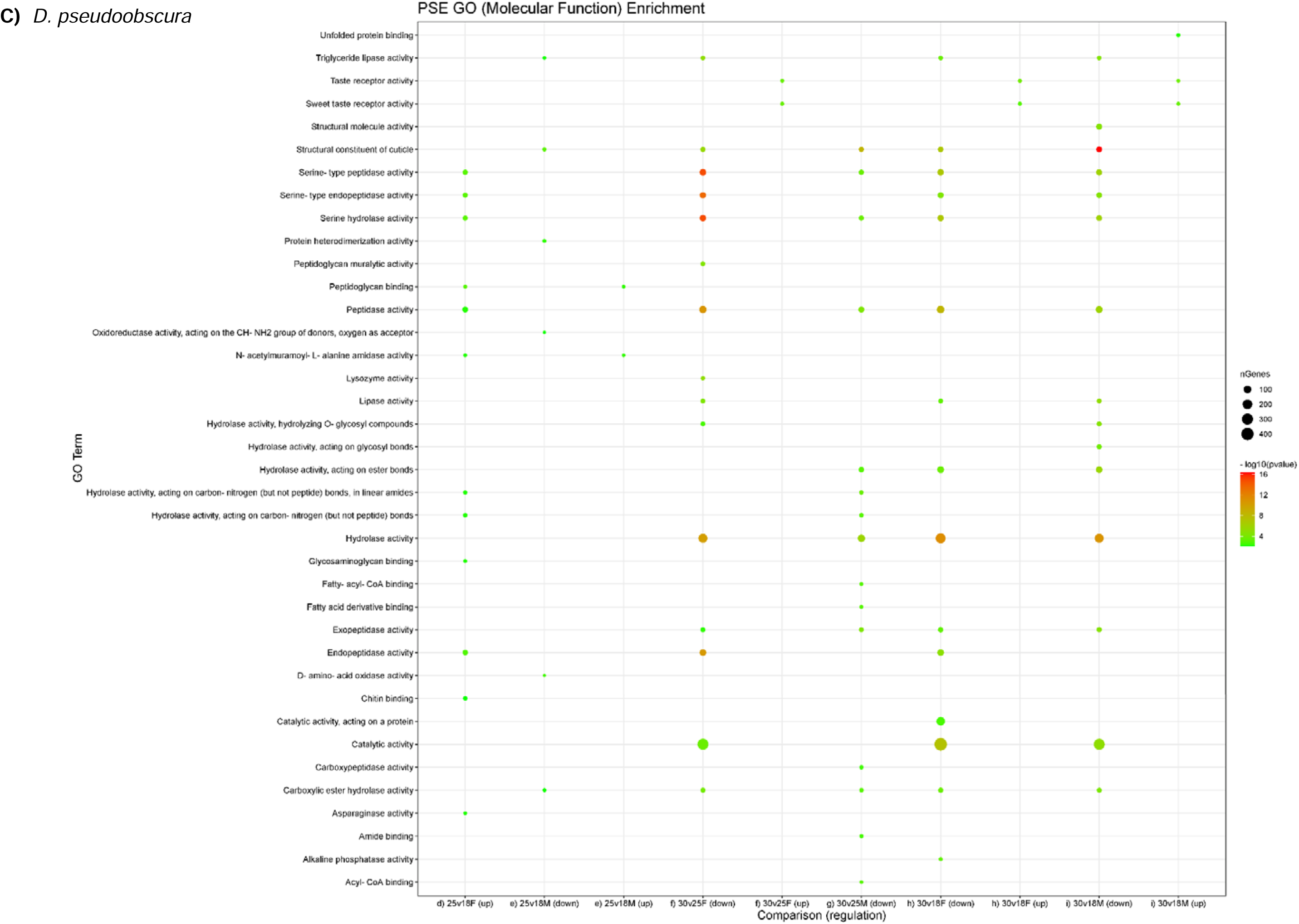
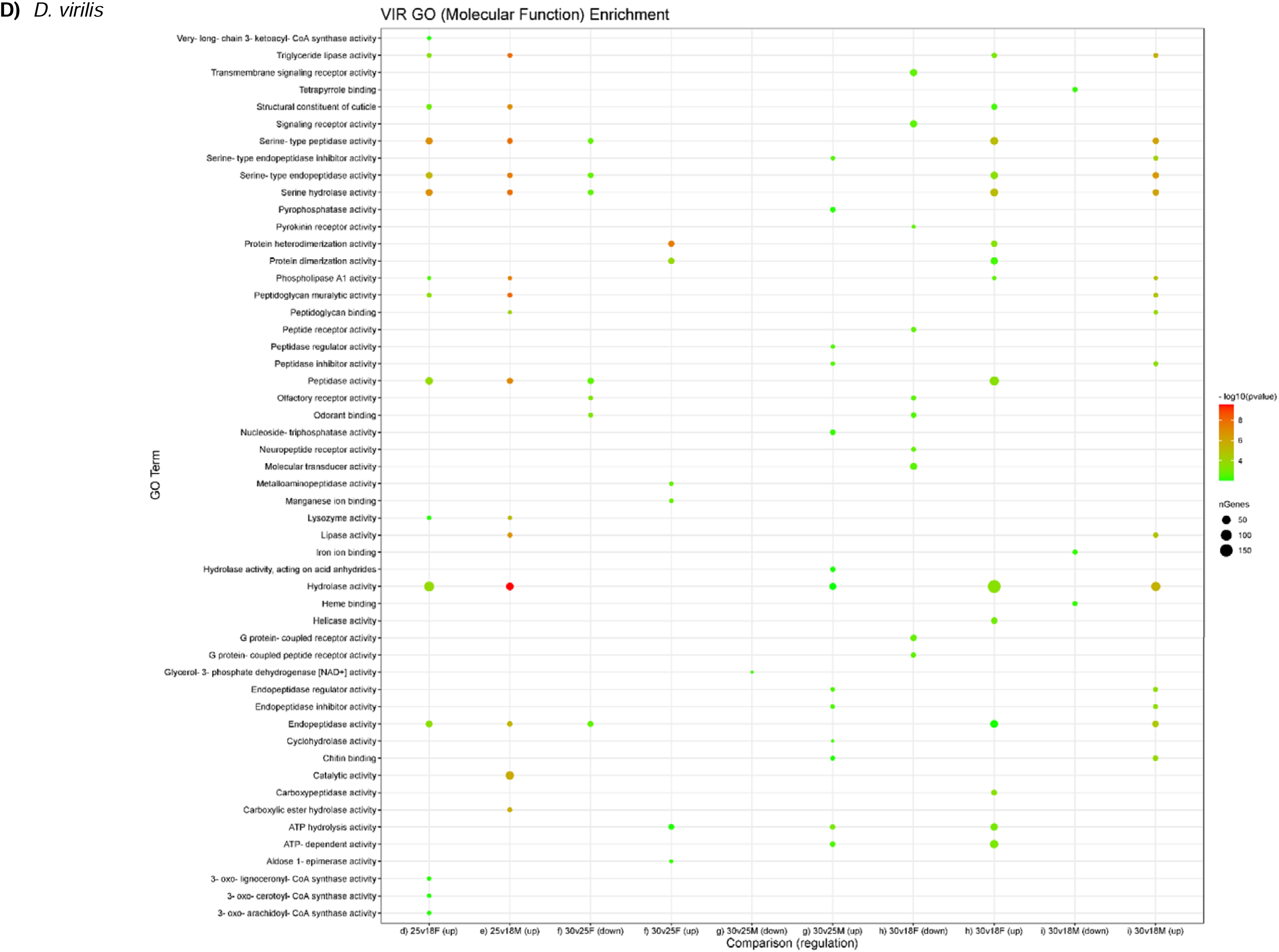
GO (Molecular Functions) Enrichment for differential expressed genes under different temperatures

**Fig. S12.**
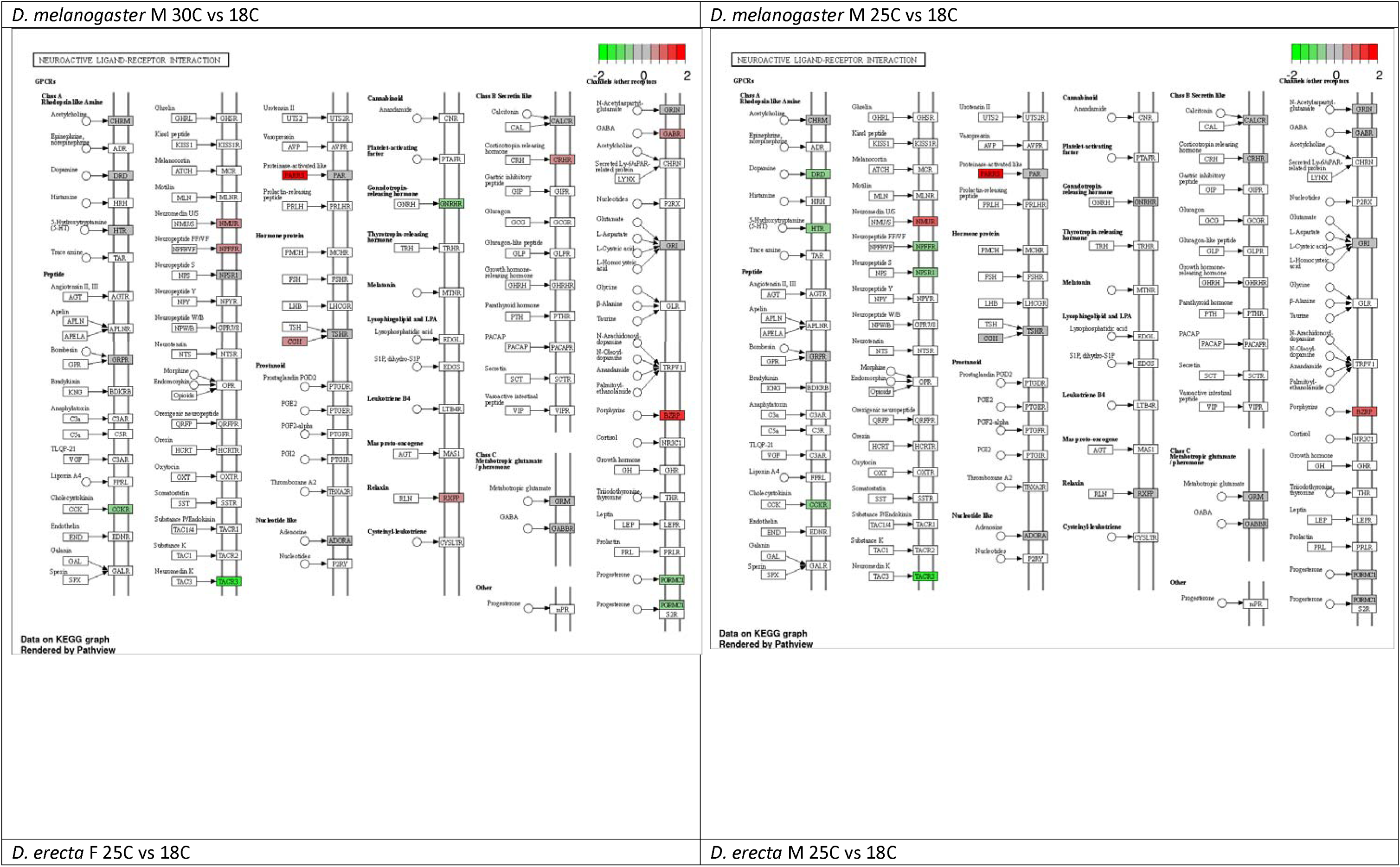
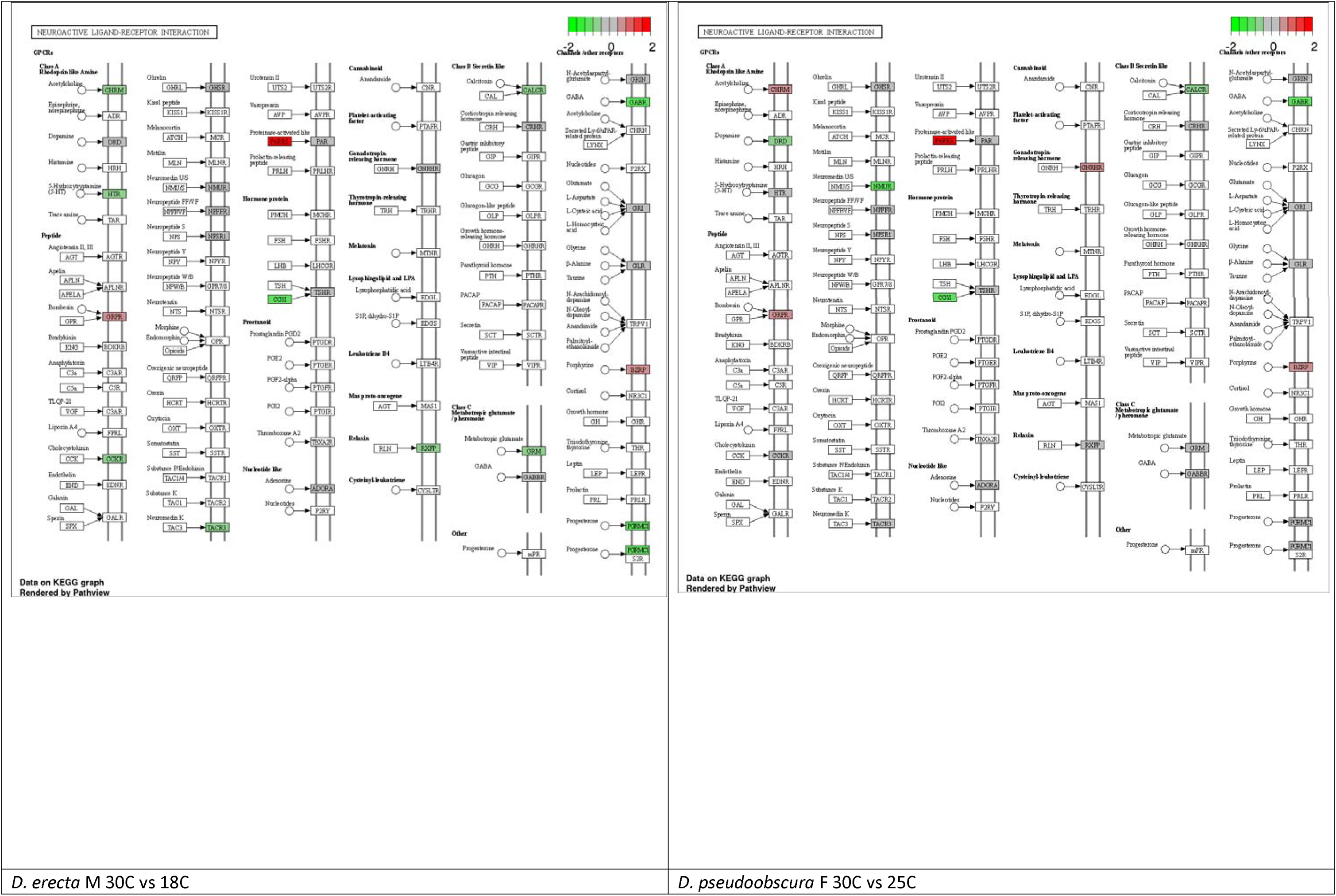
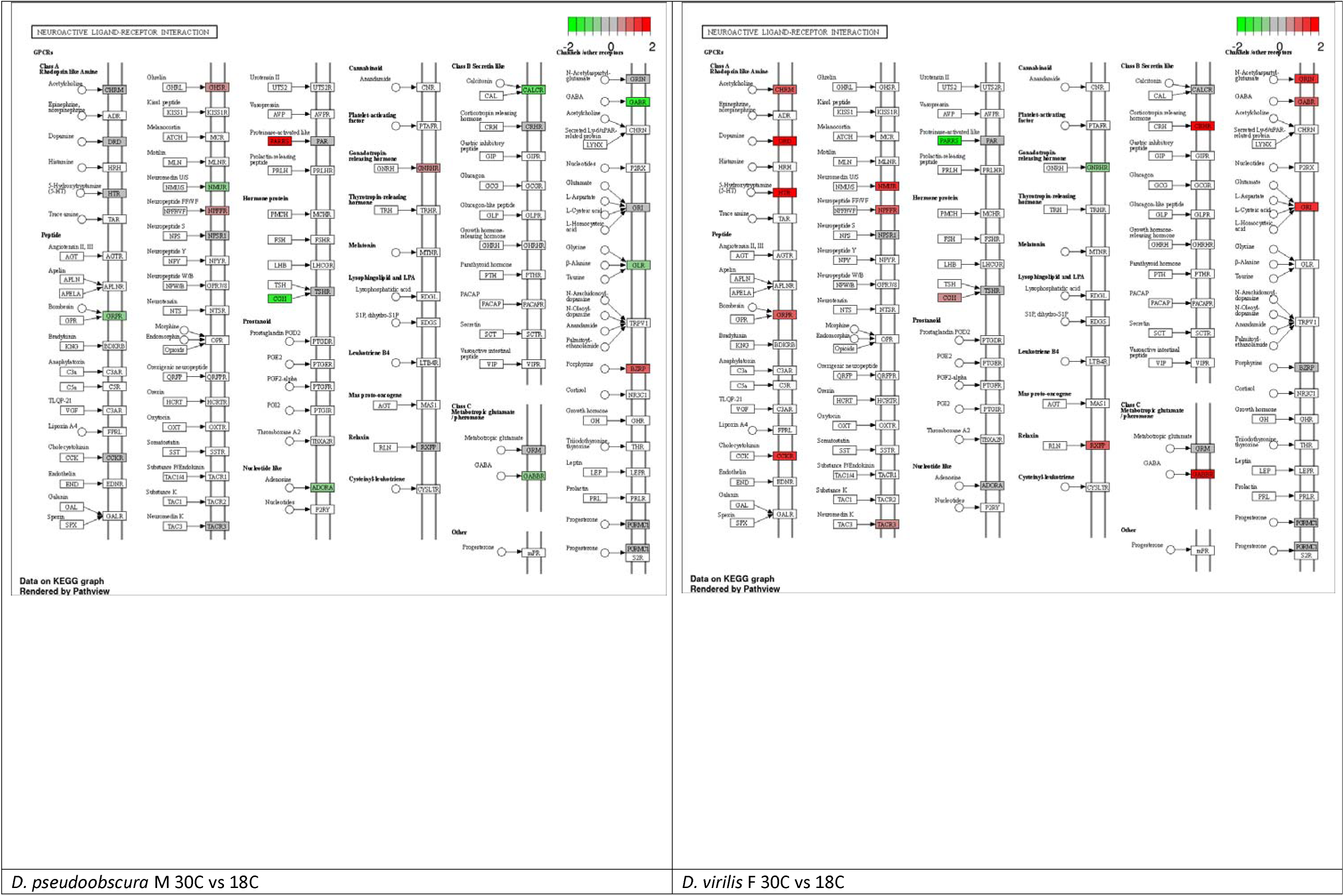
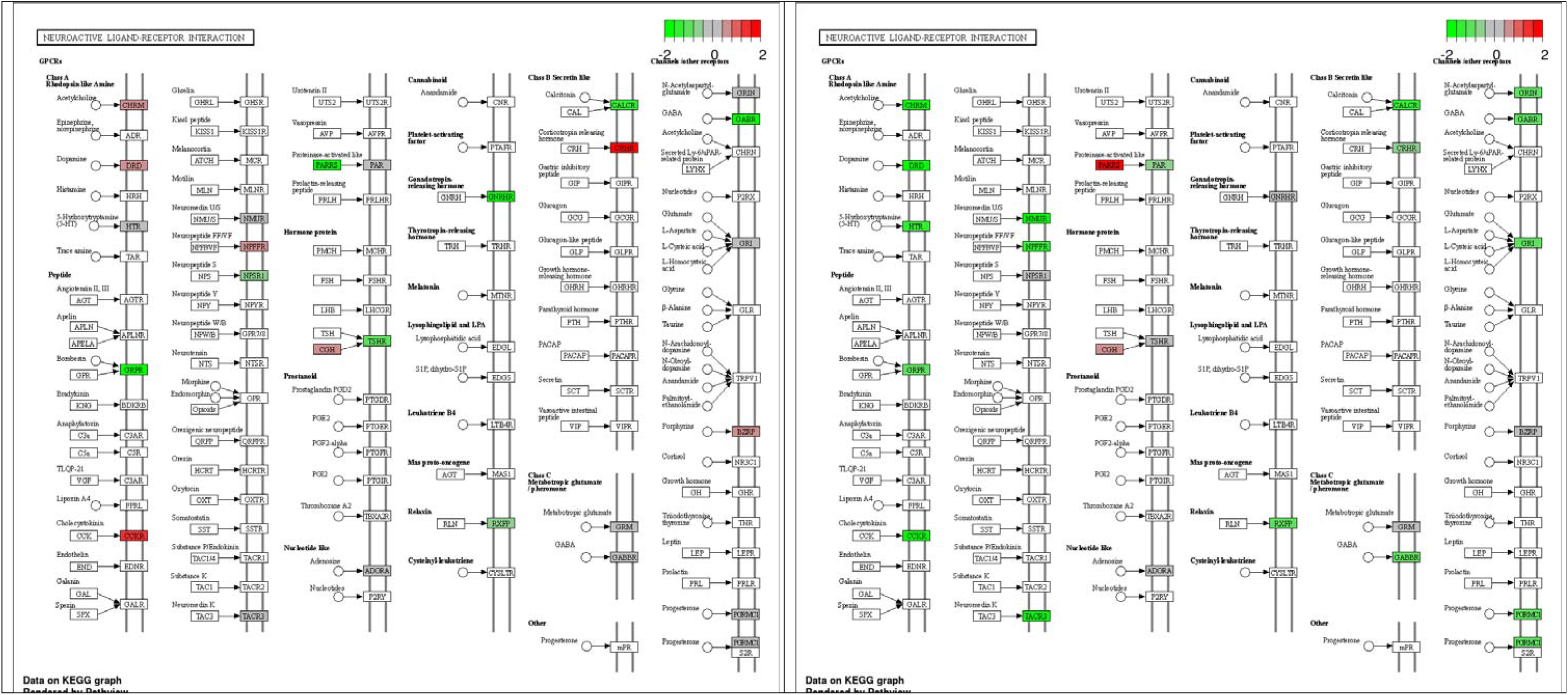
Regulation of genes in Neuroactive Ligand-receptor interaction pathway across different species under increased temperature.

**Fig. S13.**
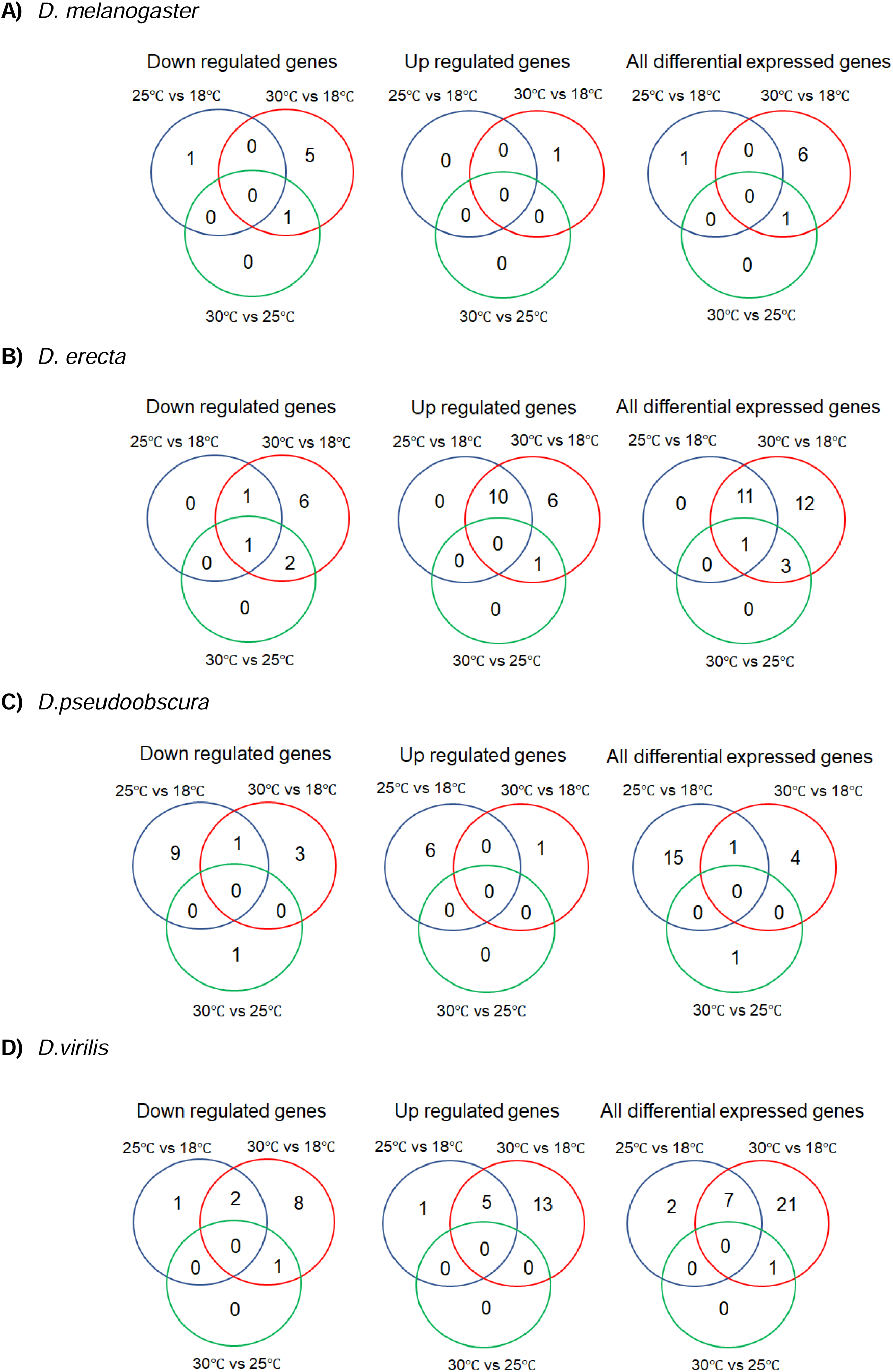
Venn diagrams showing differential expressed miRNAs in female under different temperature regimes

**Fig. S14.**
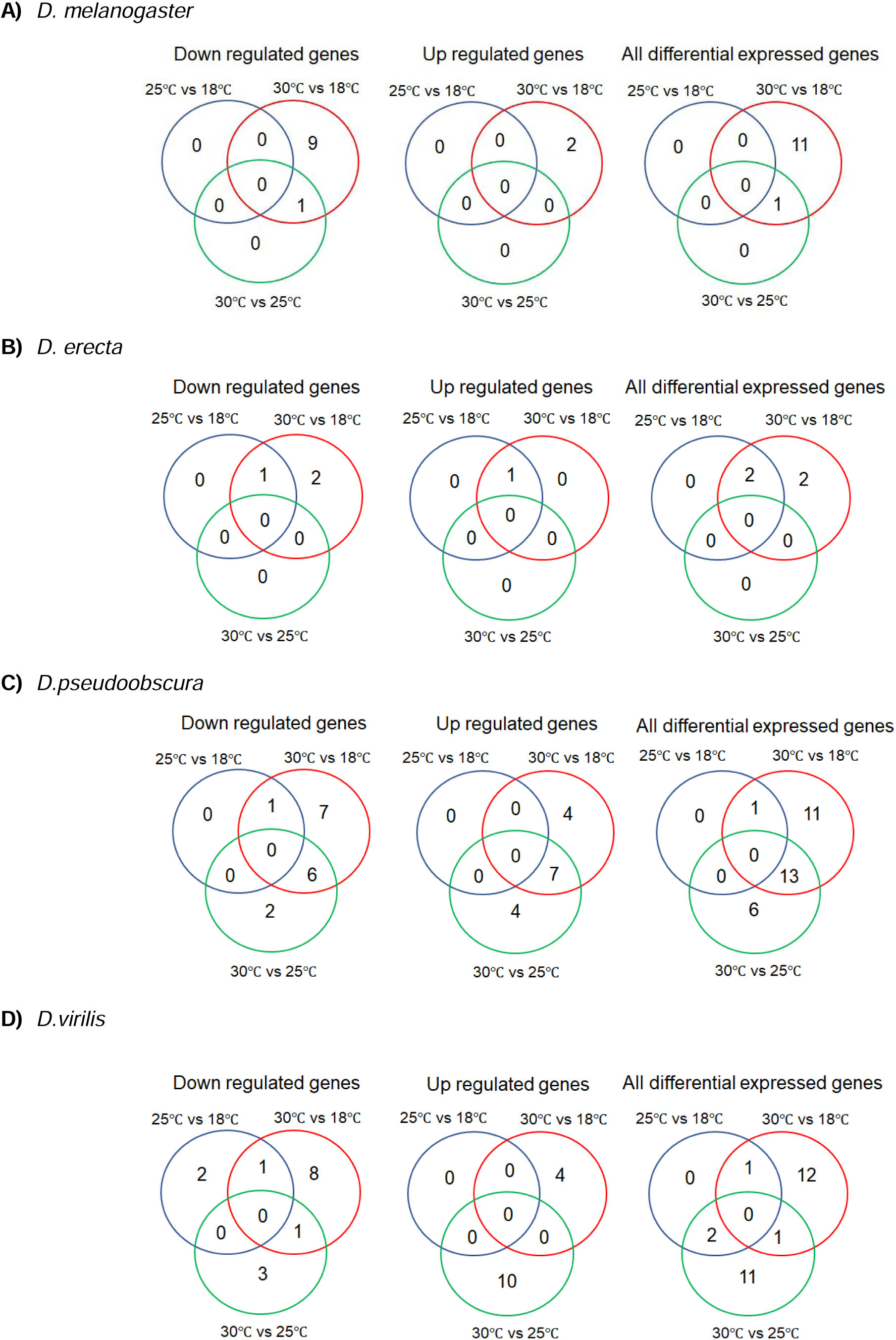
Venn diagrams showing differential expressed miRNAs in male under different temperature regimes

**Fig. S15.**
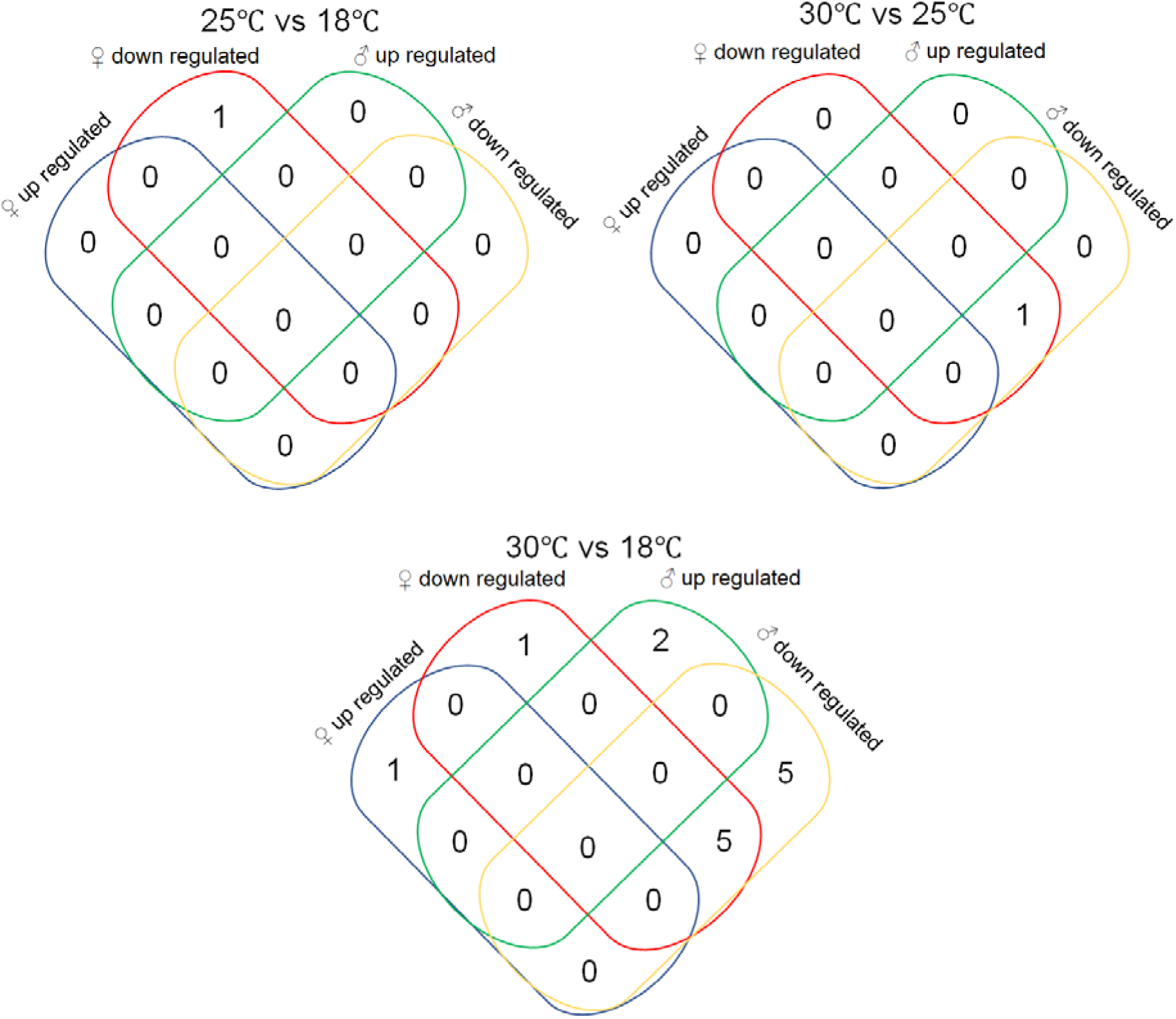
Venn diagrams showing different miRNAs regulation of male and female *D. melanogaster* under the same temperature regime

**Fig. S16.**
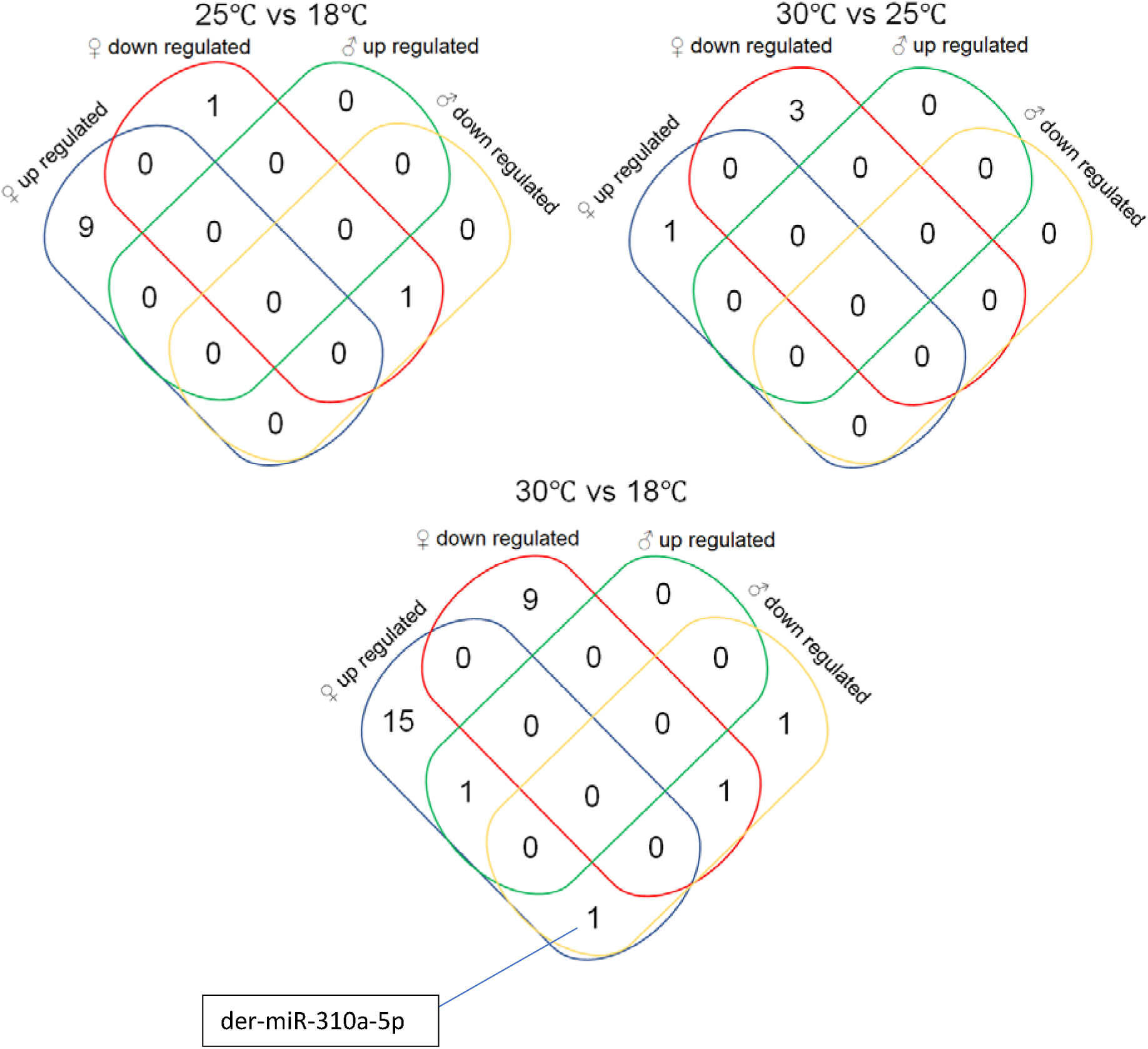
Venn diagrams showing different miRNAs regulation of male and female *D. erecta* under the same temperature regime

**Fig. S17.**
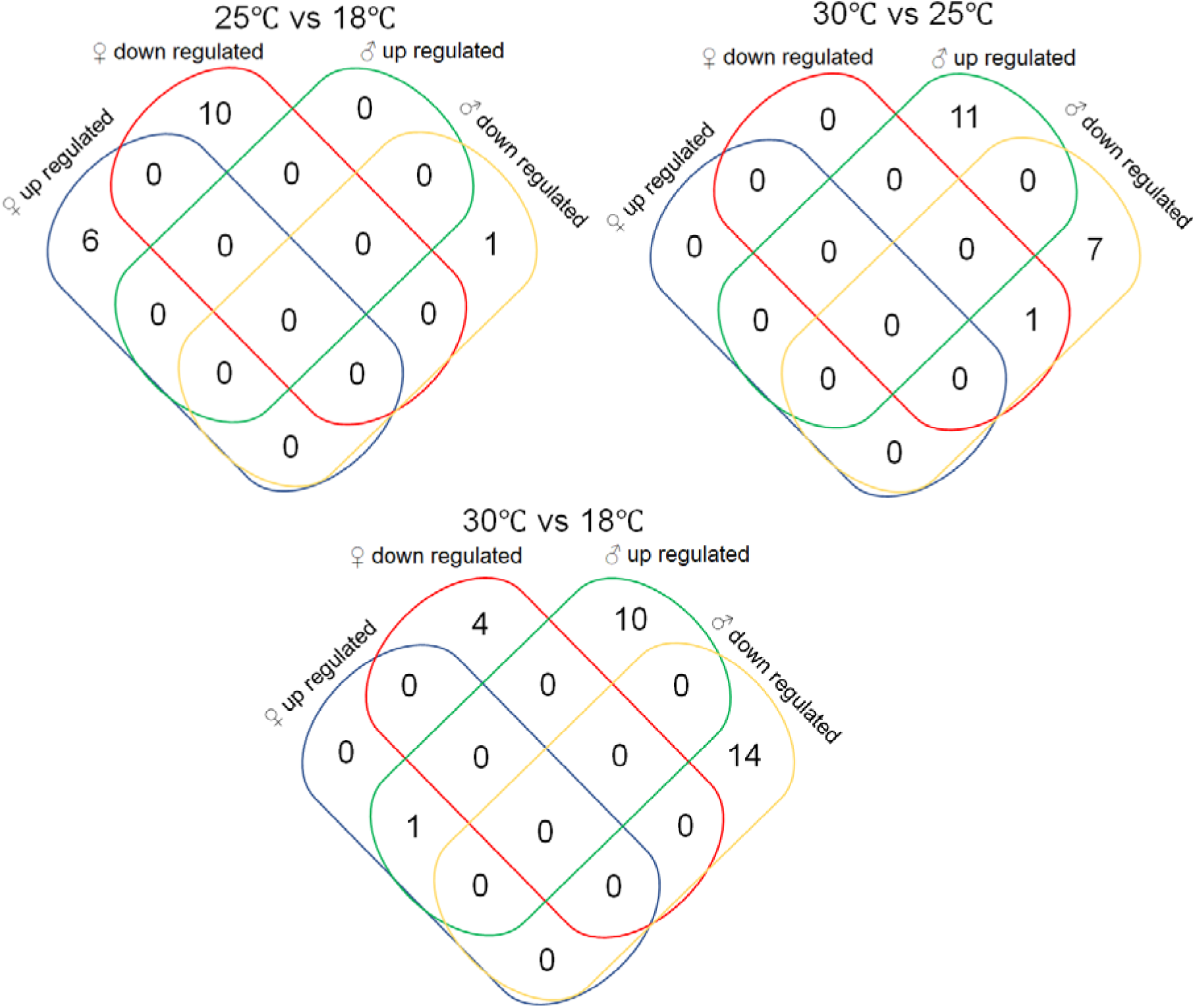
Venn diagrams showing different miRNAs regulation of male and female *D. pseudoobscura* under the same temperature regime

**Fig. S18.**
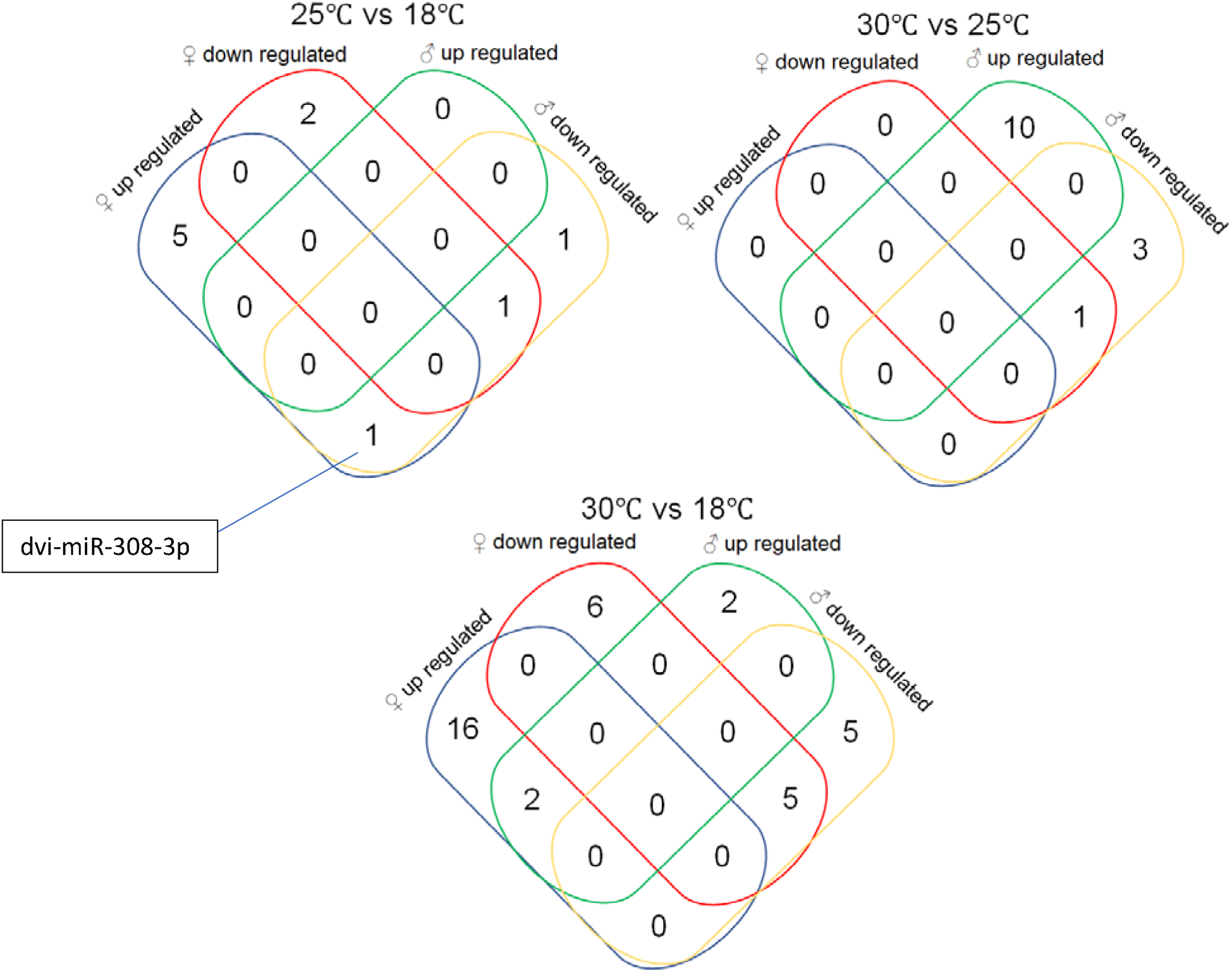
Venn diagrams showing different miRNAs regulation of male and female *D. virilis* under the same temperature regime

**Fig. S19.**
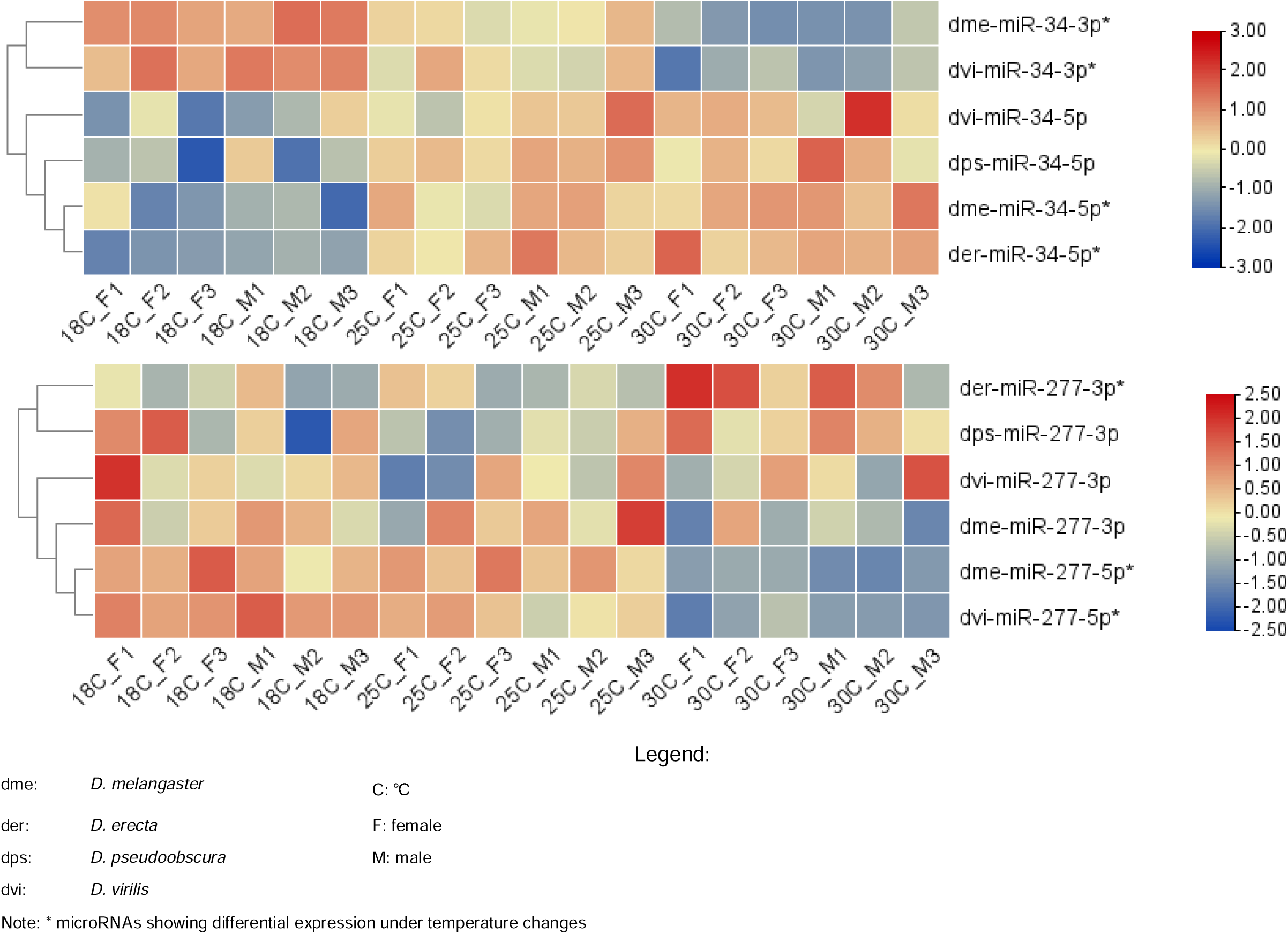
Heatmaps showing expression of miR-277 and miR-34 of all 4 drosophila species

